# Assessment of transparency indicators across the biomedical literature: how open is open?

**DOI:** 10.1101/2020.10.30.361618

**Authors:** Stylianos Serghiou, Despina G. Contopoulos-Ioannidis, Kevin W. Boyack, Nico Riedel, Joshua D. Wallach, John P. A. Ioannidis

## Abstract

Recent concerns about the reproducibility of science have led to several calls for more open and transparent research practices and for the monitoring of potential improvements over time. However, with tens of thousands of new biomedical articles published per week, manually mapping and monitoring changes in transparency is unrealistic. We present an open-source, automated approach to identify five indicators of transparency (data sharing, code sharing, conflicts of interest disclosures, funding disclosures and protocol registration) and apply it across the entire open access biomedical literature of 2.75 million articles on PubMed Central. Our results indicate remarkable improvements in some (e.g. conflict of interest disclosures, funding disclosures), but not other (e.g. protocol registration, code sharing) areas of transparency over time, and map transparency across fields of science, countries, journals and publishers. This work has enabled the creation of a large, integrated, and openly available database to expedite further efforts to monitor, understand and promote transparency and reproducibility in science.

## Introduction

Research reproducibility [1] is a fundamental tenet of science, yet recent reports suggest that reproducibility of published findings should not be taken for granted [2]. Both theoretical expectations [3] and empirical evidence [4–8] suggest that most published results may be either non-reproducible or inflated. Cardinal amongst recommendations for more credible and efficient scientific investigation [9–18] is the need for transparent, or open science [16,19].

Our group previously evaluated 441 randomly selected biomedical articles from 2000-2014 [20], illustrating that across several previously suggested indicators of transparency [21], only 136 (30.8%) articles had a conflict of interest disclosure (about the presence or absence of conflict of interest) and 213 (48.3%) had a funding disclosure (about presence or absence of funding); no article made its data or code openly available. A follow-up study of 149 articles published between 2015-2017 noted substantial improvements over time, with 97 (65.1%) sharing a conflict of interest disclosure, 103 (69.1%) sharing a funding disclosure, and 19 (12.8%) sharing at least some of their data; still, no code sharing was identified [22]. While our previous studies suggest low, but slowly improving levels of transparency and reproducibility, they are limited in that they only capture a small random subset of the biomedical literature and require laborious manual screening and data abstraction. A larger, high-throughput evaluation would be necessary to keep up-to-date with the pace of expanding literature and understand the distribution of transparency across more granular categories, such as time or fields of science.

Currently available tools can identify certain indicators of transparency, but they cannot be used to map and monitor these indicators across the published biomedical literature, their code is not openly available, their true performance is unknown or they are paid services [23–26]. A recent publication utilized methods of machine learning to identify, amongst others, conflicts of interest and funding disclosures, but it can only process acknowledgements [27]. Our work aims to expand our assessment of multiple indicators of transparency across the entire open biomedical literature, by developing and using tools and datasets that we hereby make freely available. These indicators are: Data sharing, Code sharing, Conflict of interest (COI) disclosures, Funding disclosures, and open Protocol registration.

## Results

### Manual assessment of transparency and reproducibility across 499 PubMed articles (2015-2018)

We started with a manual assessment of a random sample of 499 English language articles from PubMed published in recent years (2015-2018) (Table 1). COI disclosures and Funding disclosures were assessed in all 499 articles, whereas alternative indicators of transparency and reproducibility were only assessed in the more relevant subset of 349 articles with empirical data (henceforth referred to as “research articles”). This work expands on a previous in-depth assessment of a random sample of PubMed articles [22], as more recent literature is deemed likely to have the highest rates of transparency and yield additional data for training automated algorithms focusing on current practices.

**Table 1.** Characteristics of 499 randomly selected English language articles from PubMed between 2015-2018.

|  |  | Number (%) |
| --- | --- | --- |
| **Total PubMed articles** |  | **499** |
| **Year of publication** | **2015** | 121 (24%) |
|  | **2016** | 121 (24%) |
|  | **2017** | 143 (29%) |
|  | **2018** | 114 (23%) |
| **On PubMed Central** |  | 197 (40%) |
| **Area** | **Pre-clinical research** | 331 (66%) |
|  | **Clinical research** | 18 (4%) |
| **Field of science** | **Medicine** | 274 (55%) |
|  | **Health services** | 69 (14%) |
|  | **Brain research** | 46 (9%) |
|  | **Biology** | 46 (9%) |
|  | **Other (n = 5)** | 62 (12%) |
| **Characteristics** |  | **Median (IQR)** |
|  | **Authors per article** | 4 (2-7) |
|  | **Affiliations per article** | 5 (2-8) |
|  | **Citation count** | 2 (0-5) |
|  | **Journal Impact Factor** | 2.8 (2.0-4.3) |
|  | **Altmetric Attention Score** | 0.5 (0.0-2.7) |

*Missing values: Field of science (2, 1%), Affiliations per article (25, 5%), Citation count (6, 1%) and Journal Impact Factor (76, 15%). IQR = interquartile range. Field of science was taken from SciTech, Citation count from Crossref (30 May, 2019), Journal Impact Factor of 2018 from Web of Science and Altmetric Attention Score from Altmetric (30 May, 2019).*

Out of all 499 articles, 341 (68.3%) had a COI disclosure and 352 (70.5%) had a Funding disclosure. Of 349 research articles, 68 (19.5%) had a Data sharing statement, 5 (1.4%) had a Code sharing statement, 246 (70.5%) had a COI disclosure, 284 (81.4%) had a Funding disclosure, 22 (6.3%) had an openly registered Protocol, 175 (50.1%) made a statement of Novelty (e.g. “report for the first time”), and 33 (9.5%) included a Replication component in their research (e.g. validating previously published experiments, running a similar clinical trial in a different population, etc.) (Fig 1A). Most articles with transparency statements or disclosures claimed no COIs (88.6% [218/246] of research articles with COI disclosures), and a substantial portion mentioned use of public funds (36.3% [103/284] of research articles with Funding disclosures), availability of data upon request (25.0% [17/68] of research articles with Data sharing), and registration on ClinicalTrials.gov (50.0% [11/22] of research articles with Protocol registration) (S1 Fig). In eight (11.8%) of 68 articles with a Data sharing statement claiming that all data were available in the text, no raw data were found.

**Fig 1.**
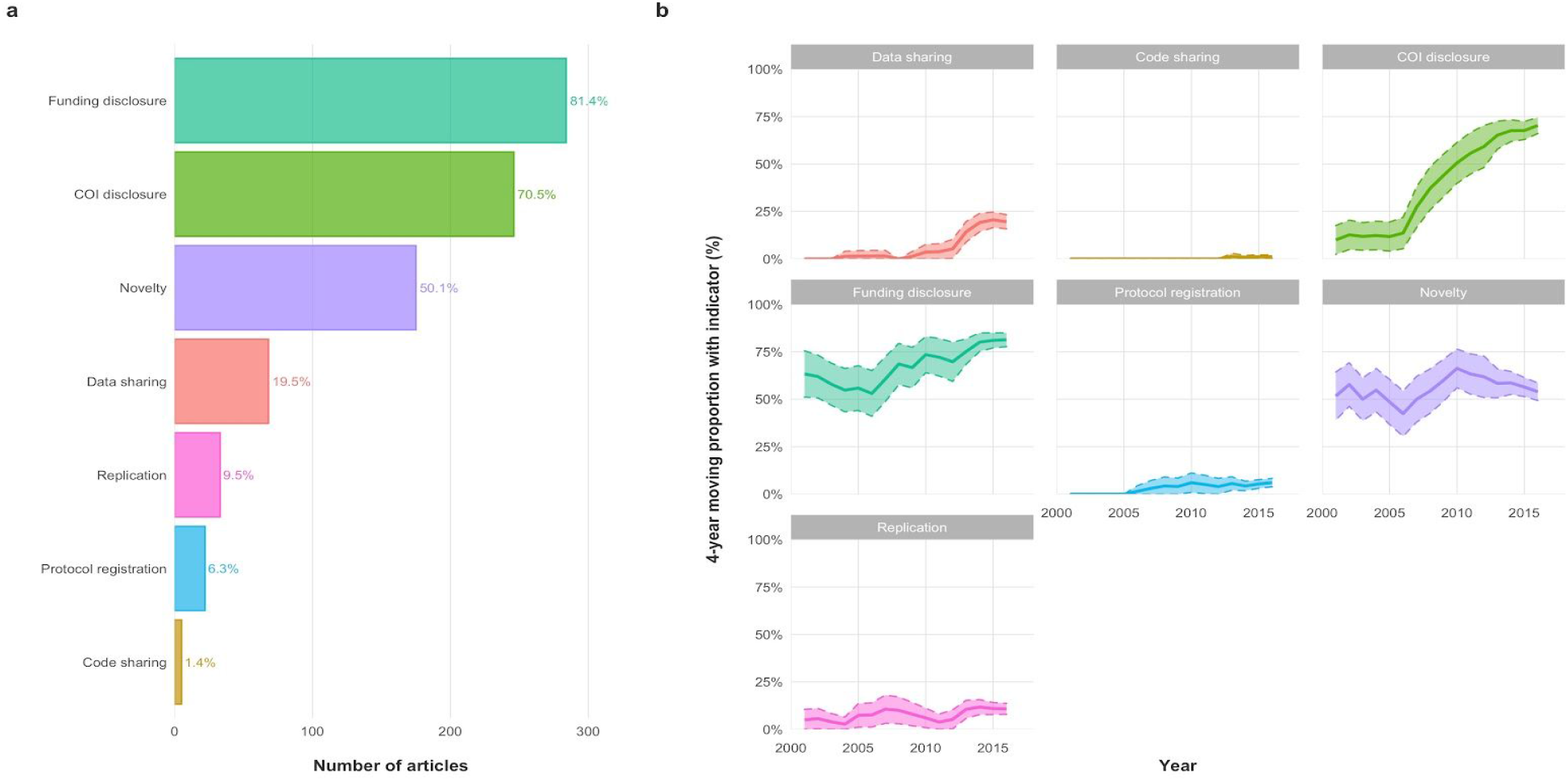
Indicators of transparency and reproducibility across publications and time. (A) Indicators of transparency and reproducibility across 349 research articles (2015-2018). Most publications included Conflict of interest (COI) or Funding disclosures, but few mentioned Data, Code or Protocol sharing. Similarly, most claimed Novelty but few mentioned a Replication component. (B) Indicators of transparency and reproducibility across time. Proportions are displayed as a 4-year centered moving average. These graphs combined data from this study on 349 research articles with data from two previous studies on another 590 PubMed articles [20,22]. The shaded region indicates the 95% Confidence Interval (CI). The most notable change is that of COI disclosures, the reporting of which increased from 12% in 2000 to 76% in 2018.

Using information found in the PubMed records of these articles alone (as opposed to screening the full text), would have missed most records with indicators of transparency (S1 Table). Abstracts are very limited in scope and PubMed records are not tailored to systematically capture these indicators, perhaps with the exception of funding. Still, PubMed records gave information on funding for merely 35% of articles, while funding disclosures were in fact present in 71% of them.

By utilizing data from previous studies [20,22], we observed a marked increase in the proportion of publications reporting COI disclosures (Fig 1B, S2 Table). An increase is also seen in the availability of Funding disclosures and Data sharing statements, even though uncertainty is too large to claim trends. For Code sharing and Protocol registration, the numbers have remained low. No apparent trends were observed in statements of Novelty and Replication.

In our sample, reporting of COI disclosures, Funding disclosures, Data sharing and Code sharing were significantly more frequently seen in articles available on PubMed Central (PMC) (S3 Table). Pre-clinical research articles reported Funding disclosures more often than clinical research articles (91% vs 70%) and clinical trials reported Protocol registration substantially more frequently than the rest (67% vs 3%).

### Automated assessment of transparency: Development and validation across 6017 PMC articles (2015-2019)

We proceeded to test the performance of algorithms automating the process of identifying indicators of transparency. Three were developed from scratch (COI disclosure, Funding disclosure and Protocol registration) and two were adopted from an already existing library [28] (Data and Code sharing) and customised to enhance efficiency (see Materials and Methods). Note that, even though in the manual assessment, Data sharing and Code sharing indicators capture any statement about data or code availability, in the automated assessment we only capture newly generated open raw data or code. In a random unseen sample of 6017 PMC records from 2015-2019, these algorithms predicted Data sharing in 764 (13%) records, Code sharing in 117 (2%), COI disclosure in 4,792 (80%), Funding disclosure in 5022 (84%), and Protocol registration in 261 (4%); 61 (1%) claimed both Data and Code sharing. Examples for the presence of any of those indicators can be found in the Supporting Information (S2 Fig).

In the tested samples of 6017 PMC articles, the accuracy, positive predictive value (PPV) and negative predictive value (NPV) of all 5 algorithms was ≥88% (Fig 2). Even though all algorithms were highly specific (>98%), the data and code sharing algorithms were not as sensitive as the rest (76% and 59% respectively vs >95% for all other indicators). This was in part because algorithms did not identify raw data made available as supplements, or did not recognise less popular code repositories (BitBucket and Open Science Framework). The estimated sensitivity of the code sharing algorithm was particularly driven by a single study that provided its code as a supplement, but was falsely labelled negative (see Code Sharing in S2 Text). As such, the algorithm made 1 mistake in 88 manually assessed research articles with no Data or Code sharing. However, with the vast majority of articles not sharing data or code (5197/6017), this 1 article was dramatically overweighted. This is reflected by the large confidence interval (34-94%), which includes the estimated sensitivity of 73% from a random sample of 800 PMC research articles of 2018 calculated by the original authors of this algorithm [29].

**Fig 2.**
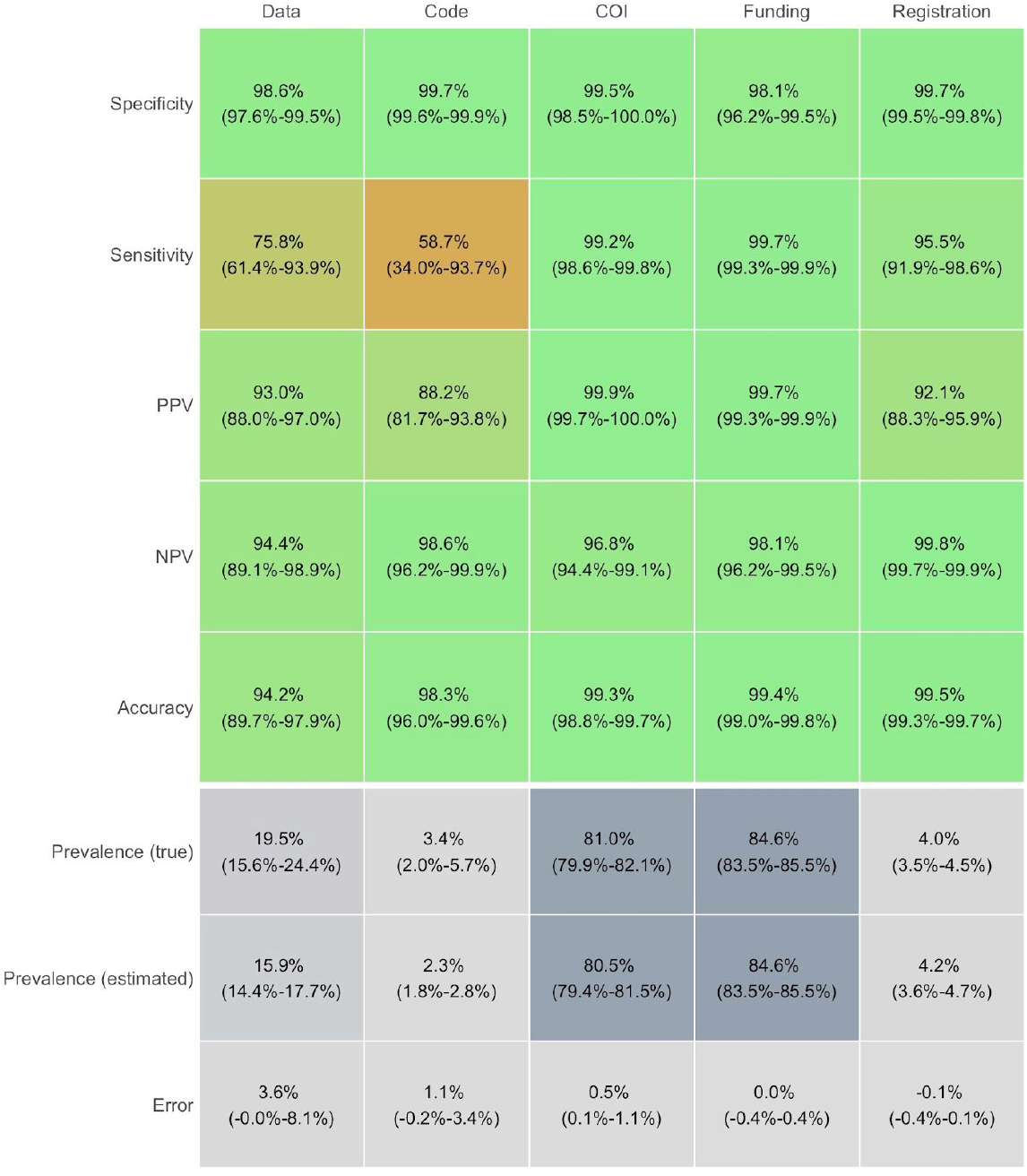
Validation of algorithms for Data sharing, Code sharing, COI disclosure, Funding disclosure and Protocol registration in 6017 PMC articles from 2015-2019. The displayed performance was assessed in subsamples from 6017 PMC articles: 189 research articles for Data sharing (100 positive, 89 negative), 291 research articles for Code sharing (110 positive, 181 negative), 325 articles for COI disclosure (100 positive, 225 negative), 326 for Funding disclosure (100 positive, 226 negative) and 308 for Protocol availability (161 positive, 147 negative). All algorithms displayed high accuracy (>94%) and low error in prevalence estimation (≤3.6%) compared to manual assessment. PPV = Positive Predictive Value (precision); NPV = Negative Predictive Value; Prevalence (true) = manual estimate of proportion of articles with indicator; Prevalence (estimated) = automated estimate of proportion of articles with indicator; Error = difference between true and estimated prevalence.

The difference between manually-adjudicated indicator prevalence (true) versus machine-adjudicated prevalence (estimated) was ≤1.1% for all algorithms other than for Data sharing, for which it was 3.6%. By design, the COI disclosure, Funding disclosure, and Protocol registration algorithms had 100% sensitivity and specificity in the training set of 499 articles (2015-2018); the Data and Code sharing algorithms had 76% sensitivity and 98% specificity in a training set of 868 random PubMed articles from 2015-2017 [28]. Detailed assessment of each algorithm can be found in the Supporting Information (S2 Text).

### Automated assessment of transparency: Transparency across 2.75 million PMCOA articles (1959-2020)

We then proceeded to test the entire PubMed Central Open Access (PMCOA) subset using these algorithms. We identified 2,751,484 records as of Feb 29, 2020, of which 2,751,420 were unique (Table 2). For each record we extracted 158 variables, of which 39 were meta-data (35 from PMC, 2 from OCC (Open Citation Collection), 2 from WOS (Web of Science), and 1 from SciTech) (S4 Table) and 119 were related to the indicators of transparency (5 for presence/absence of each indicator, 5 with extracted text for each indicator, and 93 indicating which aspects of text were found relevant for each indicator) (S5 Table). Of these, 2,285,193 (83.1%) had been labelled as research articles by OCC and 2,498,496 (90.8%) were published from 2000 onwards in a total of 10,570 different journals.

**Table 2.**
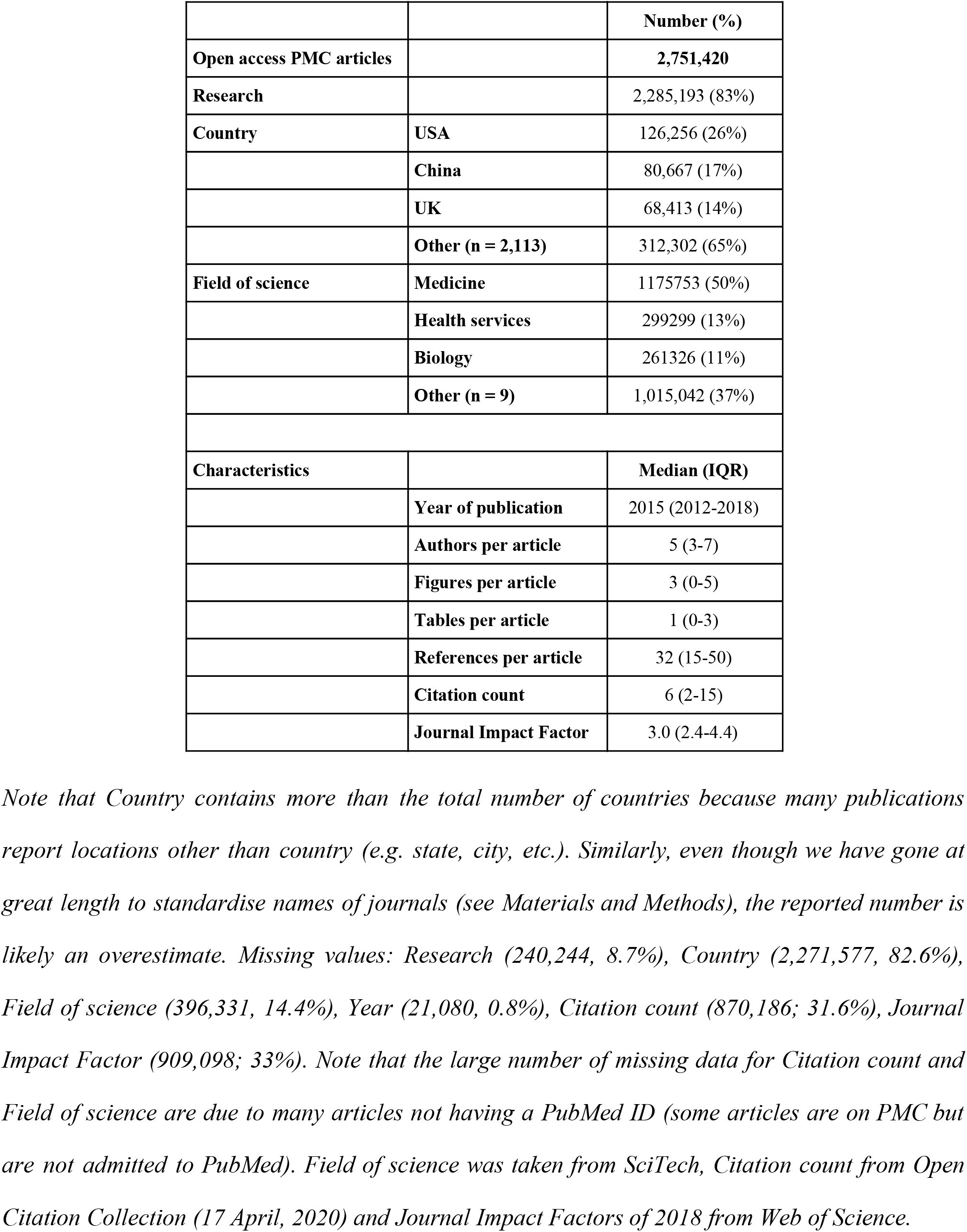
The distribution of meta-data across all 2,751,420 open access publications on PubMed Central (PMCOA). Note that Country contains more than the total number of countries because many publications report locations other than country (e.g. state, city, etc.). Similarly, even though we have gone at great length to standardise names of journals (see Materials and Methods), the reported number is likely an overestimate. Missing values: Research (240,244, 8.7%), Country (2,271,577, 82.6%), Field of science (396,331, 14.4%), Year (21,080, 0.8%), Citation count (870,186; 31.6%), Journal Impact Factor (909,098; 33%). Note that the large number of missing data for Citation count and Field of science are due to many articles not having a PubMed ID (some articles are on PMC but are not admitted to PubMed). Field of science was taken from SciTech, Citation count from Open Citation Collection (17 April, 2020) and Journal Impact Factors of 2018 from Web of Science.

|  |  | Number (%) |
| --- | --- | --- |
| **Open access PMC articles** |  | **2,751,420** |
| **Research** |  | 2,285,193 (83%) |
| **Country** | **USA** | 126,256 (26%) |
|  | **China** | 80,667 (17%) |
|  | **UK** | 68,413 (14%) |
|  | **Other (n = 2,113)** | 312,302 (65%) |
| **Field of science** | **Medicine** | 1175753 (50%) |
|  | **Health services** | 299299 (13%) |
|  | **Biology** | 261326 (11%) |
|  | **Other (n = 9)** | 1,015,042 (37%) |
| **Characteristics** |  | **Median (IQR)** |
|  | **Year of publication** | 2015 (2012-2018) |
|  | **Authors per article** | 5 (3-7) |
|  | **Figures per article** | 3 (0-5) |
|  | **Tables per article** | 1 (0-3) |
|  | **References per article** | 32 (15-50) |
|  | **Citation count** | 6 (2-15) |
|  | **Journal Impact Factor** | 3.0 (2.4-4.4) |

Out of 2,751,420 open access PMC records (1959-2020), our algorithms identified mentions of Data sharing in 243,783 (8.9%), Code sharing in 33,405 (1.2%), COI disclosure in 1,886,907 (68.6%), Funding disclosure in 1,858,022 (67.5%) and Protocol registration in 70,469 (2.6%) (Fig 3A, S6 Table). Adjusting for the discrepancies between predicted and true estimates seen in the validation set, we estimate that Data sharing information was mentioned in 14.5% (95% CI, 11.0-18.8%), Code sharing information in 2.5% (95% CI, 1.2-4.7%), COI disclosure information in 69.5% (95% CI, 69.0-70.1%), Funding disclosure information in 67.9% (67.6-68.3%), and Protocol registration information in 2.5% (95% CI, 2.5-2.6%). The majority of COI disclosures reported no conflicts of interest (1,531,018; 81.1%) and the majority of Funding disclosures reported receipt of funds (1,675,513; 90.2%). Associations between indicators and literature characteristics are reported in the Supporting Information (S3 Text and S7 Table). Note that unlike in our manual assessment, the numbers reported in this section refer to the entire literature on PMCOA, not merely research articles; all analyses were repeated in research articles alone with no meaningful changes.

**Fig 3.**
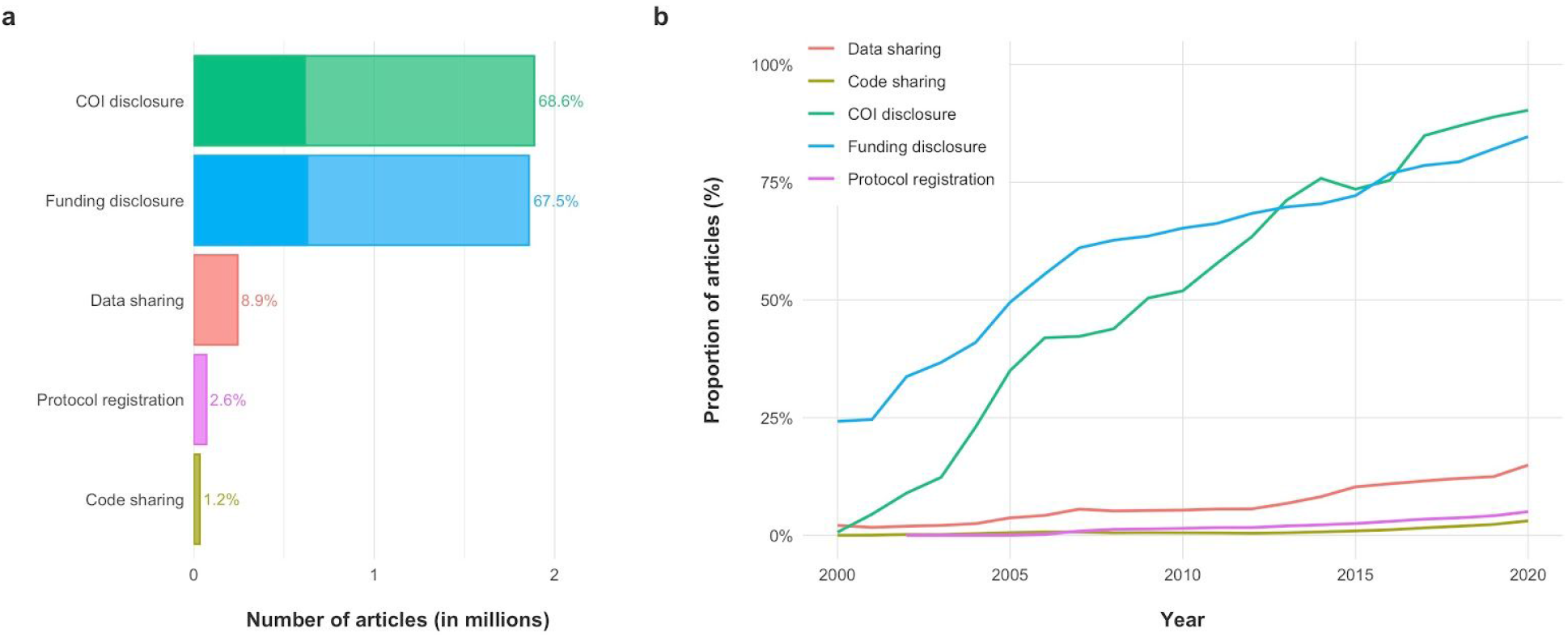
Indicators of transparency across the entire open biomedical literature on PMC (PMCOA) and time. (A) Most open biomedical articles report COI disclosures and Funding disclosures, but only a minority report any Data sharing, Code sharing, or open Protocol registration. Note that this figure displays the obtained results with no adjustments (see text for adjustments). The opaque section of the bars for COI and Funding disclosure denotes the number of publications that would have been recognized had we only used information currently provided on PMC. Such information appears to underestimate the true prevalence of these indicators by two-thirds. (B) Transparency in the open biomedical literature on PMC (2000-2020). Reporting on all indicators of transparency has been increasing for the past 20 years. However, the increase for COI and Funding disclosures has been much more dramatic than for Data sharing, Code sharing, and Protocol registration.

### Transparency across time, countries, fields of science, journals and publishers

Considering the 2,751,420 open access records on PMC (PMCOA) (1959-2020), over time, all indicators have experienced an upward trend in reporting (Fig 3B). However, this increase has been far more dramatic for COI and Funding disclosures than for Data sharing, Code sharing, and Protocol registration. Specifically, all indicators have risen from approximately 0% in 1990 to an estimated 14.9% in 2020 for Data sharing, 3.1% for Code sharing, 90.3% for COI disclosures, 84.7% for Funding disclosures, and 5.0% for Protocol registration. For research articles, the respective proportions for 2020 are 16.6%, 3.5%, 90.6%, 88.8% and 5.7%, respectively. The proportion of publications reporting on these indicators was homogeneous across countries, with most reporting COI and Funding disclosures, but only the minority reporting Data sharing, Code sharing or Protocol registration (Fig 4).

**Fig 4.**
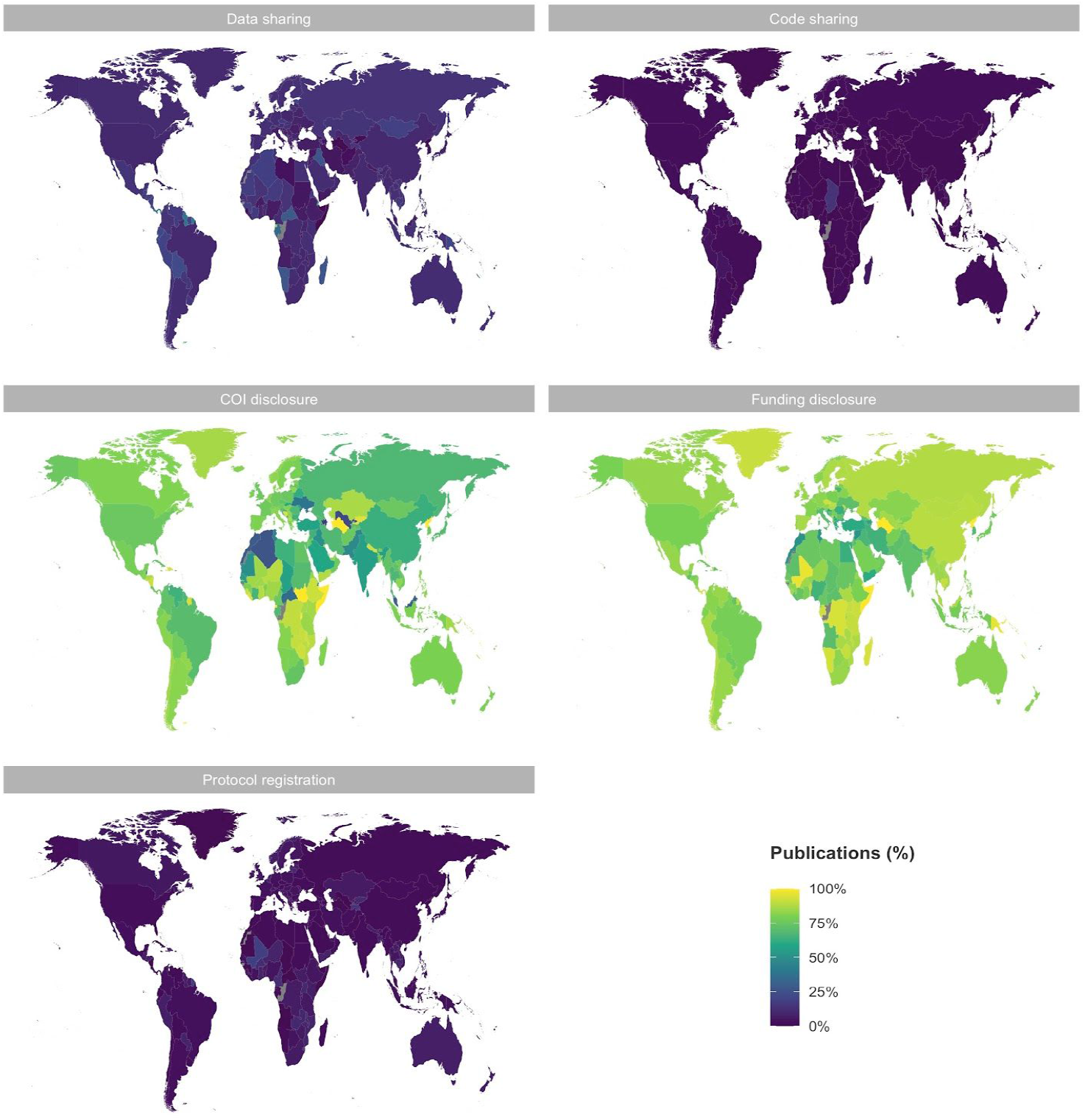
Indicators of transparency across countries of affiliation for all open access articles on PMC (PMCOA) for 1990-2020. Indicators of transparency are roughly homogeneously reported across countries. The light green to yellow maps indicate that the majority of publications from most countries reported COI and Funding disclosures. The purple maps indicate that the majority of publications from most countries did not report any Data, Code or Protocol sharing. Note that the country was only reported in 479,843 articles.

In terms of field of science (see S8 Table for examples), all fields reported COI and Funding disclosures more frequently than other indicators of transparency (Fig 5A; see S9 Table for example phrases across fields and indicators). However, publications from different fields tended to report indicators of transparency at substantially different proportions. For example, publications classified within Biology or Infectious diseases tended to share Data (29.3% and 17.0%, respectively) and Funding disclosures (90.0 and 86.2%, respectively) more than publications within Medicine (7.0% for data, 71.2% for funding) or Health services (3.6% for data, 68.0% for funding). On the contrary, publications within Health services were more likely than other fields of science to share COI disclosures (81.4% vs 77.4%; 95% CI of difference, 3.9-4.2%) or practice open Protocol registration (5.7% vs 2.5%; 95% CI of difference, 3.1-3.3%).

**Fig 5.**
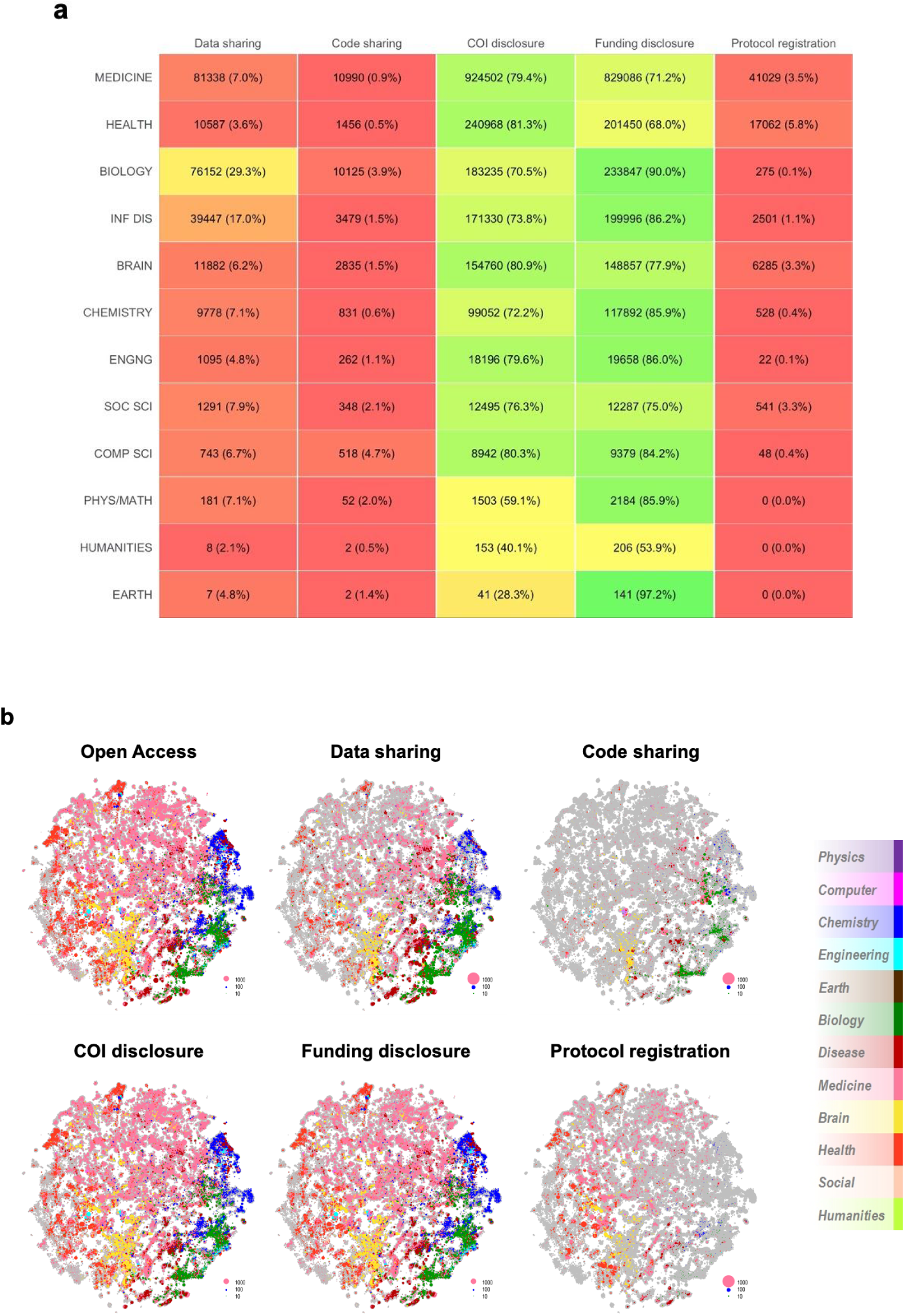
Indicators of transparency across fields of science on PMCOA. (A) Reporting of indicators of transparency across fields of science for all articles on PMCOA since 1990. COI and Funding disclosures are by far the most highly reported indicators within all fields, whereas Code sharing and Protocol registration are by far the least reported. The indicator with the biggest proportional difference between minimum and maximum reporting across fields of science was Protocol registration (0.1% vs 5.8%; Coefficient of variation, 130) and the indicator with the smallest proportional difference was Funding disclosure (53.9% vs 90.0%; Coefficient of variation, 14.5). (B) Indicators of transparency in all articles of PMCOA published between 2015-2019 across galaxies of science. The galaxy in grey represents all clusters of articles published between 2015-2019. On top of the grey galaxy, we overlaid colored representations of the proportion of each cluster that is Open access or reports on any of the indicators of transparency. The Open access galaxy is very similar to that of COI and Funding disclosures, suggesting that most of the open literature reports on both. A number of Chemistry (blue) and Biology (green) clusters are smaller in COI disclosure, whereas a number of Health services (red) and Infectious diseases (burgundy) clusters are smaller in Funding disclosure. Biology (green) and Infectious diseases (burgundy) are pronounced in Data sharing. A very small proportion of clusters report Code sharing or open Protocol registration - of those, the majority are Biology (green) and Health services (red) clusters, respectively.

We then proceeded to further map indicators of transparency across galaxies of science (Fig 5B). These galaxies were previously developed using 18.2 million PubMed articles published between 1996-2019 and divided into ~28,000 clusters of articles [30]. Each cluster comprises similar articles and is colored according to the field associated with the most prevalent journals within the cluster. The size of each colored cluster was modified to reflect the proportion of articles published between 2015-2019 that are open access (1,490,270 articles) or report Data sharing (177,175 articles), Code sharing (25,650 articles), COI disclosure (1,263,147 articles), Funding disclosure (1,192,4996 articles) or Protocol registration (52,798 articles). The most recent 5 years cover more than half of the articles on PMCOA and were chosen to most accurately portray the current practice of transparency. These galaxies further corroborate that most recent open access articles share COI and Funding disclosures, but do not share Data, Code or Protocol.

Out of 2,477 journals with at least 100 articles on PMCOA between 1990-2020, the majority consistently reported COI disclosures (41.7% of journals reported them in ≥90% of their publications; 63.9% in ≥70% of their publications) and Funding disclosures (21.4% of journals reported them in ≥90% of their publications; 53.7% in ≥70% of their publications), but only the minority reported consistently on Data sharing (0.1% of journals in ≥90% of their publications; 0.5% of journals in ≥70% of their publications), Protocol registration (only 1 journal in ≥70% of its publications) or Code sharing (no journal in ≥70% of its publications; highest percentage was 39.7% in GigaScience) (Fig 6A). However, 79.0% of these journals have shared data, 70.8% have shared a protocol and 39.1% have shared code at least once (i.e. 21.0%, 29.2% and 60.9% have never shared any data, protocol or code, respectively). These numbers did not change meaningfully when considering research articles alone.

**Fig 6.**
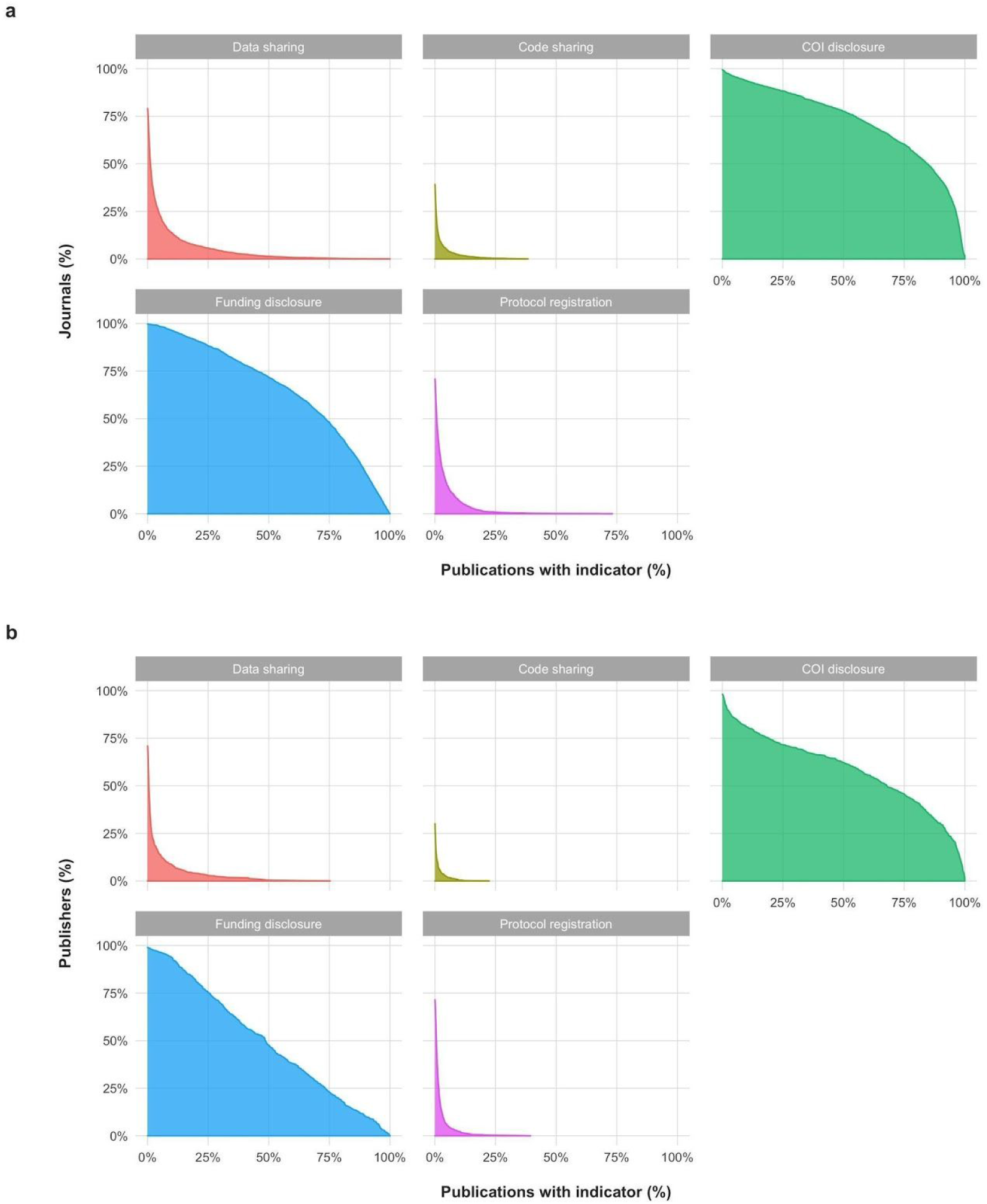
Indicators of transparency across 2,477 journals and 609 publishers with at least 100 publications on PMCOA between 1990-2020. (A) Proportion of journals with at least a designated proportion of publications reporting on each indicator of transparency. For example, for the graph on Funding disclosures, ~50% of journals (vertical axis) report Funding disclosures in at least ~75% of their publications (horizontal axis). Similarly, they indicate that no journal reports Protocol registration in more than ~75% of publications. The concave (bent outward) distribution of COI and Funding disclosures indicates that most publications in most journals disclose COI or Funding, respectively. However, the convex (bent inward) distribution of Data sharing, Code sharing and Protocol registration indicate that most publications do not report on each of these indicators in most journals. About three-quarters of journals have hosted at least one publication sharing data and about one-third of journals have hosted at least one publication sharing code. (B) Proportion of publishers with at least a designated proportion of publications reporting on each indicator of transparency. For example, these graphs indicate that ~50% of journals (vertical axis) report Funding disclosures in at least ~50% of their publications (horizontal axis). Similarly, no publisher reports Data sharing in more than ~75% of publications. As in (A), the shapes of these distributions indicate that most publications of most publishers report on COI and Funding, but most do not report on Data sharing, Code sharing or Protocol registration. Almost three-quarters of publishers host publications that never report Code sharing and roughly one-quarter of publishers host publications that never report Data sharing.

Out of 609 publishers with at least 100 publications on PMCOA between 1990-2020, the majority consistently reported COI disclosures (30.0% disclosures in at least 90% of publications; 48.8% disclosures in at least 70% of publications) and Funding disclosures (10.2% disclosures in ≥90% of publications; 28.4% disclosures in ≥70%), but only the minority reported consistently on Data sharing (1 publisher in ≥70%), Protocol registration (0 in ≥70%) or Code sharing (0 in ≥70%) (Fig 6B). However, 71.4% of publishers have shared a protocol, 70.9% have shared data and 30.0% have shared code at least once (i.e. 28.6%, 29.1% and 70.0% of publishers have never shared protocol, data or code, respectively). These numbers did not change meaningfully when considering research articles alone.

## Discussion

Our evaluation of 2,751,420 open access PubMed Central articles (1959-2020) suggests that there have been substantial improvements in reporting of COI and Funding disclosures across time, fields of science and publication venues, but that Data sharing, Code sharing, and Protocol registration are significantly lagging. It also illustrates that using automated approaches to study and understand the biomedical literature is possible and can yield insights over and above those possible with manual assessments of small random samples. This effort has led to the creation of a database of all open access biomedical literature (2.75 million articles), alongside granular information about indicators of transparency and meta-data integrated from PMC, OCC and SciTech that is now openly available (see our Data Availability statement). We envision that this resource will encourage future efforts trying to further improve on our openly available tools and understand the uptake of transparency and reproducibility practices and their impact on science.

The currently presented algorithms were able to extract relevant text and by construction make multiple pieces of information about their predictions available. For example, in the case of COI disclosures, the algorithm may label a text as referring to no conflicts of interest present, commercial interests present, financial disclosures present, receipt of consulting fees, etc. As such, they can facilitate other teams in building systems on top of this with more nuanced definitions of COI without having to retrain and re-calibrate complex black box systems. Indeed, we hope our data provide the foundational map of transparency to expedite further research in the area.

Nevertheless, given the recent rise in transparency, this is a good opportunity for publishers and the National Library of Medicine (NLM) to standardize reporting of such indicators. Even though the NLM provides XML tags for labelling aspects of an article in a machine-readable way [31], our data illustrate that when these exist (e.g. COI and Funding disclosures) they are used by a minority of journals, and that these do not exist for many other important aspects of transparency (e.g. Protocol registration and Code sharing). Given our data, only a minority of journals seem to use appropriate tags to label transparency statements and no journal used purpose-built tags for Protocol registration or Code sharing. A number of initiatives aim to convert PDFs to tagged files [32], but the institution of a journal-wide requirement for appropriate tagging would enormously reinforce these efforts. In the meantime, both PubMed and other journals can, if they so desire, use our openly available algorithms to improve the availability of information about transparency.

### Limitations

First, the performance of our algorithms was assessed in a recent sample of PMC. As such, we cannot be certain of their performance across a sample of articles outside PMC, older articles, or articles published in the future. Similarly, we cannot be certain that average performance is representative of performance within specific fields of science, journals or publishers. However, the majority of articles in the published literature have been published after 2000 and the findings of our algorithms corroborate those seen in the current and previous manual assessments [20,22]. Second, we are only using the open access subset of PMC. There are other freely available articles within as well as outside PMC that we did not assess. However, previous publications have illustrated that the Open Access Subset is of similar composition as the non-Open Access Subset of PMC [22], and PMC represents about half of the recently published literature available on PubMed. Third, algorithms in general do not guarantee that the statements identified are either accurate, full, or true. This is why we have designed our algorithms to not only extract text, but also indicate why an article was labelled as including information on an indicator. We hope that future work will study these questions further. Finally, some of the XML versions used in our biomedical open literature-wide search do not include the COI or Funding disclosures that were included in the published PDF. Nevertheless, given that the numbers identified are very similar to those of manual labelling, we do not believe that this is a significant concern.

Allowing for these caveats, we provide tools that can expedite the massive assessment of the scientific literature for the most important indicators of transparency. This may help reinforce efforts for making such transparency features even more routine across published articles.

## Materials and Methods

This manuscript was prepared using guidance from the STROBE reporting guidelines [33] for observational studies and the TRIPOD guidelines for reporting prediction models [34].

### Data sources

#### PubMed

First, we randomly assembled a retrospective cohort of 520 records made available on PubMed between 2015-2018. PubMed is maintained by the United States National Library of Medicine (NLM) and provides citations to 30,732,929 references of the biomedical and life sciences literature (as of March, 2020), 4,552,825 of which were published between 2015-2018 [35]. All English articles published between 2015-2018 were eligible for analysis.

#### PubMed Central (PMC)

PubMed Central (PMC) is a free full-text archive for a subset of the publications available on PubMed. It was set up in 2000 and includes publications from journals that have agreed to either share all of their publications, NIH-funded publications or a select subset of their publications [35]. Out of 5,747,776 publications made available on PubMed between 2015-2019 (in terms of Entrez Date), 2,497,046 (43.4%) were also made available on PMC. We randomly identified and downloaded the PDF of 6017 of these records.

#### PubMed Central Open Access Subset (PMCOA)

As of Feb 29, 2020, out of 6,016,911 records ever made available on PMC (1795-2020), 2,754,689 are part of the PMC Open Access Subset (PMCOA) [36]; *not* all articles on PMC are part of PMCOA. These articles are made available under a Creative Commons or similar license and their full text may be downloaded in bulk (as XML files) and used for research; these are the publications that were used to estimate the open access-wide biomedical literature degree of transparency.

#### Sources of meta-data

We extracted all meta-data related to the articles of interest provided by PubMed and PubMed Central (PMC) (e.g. journal of publication, publisher, authors, affiliations, etc.). For all manually assessed articles, we extracted all social media-related data from Altmetric on 30 May 2019. Altmetric captures and tracks the use and sharing of articles across social media. For all manually-assessed articles, we extracted citation counts from Crossref on 30 May 2019; for automatically-assessed articles, we extracted citation counts from Open Citation Collection (OCC; iCite 2.0) [37] on 17 April 2020. OCC is a recent initiative of the National Libraries of Medicine (NLM), which attempts to map all citations from PubMed records to PubMed records and make them openly available. Note that OCC only has citation data for articles with a PubMed ID (PMID), so for articles with a PMCID (PubMed Central ID) but no PMID, we had no citation data. We used 2018 journal impact factors made available by InCites Journal Citation Reports of Web of Science (the latest available at the time). We used the categorizations of PMC articles across fields of science provided by the galaxy of science developed by SciTech (author: K.W.B.) [38]. Briefly, this approach clusters similar articles together and allocates each cluster to the field of the dominant journal within that cluster. Note that using this approach, it may happen that, for example, articles from medical journals are labelled as “Chemistry” if they end-up in a cluster dominated by articles from chemistry journals. Also note that this galaxy of science is based on PubMed, for which reason articles found on PMC but not on PubMed have not been given a field allocation - this also applies to OCC.

### Manual assessment of transparency and reproducibility across 499 PubMed articles (2015-2018)

#### Extraction of article characteristics

For each article, we gathered the following information: PubMed ID (PMID), PubMed Central ID (PMCID), title, authors, year of publication, journal, first author country of affiliation, field of study (as provided by Web of Science (WOS); more information in our protocol [39]) and type of publication (e.g. research article, review article, case series, etc.) as detailed in our protocol^7^.

#### Extraction of indicators of transparency and reproducibility

Two reviewers (S.S. and D.G.C.I.) used a previously published protocol from Wallach et al. [39] to extract appropriate information from eligible articles.

We first extracted any mention of a conflict of interest or funding disclosures from all abstracts and full-text articles in English. Then, for all English records with empirical data (henceforth referred to as “research articles”) we further extracted whether (a) the abstract and/or full-text of each eligible record mentions any protocol registration (whether for the whole study or part of the study), data sharing or code sharing, and whether (b) the abstract and/or introduction of each eligible record implies any novelty (e.g. “our study is the first to identify this new protein”, etc.) or that at least part of this study represents a replication of previous work (e.g. “our study replicates previously reported results in a new population”, etc.). We further extracted whether (a) conflict of interest disclosures mentioned any conflict or not, (b) disclosures of funding mentioned any of public or private funds and (c) whether websites included within data sharing statements were indeed accessible. In addition to the data extracted on the basis of the aforementioned protocol, for each one of the extracted indicators, we also extracted the text in which the information was identified to facilitate our work on automated extraction of these indicators. We only considered clear statements of these indicators, did not attempt to identify whether these statements were complete (e.g. did the authors report all of their conflicts of interest?) or truthful (e.g. has this finding truly never been published before?) and did not consider statements that were not included in the PubMed site or full text.

Note that as per the Wallach et al. protocol and in a deviation from Iqbal et al., which counted all studies with supplementary materials as potentially containing a partial/complete protocol, in this study we downloaded and examined all supplementary materials to verify whether they indeed contain any of data, code or protocol registration. Given differences in languages between fields, it should be noted that by protocol registration we refer to active pre-registration and public availability of a study protocol, such as those found for clinical trials on ClinicalTrials.gov.

Note also that we found the Novelty and Replication indicators particularly ambiguous, for which reason we have created a document with further specifications of non-trivial cases (S1 Text). After compiling this document, we proceeded to have both main reviewers (S.S. and D.G.C.I.) re-assess and cross-check all of their articles, to reduce variability in labelling due to systematic reviewer differences.

Information about 40 randomly identified articles was extracted by three reviewers (S.S., D.G.C.I., J.D.W.). Upon studying discrepancies and clarifying aspects of the protocol, two reviewers (S.S., D.G.C.I.) extracted relevant information, each from 240 articles. Any uncertainties were discussed between all three reviewers to maintain a homogeneous approach. Discrepancies were identified in adjudication of novelty and replication, for which reason these indicators were re-extracted for each article and all unclear articles were discussed between reviewers. Information about concordance is available as a supplement (S1 Table). All extracted data were harmonized into a unified database, which can be accessed on Open Science Foundation (OSF) at https://osf.io/e58ws/?view_only=aadd4846fb3644c8b9528ee96a146f70.

### Automated assessment of transparency: Development

We adjusted a previously reported algorithm developed by N.R. [28] to identify data and code sharing and developed algorithms to identify conflict of interest (COI) disclosures, funding disclosures, and protocol registration statements. All algorithms were constructed to take a PDF file of a typical article available on PubMed and output (a) whether it has identified a statement of relevance, (b) why it is of relevance and (c) what the exact phrase of relevance is. This flexibility was built-in (a) to help integrate these algorithms into the researcher workflow by combining manual and automated inspection, (b) to allow for different definitions of the indicators by different investigators (e.g. consider only COI disclosures that were specifically included as a stand-alone statement, rather than within acknowledgements) and (c) to ease adjudication of their performance. All algorithms, including those for data and code sharing, were further adopted to work with XML files from PubMed Central (i.e. using the NLM XML structure).

Before using any of the algorithms, we preprocessed the text to fix problems with text extraction from PDF files (e.g. inappropriately broken lines, non-UTF8 symbols, etc.), remove non-informative punctuation (e.g. commas, fullstops that do not represent the end of a phrase (e.g. “no. 123”), etc.) and remove potentially misleading text (e.g. references, etc.). For COI disclosures, Funding disclosures and Protocol registration, text was not converted to lower or upper case, we did not use stemming (e.g. we did not convert “processing” into “process”) and we did not remove stop words (e.g. “and”, “or”, etc.) - even though these are frequent preprocessing steps in natural language processing, we found these nuances informative and exploitable. Text was tokenized into phrases for data and code sharing, and tokenized into paragraphs for all other algorithms; for the algorithms we developed from scratch, we used a custom-made tokenizer because already available tokenizers were not found to be accurate or flexible enough. Even though we also considered using machine learning approaches to extract these indicators, we found that the current approach performed well and afforded a level of interpretability and flexibility in definitions that is not easily achievable by alternative methods. All programming was done in R [40] and particularly depended on the packages tidyverse [41], stringr [42] and xml2 [42,43].

The programs were structured into several different kinds of functions. Helper functions (n=7) were developed to help in creating complex regular expressions more easily and in dealing with XML files. Pre-processing functions (n=9) were developed to correct mistakes introduced by the conversion from PDF to text and turn the text into as conducive a document to text mining as possible. Masking functions (n=7) were developed to mask words or phrases that may induce mislabelling (e.g. in searching for funding disclosures, we are masking mentions of finances within conflict of interest disclosures to avoid mislabelling those statements as funding disclosures). Labelling functions (n=81; 20 for COI, 39 for Funding, 22 for Registration) used regular expressions to identify phrases of interest (described below). The regular expressions of these labelling functions also take into account transformations of the text to improve performance (e.g. labelling functions can capture both “we report conflicts of interest” and “conflicts of interest are reported”). Localisation functions (n=3) were developed to identify specific locations within the text (e.g. acknowledgements). Labelling functions that were more sensitive, were only applied within small localised sections of the text to reduce mislabelling. Negation functions (n=7) were developed to negate potentially false labels by the labelling functions. XML functions (n=17) were developed to pre-process and take advantage of the National Library of Medicine (NLM) XML structure. A dictionary was constructed with all phrases and synonyms used by the regular expressions (n=637).

#### Data and code sharing

A member of our team (N.R.) had already developed algorithms to automatically extract information about data and code sharing [28]. Briefly, these algorithms use regular expressions to identify whether an article mentions (a) a general database in which data is frequently deposited (e.g. “figshare”), (b) a field-specific database in which data is frequently deposited (e.g. dbSNP), (c) online repositories in which data/code is frequently deposited (e.g. GitHub), (d) language referring to the availability of code (e.g. “python script”), (e) language referring to commonly shared file formats (e.g. “csv”), (f) language referring to the availability of data as a supplement (e.g. “supplementary data”) and (g) language referring to the presence of a data sharing statement (e.g. “data availability statement”). It finally checks whether these were mentioned in the context of positive statements (e.g. “can be downloaded”) or negative statements (e.g. “not deposited”) to produce its final adjudication. This adjudication (a) indicates whether a data/code sharing statement is present, (b) which aspect of data sharing was detected (e.g. mention of a general database) and (c) extracts the phrase in which this was detected. In this study, these algorithms were customized to avoid text in tables and references and run faster - this is the version of the algorithms that was used to study the PMCOA.

#### Funding disclosures

Briefly, our approach recognizes funding disclosures using regular expressions to identify whether a publication mentions: (a) phrases commonly associated with a funding disclosure (e.g. “This study was financially supported by …”, “We acknowledge financial support by …”, etc.), (b) titles of sections associated with a funding disclosure (e.g. “Funding”, “Financial Support”, etc.), (c) phrases commonly associated with support by a foundation (e.g. “S.S. received financial support by the NIH”, etc.), (d) references to authors (e.g. “This author has received no financial support for this research.”, etc.), (e) thank you statements (e.g. “We thank the NIH for its financial support.”, etc.), (f) mentions of awards or grants (e.g. “This work was supported by Grant no. 12345”, etc.), (g) mentions of no funding (e.g. “No funding was received for this research”) and (h) acknowledgement sections containing phrases with relevant words (e.g. “funded by NIH”, etc.). This algorithm was also designed to avoid mentions of funding related to COI disclosures (e.g. “S.S. has financial relationships with GSK”, etc.).

#### Protocol registration statements

Briefly, we recognize registration statements using regular expressions developed to identify the following: (a) mentions of registration on ClinicalTrials.gov and other clinical trial registries (e.g. “This study was registered on ClinicalTrials.gov (NCT12345678)”, etc.), (b) mentions of registration on PROSPERO (e.g. “This study was registered on PROSPERO (CRD42015023210)”, etc.), (c) mentions of registration of a protocol or a study (e.g. “Our protocol was registered on the Chinese Clinical Trials Register (ChiCTR-IOR-12345678)”, etc.), (d) mentions of research being available on a specific register (e.g. “Our research protocol is available on the ClinicalTrials.gov registry (NCT12345678)”, etc.), (e) titles commonly associated with registration (e.g. “Registration Number”, “Trial registration: NCT12345678”, etc.), (f) previously published protocols of studies (e.g. “Our study protocol was previously published (Serghiou et al. 2018)”, etc.), and (g) registration statements within funding disclosures (e.g. “Funded by the NIH. SPECS trial (NCT12345678)”, etc.). This algorithm was developed to specifically avoid mentions of registry or registration that were not relevant (e.g. “This study enrolled patients in our hospital registry.”, etc.) or registrations with no open protocol availability (e.g. “Our protocol was approved by the IRS (registration no. 123456)”).

### Automated assessment of transparency: Validation across 6017 PMC articles (2015-2019)

#### Data acquisition

The 6017 records obtained from PMC were used to (a) calibrate the algorithms developed in the initial dataset to the PubMed Central dataset and (b) test the algorithms in previously unseen data. We then proceeded by using an importance sampling approach. First, we tested the 6017 records by using the algorithms developed in the training set. Then, we manually assessed 100 of the articles predicted positive by the algorithm and 100 of the articles predicted negative. If at least one mistake was found, we held out 225 articles of the unseen data as a test set and re-developed the algorithm in the remaining data until no meaningful improvement was seen. In the case of registration, only 261 out of 6017 articles were predicted positive, for which reason we only held 161 articles as a test set. More details on our approach can be found in the Supporting Information (S2 Text). All of the aforementioned data and columns indicating which articles were used for training versus testing have been made available (see Data Sharing statement).

#### Algorithm validation

All algorithms were tested in a sample of the PMC that had not been seen before testing (the test set, algorithm predictions and our manual assessment are all openly available, see the Data availability statement). Data sharing was evaluated in 100/764 research articles predicted to share data and 89/5253 predicted to not share data (less than 100 because not all articles were research articles). Code sharing was evaluated in 117/117 articles predicted to share code, in 93/764 articles predicted to share data and in 88/5253 articles predicted to not share data (7 of data sharing and 1 of non-data-sharing articles had been included in the code sharing dataset).

As indicated above, the COI and funding disclosures algorithms were redeveloped in the articles predicted negative out of 6017. To do so, we set aside 225 articles and then redeveloped the algorithms in the remaining data. When confident that further development would not meaningfully improve performance in the test, we tested in the unseen 225. As such, COI disclosures were evaluated in 100/4792 predicted positive and 226/1225 predicted negative (one extra was mistakenly included and not subsequently removed to avoid bias). Funding disclosures were evaluated in 100/5022 predicted positive and 225/995 predicted negative.

For protocol registration we used a modified approach because of its scarcity. First, only 261/6017 articles were predicted positive. As such, the algorithm was re-developed in the first 100 articles predicted positive and tested in the remaining 161. Second, out of 5756 articles predicted negative, it was very unlikely that the general article would be a false negative. As such, we employed an importance sampling approach (similar to that for code sharing), such that for each article we identified whether (a) it mentions any words of relevance to protocol registration (e.g. “registration”, “trial”, etc.), (b) whether it mentions any words related to the title of a methods section (e.g. “Methods”, “Materials and Methods”, etc.), (c) whether it mentions an NCT code, which is the code given to randomized controlled trials by ClinicalTrials.gov, and (d) whether the algorithm re-developed in 100 articles initially predicted positive indicated that any of the previously negative articles should be positive. Out of 5756 predicted negative, we used a stratified/importance sampling procedure by sampling 21/3248 articles deemed irrelevant (i.e. do not mention the words regist*/trial/NCT), 9/451 that were deemed relevant, 44/1951 out of those that were deemed relevant and had a Methods section, 58/91 that contained an NCT identification number and all 15/15 that the new algorithm predicted were positive, but the old algorithm predicted were negative. Detailed breakdown:

| Is relevant | Has methods | Has NCT | True after update | Sample (n) | Total (N) |
| --- | --- | --- | --- | --- | --- |
| No | - | - | No | 21 | 3248 |
| Yes | No | No | No | 9 | 451 |
| Yes | No | No | Yes | 1 | 1 |
| Yes | No | Yes | No | 9 | 24 |
| Yes | No | Yes | Yes | 1 | 1 |
| Yes | Yes | No | No | 44 | 1951 |
| Yes | Yes | No | Yes | 11 | 11 |
| Yes | Yes | Yes | No | 49 | 67 |
| Yes | Yes | Yes | Yes | 2 | 2 |

### Transparency across the open access biomedical literature

First, we downloaded all of the PMCOA with clear commercial and non-commercial use labels in XML format from the PMC FTP Service [36]. Then, we processed all articles in batches of 4,096 articles and in parallel across 8 CPU cores to check for the aforementioned transparency indicators and extract meta-data. Running the COI, Funding and Protocol registration algorithms together in this fashion led to a mean processing time of 0.025 seconds per paper; running the Data and Code sharing algorithms together in this fashion led to a mean processing time of 0.106 seconds per paper. We finally combined the extracted data with data from OCC (citation counts and whether this is a research article) and SciTech field of science.

Note that certain variables from PMCOA were not reported in a consistent standardised form (e.g. country could be reported as “USA”, “United States”, “US”, “Texas, USA”, etc.) - we corrected all variations that occurred 20 or more times into a standardised form (e.g. if “Texas, USA” occurred 20 or more times, we changed it into “USA”) - this is a time-consuming process and we do not believe that standardising variations that occur less than 20 times would meaningfully change the results presented. Data were standardised to the most common label to start with (i.e. if “BMJ” occurred more commonly than “British Medical Journal”, then the two were standardised into “BMJ”), apart from countries, which were standardised to the name given in ggplot2 [44], which is an R package that we used to create maps.

We used these data to create univariable descriptive statistics and frequency counts across journals, publishers, country of affiliation and field of science, as they were reported by PMCOA and SciTech. Unlike the pre-planned analysis of indicator prevalence and distribution across time and field, all other analyses were exploratory. To mitigate data dredging inherent to such exploratory analyses, these analyses were developed in a random sample of 10,000 records and then applied to all data.

All variables considered were: presence and text of COI/funding disclosures, registration statements, title, authors, affiliations, journal, publisher, type of article, references, number of figures, number of tables, citations and field of science. Our *a priori* associations of interest were the distribution of indicators across time and across fields of science, as they are defined by SciTech.

It should be noted that certain journals (e.g. *Scientific Reports* and *Nature Communications*) make conflicts of interest disclosures available in the published PDF version of an article, but do not always include those in the XML versions. As such, our estimate of the presence of conflicts of interest in the open biomedical literature is a valid estimate of which statements are included in the PMC version of the text, but a relative underestimation in terms of what statements are included in the print versions.

### Statistical information

Homogeneity between the two reviewers as well as with the previous reviewer (JW) was assessed by quantifying the frequency of identification of each feature by each reviewer. These frequencies were statistically compared using Fisher’s exact test of independence and calculating a two-sided p-value. Validation of the automated feature extraction algorithms was evaluated using accuracy, sensitivity (=recall), specificity, positive and negative predictive value (PPV, NPV) (PPV=precision), prevalence of the indicator and error between estimated and true prevalence (in terms of absolute difference) (for a detailed explanation of definitions and procedures, see S4 Text). The 95% confidence interval around the diagnostic metrics was built using the non-parametric bootstrap with 5,000 iterations and taking the 2.5th and 97.5th quantiles - in building this confidence interval, we considered the variability introduced by all sampling steps (i.e. sampling 6017 from PMC and sampling 225 from those predicted positive or negative). For the whole PMCOA, we produced univariable frequency statistics for all variables and frequency statistics of each indicator variable across years, journal, publisher, country and field of science. The estimate of indicator prevalence was adjusted by considering the observed PPV and NPV in the test set, such that for an observed prevalence *p*, the adjusted prevalence was *p* × *PPV* + (1 − *p*) × (1 − *NPV*). P-values were produced using non-parametric tests (Kruskal-Wallis for continuous data and Fisher’s exact test for discrete). Correlation coefficients were calculated using the Spearman’s correlation coefficient.

## Data sharing

All data are available at https://osf.io/e58ws/?view_only=aadd4846fb3644c8b9528ee96a146f70.

## Code sharing

All code is available on GitHub at https://github.com/serghiou/transparency-indicators/ and our algorithms are available under a GNU-3 license as an R package called rtransparent on GitHub at https://github.com/serghiou/rtransparent.

## Acknowledgements

We would like to thank the National Library of Medicine, Altmetric, Crossref and the open source community for creating the open data and tools that made this work possible.

