## Supporting Information for "Assessment of transparency indicators across the biomedical literature: how open is open?"

### Supporting Tables

**S1 Table. Indicator identification using the full-text vs the PubMed record of an article from PubMed published between 2015-2018.**

| Indicator | | Full-text | PubMed |
|---|---|---|---|
| | | N = 499 articles | |
| COI disclosure | Yes | 341 (68%) | 36 (7%) |
| | No | 158 (32%) | 463 (93%) |
| Funding disclosure | Yes | 352 (71%) | 177 (36%) |
| | No | 147 (29%) | 322 (64%) |
| | | n = 349 research articles | |
| Data sharing | Yes | 68 (20%) | 3 (1%) |
| | No | 281 (80%) | 346 (99%) |
| Code sharing | Yes | 5 (1%) | 0 (0%) |
| | No | 344 (99%) | 349 (100%) |
| Protocol registration | Yes | 22 (6%) | 12 (3%) |
| | No | 327 (94%) | 337 (97%) |
| Novelty or Replication statement | Yes | 193 (55%) | 110 (32%) |
| | No | 156 (45%) | 239 (69%) |

*Full-text = full-text article; PubMed = the PubMed record of an article. COI = Conflict of Interest. Novelty or Replication statement = language suggesting that the authors are claiming novelty and/or replication (Yes), or neither (No).*

**S2 Table. Indicators of transparency across three different random PubMed samples studying articles from 2000-2014, 2015-2017 and 2015-2018 (current publication).**

| Indicator | | | 2000-2014 | 2015-2017 | 2015-2018 |
|---|---|---|---|---|---|
| All articles | | | N = 442 | N = 148 | N = 499 |
| COI disclosure | Yes | Reported no presence of conflicts | 110 (24.9%) | 87 (58.8%) | 301 (60.3%) |
| | | Reported conflicts | 26 (5.9%) | 10 (6.8%) | 40 (8.0%) |
| | No | | 306 (69.2%) | 51 (34.5%) | 158 (31.7%) |
| Funding disclosure | Yes | Public | 87 (19.7%) | 55 (37.2%) | 122 (24.4%) |
| | | Public + Private | 9 (2.0%) | 2 (1.4%) | 6 (1.2%) |
| | | Public + Private + NGO | 4 (0.9%) | 3 (2.0%) | 6 (1.2%) |
| | | Public + NGO | 54 (12.2%) | 18 (12.2%) | 79 (15.8%) |
| | | Private | 19 (4.3%) | 2 (1.4%) | 12 (2.4%) |
| | | No funding | 12 (2.7%) | 10 (6.8%) | 46 (9.2%) |
| | | NGO | 29 (6.6%) | 12 (8.1%) | 74 (14.8%) |
| | | NGO + Private | 1 (0.2%) | 1 (0.7%) | 7 (1.4%) |
| | No | | 227 (51.4%) | 45 (30.4%) | 147 (29.5%) |
| Research articles | | | n = 260 | n = 95 | n = 349 |
| Code sharing | | Yes | 0 (0.0%) | 0 (0.0%) | 5 (1.4%) |
| | | No | 260 (100.0%) | 95 (100.0%) | 344 (98.6%) |
| Data sharing | | Yes | 5 (1.9%) | 19 (20.0%) | 68 (19.5%) |
| | | No | 255 (98.1%) | 76 (80.0%) | 281 (80.5%) |
| Registration | | Yes | 6 (2.3%) | 31 (32.6%) | 22 (6.3%) |
| | | No | 254 (97.7%) | 64 (67.4%) | 327 (93.7%) |
| Novelty | | Yes | 139 (53.5%) | 64 (67.4%) | 175 (50.1%) |
| | | No | 121 (46.5%) | 31 (32.6%) | 174 (49.9%) |
| Replication | | Yes | 10 (3.8%) | 15 (15.8%) | 33 (9.5%) |
| | | No | 250 (96.2%) | 80 (84.2%) | 316 (90.5%) |

*COI = Conflict of Interest; NGO = Non-Governmental Organisation; N = Number of all articles; n = Number of research articles.*

#### S3 Table. Indicator prevalence by presence or absence from PubMed Central.

| Indicator | | non-PMC | PMC | P-value |
|---|---|---|---|---|
| | | **N = 197** | **N = 302** | |
| **COI disclosure** | No | 37 (18.8%) | 121 (40.1%) | 4.4E-07 |
| | Yes | 160 (81.2%) | 181 (59.9%) | |
| **Funding disclosure** | No | 46 (23.4%) | 101 (33.4%) | 0.016 |
| | Yes | 151 (76.6%) | 201 (66.6%) | |
| | | **n = 145** | **n = 204** | |
| **Data sharing** | No | 91 (62.8%) | 190 (93.1%) | 2.0E-12 |
| | Yes | 54 (37.2%) | 14 (6.9%) | |
| **Code sharing** | No | 141 (97.2%) | 203 (99.5%) | 0.16 |
| | Yes | 4 (2.8%) | 1 (0.5%) | |
| **Protocol registration** | No | 135 (93.1%) | 192 (94.1%) | 0.82 |
| | Yes | 10 (6.9%) | 12 (5.9%) | |
| **Novelty** | No | 71 (49.0%) | 103 (50.5%) | 0.83 |
| | Yes | 74 (51.0%) | 101 (49.5%) | |
| **Replication** | No | 134 (92.4%) | 182 (89.2%) | 0.36 |
| | Yes | 11 (7.6%) | 22 (10.8%) | |

*non-PMC = Articles available on PubMed, but not PMC; PMC = articles available on PubMed and PMC; N = all articles; n = research articles. P-values were calculated using the Fisher's exact test.*

**S6 Table. The 10 most common combinations of indicator co-occurrence in all 2,751,420 open access PubMed Central (PMCOA) publications.**

| Data | Code | COI | Funding | Protocol | Number | % |
|---|---|---|---|---|---|---|
| No | No | Yes | Yes | No | 1,273,382 | 46.3% |
| No | No | No | No | No | 535,428 | 19.5% |
| No | No | Yes | No | No | 339,233 | 12.3% |
| No | No | No | Yes | No | 278,282 | 10.1% |
| Yes | No | Yes | Yes | No | 172,393 | 6.3% |
| No | No | Yes | Yes | Yes | 58,456 | 2.1% |
| Yes | No | No | Yes | No | 38,515 | 1.4% |
| Yes | Yes | Yes | Yes | No | 16,020 | 0.6% |
| No | Yes | Yes | Yes | No | 11,426 | 0.4% |
| No | No | Yes | No | Yes | 5,722 | 0.2% |

**S7 Table. Meta-data across indicators from 2,498,496 articles published from 2000 onwards.**

| Indicator | Variable | Absent | Present | P-value |
|---|---|---|---|---|
| | | Median (IQR) | Median (IQR) | |
| Data sharing | Citation count | 6 (2-14) | 7 (3-17) | $< 10^{-16}$ |
| | Journal Impact Factor | 3 (2-4) | 3 (3-5) | $< 10^{-16}$ |
| | Number of affiliations | 3 (1-4) | 4 (2-5) | $< 10^{-16}$ |
| | Number of authors | 5 (3-8) | 6 (4-9) | $< 10^{-16}$ |
| | Number of figures | 3 (1-5) | 5 (3-7) | $< 10^{-16}$ |
| | Number of references | 34 (19-50) | 48 (34-65) | $< 10^{-16}$ |
| | Number of tables | 1 (0-3) | 2 (0-3) | $< 10^{-16}$ |
| | Year | 2016 (2013-2018) | 2017 (2015-2018) | $< 10^{-16}$ |
| Code sharing | Citation count | 6 (2-15) | 6 (2-15) | 0.20 |
| | Journal Impact Factor | 3 (2-4) | 4 (3-7) | $< 10^{-16}$ |
| | Number of affiliations | 3 (2-4) | 3 (2-5) | $< 10^{-16}$ |
| | Number of authors | 5 (3-8) | 5 (3-8) | 0.97 |
| | Number of figures | 3 (1-5) | 5 (3-7) | $< 10^{-16}$ |
| | Number of references | 35 (20-52) | 46 (31-64) | $< 10^{-16}$ |
| | Number of tables | 1 (0-3) | 1 (0-3) | $3.5 \times 10^{-11}$ |
| | Year | 2016 (2013-2018) | 2018 (2016-2019) | $< 10^{-16}$ |
| COI disclosure | Citation count | 7 (3-18) | 6 (2-14) | $< 10^{-16}$ |
| | Journal Impact Factor | 3 (2-5) | 3 (2-4) | $< 10^{-16}$ |
| | Number of affiliations | 2 (1-3) | 3 (2-4) | $< 10^{-16}$ |
| | Number of authors | 4 (3-7) | 5 (4-8) | $< 10^{-16}$ |
| | Number of figures | 2 (0-5) | 3 (1-6) | $< 10^{-16}$ |
| | Number of references | 22 (5-42) | 38 (25-54) | $< 10^{-16}$ |
| | Number of tables | 0 (0-2) | 1 (0-3) | $< 10^{-16}$ |
| | Year | 2014 (2010-2016) | 2016 (2014-2018) | $< 10^{-16}$ |
| Funding disclosure | Citation count | 4 (2-11) | 6 (2-16) | $< 10^{-16}$ |
| | Journal Impact Factor | 3 (2-4) | 3 (3-4) | $< 10^{-16}$ |
| | Number of affiliations | 2 (1-3) | 3 (2-5) | $< 10^{-16}$ |
| | Number of authors | 4 (2-6) | 6 (4-8) | $< 10^{-16}$ |
| | Number of figures | 1 (0-3) | 4 (2-6) | $< 10^{-16}$ |
| | Number of references | 18 (5-34) | 40 (27-56) | $< 10^{-16}$ |
| | Number of tables | 0 (0-2) | 1 (0-3) | $< 10^{-16}$ |
| | Year | 2015 (2012-2017) | 2016 (2013-2018) | $< 10^{-16}$ |
| Protocol registration | Citation count | 6 (2-15) | 6 (2-13) | $2.3 \times 10^{-9}$ |
| | Journal Impact Factor | 3 (2-4) | 3 (2-4) | $< 10^{-16}$ |
| | Number of affiliations | 3 (2-4) | 4 (3-7) | $< 10^{-16}$ |
| | Number of authors | 5 (3-8) | 7 (5-11) | $< 10^{-16}$ |
| | Number of figures | 3 (1-6) | 2 (1-3) | $< 10^{-16}$ |
| | Number of references | 35 (20-52) | 36 (27-49) | $< 10^{-16}$ |
| | Number of tables | 1 (0-3) | 3 (1-4) | $< 10^{-16}$ |
| | Year | 2016 (2013-2018) | 2017 (2015-2018) | $< 10^{-16}$ |

*The P-value was calculated using the Kruskal-Wallis non-parametric test (tests whether samples originate from the same distribution) - for our data, this is equivalent to the Mann-Whitney U test. P-values < $10^{-16}$ have been replaced by < $10^{-16}$. All values are per article. Citation count was taken from Open Citation Collection and Journal Impact Factor of 2018 from Web of Science.*

#### S8 Table. Representative clusters, journals and reviews for each field of science.

| Field | Clusters | Journals | Reviews |
|---|---|---|---|
| **BIOLOGY** | snake venom composition; legged locomotion; silicon and plant growth | Toxicon; J Exp Biol; Front Plant Sci | Guiding recombinant antivenom development by omics technologies; What factors determine the preferred gait transition speed in humans? A review of the triggering mechanisms; Role of silicon in plant stress tolerance: opportunities to achieve a sustainable cropping system |
| **BRAIN** | intracranial aneurysms, MRA; computed tomography perfusion; transient ischemic attack | AJNR Am J Neuroradiol; Stroke; Stroke | MRA versus DSA for the follow-up imaging of intracranial aneurysms treated using endovascular techniques: a meta-analysis; Computed Tomography, Computed Tomography Angiography, and Perfusion Computed Tomography Evaluation of Acute Ischemic Stroke; Clinical Risk Score for Predicting Recurrence Following a Cerebral Ischemic Event |
| **CHEMISTRY** | microgel particles; coffee beans, bioactive compounds; H2O2 biosensing | Soft Matter; Food Chem; Biosens Bioelectron | Stimuli-Responsive Microgels and Microgel-Based Systems: Advances in the Exploitation of Microgel Colloidal Properties and Their Interfacial Activity; Furan in roasted, ground and brewed coffee; Quantitative analysis of hydrogen peroxide with special emphasis on biosensors |
| **COMP SCI** | image segmentation; adaptive dynamic programming; spatial frequency domain imaging | IEEE Trans Image Process; IEEE Trans Neural Netw Learn Syst; J Biomed Opt | A Survey of Graph Cuts/Graph Search Based Medical Image Segmentation; Distributed Estimation Techniques for Cyber-Physical Systems: A Systematic Review; Advances in the simulation of light-tissue interactions in biomedical engineering |
| **ENGNG** | lead bioaccessibility; droplet microfluidics; polycyclic aromatic hydrocarbons | Sci Total Environ; Lab Chip; Sci Total Environ | Oral Bioavailability of As, Pb, and Cd in Contaminated Soils, Dust, and Foods based on Animal Bioassays: A Review; Continuous magnetic droplets and microfluidics: generation, manipulation, synthesis and detection; Spatial distribution of polycyclic aromatic hydrocarbon contamination in urban soil of China |
| **HEALTH** | self-rated health; total hip arthroplasty; active school transportation | PLoS One; J Arthroplasty; J Phys Act Health | The effect of self-reported health on latent herpesvirus reactivation and inflammation in an ethnically diverse sample; A systematic review and meta-analysis of the direct anterior approach for hemiarthroplasty for femoral neck fracture; Effectiveness of active school transport interventions: a systematic review and update |
| **INF DIS** | Burkholderia mallei; bluetongue virus; ESBL-producing E. coli | PLoS Negl Trop Dis; Vet Ital; Front Microbiol | Melioidosis; Prospects of Next-Generation Vaccines for Bluetongue; Reviewing Interventions against Enterobacteriaceae in Broiler Processing: Using Old Techniques for Meeting the New Challenges of ESBL E |
| **MEDICINE** | colon mucus layer; avian influenza A; malignant hyperthermia | Sci Rep; J Infect; Anesthesiology | Fight them or feed them: how the intestinal mucus layer manages the gut microbiota; Did the Highly Pathogenic Avian Influenza A(H7N9) Viruses Emerged in China Raise Increased Threat to Public Health?; Malignant Hyperthermia in the Post-Genomics Era: New Perspectives on an Old Concept |
| **PHYS/MATH** | Gale crater; cavitation bubbles; sonocatalytic degradation | Astrobiology; Ultrason Sonochem; Ultrason Sonochem | Catalytic/Protective Properties of Martian Minerals and Implications for Possible Origin of Life on Mars; Using power ultrasound to accelerate food freezing processes: Effects on freezing efficiency and food microstructure; Hybrid Advanced Oxidation Processes Involving Ultrasound: An Overview |
| **SOC SCI** | prosocial behavior; peer victimization; causal mediation analysis | J Exp Child Psychol; J Interpers Violence; Epidemiology | The multidimensional nature of early prosocial behavior: a motivational perspective; Annual Research Review: The persistent and pervasive impact of being bullied in childhood and adolescence: implications for policy and practice; Can a Mediator Moderate? Considering the Role of Time and Change in the Mediator-Moderator Distinction |

*This table was created like so: First, we only kept clusters with at least 250 articles between 2015-2019. Then, we randomly sampled 3 clusters within each field. We then identified the name of each cluster, the most prevalent journal within that cluster and the title of a representative review for each cluster. Each cluster, journal and name within each field has been separated by a semi-colon. Two fields were excluded (EARTH, HUMANITIES) due to the 250 papers threshold - these fields have very little presence in PubMed.*

#### S9 Table. Representative text for each indicator of transparency across fields of science.

| Field | Data sharing | Code sharing | COI disclosure | Funding disclosure | Protocol registration |
|---|---|---|---|---|---|
| **BIOLOGY** | the sequencing data of the five degradome libraries are available under ncbi-geo series accession [...] | software used in this manuscript is freely available at https://github com/smith-chem-wisc/gptmd [...] | Competing interests The authors declare that they have no competing interests. | This paper was supported by CNPq, CNPq/INCT-Doen,àö√ üas Tropicais, FAPEMIG, CAPES/PROCAD, and CAPES. | [...] a protocol was registered with the Prospero database (registration number CRD42013005307). |
| **BRAIN** | this data is available for download at the dbgap database (phs000092 v1 p1). | matlab scripts used in this analysis are available at: https://github com/vsevans/cest . | Competing Interests: The authors have declared that no competing interests exist. | This research received no specific grant from any funding agency in the public, commercial or not-for-profit sectors. | Trial registration This trial was registered with ClinicalTrials.gov, number NCT01716481. |
| **CHEMISTRY** | the data set is deposited in the gene expression omnibus (geo) database [ ] under accession no gse6997. | [...] this manuscript was produced with data and code archived at doi:10 5281/zenodo 3403173. [...] | The authors declare no competing financial interests. | Funding. This study was supported in part by the National Natural Science Foundation of China (31470407) [...] | Trial Registration ClinicalTrials.gov NCT01095848 |
| **COMP SCI** | [...] the datasets supporting the conclusions of the article are available in the dataverse [...] | bash and python scripts used to create the nfbs repository [...] are available on github at [...] | The authors declare that the research was conducted in the absence of any [...] conflict of interest. | The authors are grateful to [...] for supporting this research financially under Grants DIP-2012-03 [...] | The following systematic review was registered in PROSPERO with RN: CRD42019120058. |
| **EARTH** | our database ( supplementary data 1 ) consists of ~11500 5-min samples covering ~960 h dwell time [...] | the vpic code is a general-purpose pic simulation and available online ( https://github com/lanl/vpic ). | Competing interests The authors declare no competing financial interests. | Work at ICL was funded by STFC (UK) grant ST/N000692/1. [...] | - |
| **ENGNG** | s1 file (xlsx data availability section and the s4 file 10 1371/journal pone 0221363 | our code is available at: https://githubcom/raharjaliu/microtrench-chemotherapeutic-vision | Competing interests: The authors declare that they have no competing interests. | [...] thank the Alsace Region and the French-German Research Institute of Saint-Louis for funding this work. | Trial Registration Current Controlled Trials ISRCTN91381117 |
| **HEALTH** | raw data is attached to this article. | python code and associated search keywords are publicly available on a repository [...] | Conflicts of Interest: None declared. | This project is supported by General Research Fund of the Research Grants Council of Hong Kong (HKU 769408 M). | Systematic review registration PROSPERO 2015: CRD42015017327 |
| **INF DIS** | sequence data [...] have been deposited in genbank with the accession codes mf942137-mf942331. | code for these simulations is available at https://github com/braindynamicsusyd/spikenet | Competing interests The authors declare that they have no competing interests. | Funding. This work was supported by National Institutes of Health Grant CA 19014 (to NR-T). | Trial Registration ClinicalTrials.gov NCT00295581 |
| **HUMANITIES** | supplemental material available at figshare: https://doi org/10 25386/genetics6304502 . | the source code was highly valuable-arguably more so than the software itself-because [...] | The authors declare they have no competing financial interests. | Funding: The author received no specific funding for this article. | - |
| **MEDICINE** | all data are available from the geo accession numbers gse39040 gse28425 gse36004 and cse79181 [...] | custom analysis and heatmap generation code is available from https://githubcom/petercomb [...] | The authors declare no competing financial interests. | Funding Not applicable. | Trial registration ClinicalTrials.gov ID NCT00371540 |
| **PHYS/MATH** | all data [...] have been made publicly available [...] through the following dois: 10 5281/zenodo 3238621 [...] | it is opensource and freely available on github [ref] | Competing interests The authors declare that they have no competing interests. | This study was supported by L'Agence Nationale de la Recherche (ANR); reference: ANR-09-BLAN-0093-03. | - |
| **SOC SCI** | the data are available through dryad at the following link: https://doi org/10 5061/dryad 53t31 . | we include r code for step-by-step instructions for our analysis in the supplemental material [...] | Competing interests: The authors declare that they have no competing interest. | Funding The research reported was not externally funded. | Trial registration Australian New Zealand Clinical Trials Registry ACTRN12611000438954 |

*These phrases were chosen like so: First, a random sample of 10 articles was identified for each field-indicator pair (seed 1515). Then, the sample was ordered in terms of descending citation counts and was inspected from top to bottom for succinct representative phrases. The text was truncated to fit in three lines and any truncated text is denoted as "[...]". The extracted text for "Code sharing" in Humanities is a false positive - 2 (0.5%) texts from the Humanities were labelled as sharing code, both of which were false positives; nevertheless, both texts were highly relevant to code. For the code and data used to construct this table, see Data and Code Availability statements.*

#### S10 Table. Reviewer concordance.

| Indicator | | J.D.W. | D.G.C.I. | S.S. | P-value |
|---|---|---|---|---|---|
| Data sharing | No | 76 (80.0%) | 138 (83.6%) | 143 (77.7%) | 0.369 |
| | Yes | 19 (20.0%) | 27 (16.4%) | 41 (22.3%) | |
| Code sharing | No | 95 (100.0%) | 162 (98.2%) | 182 (98.9%) | 0.454 |
| | Yes | 0 (0.0%) | 3 (1.8%) | 2 (1.1%) | |
| COI disclosure | No | 51 (34.5%) | 69 (30.3%) | 89 (32.8%) | 0.673 |
| | Yes | 97 (65.5%) | 159 (69.7%) | 182 (67.2%) | |
| Funding disclosure | No | 45 (30.4%) | 65 (28.5%) | 82 (30.3%) | 0.880 |
| | Yes | 103 (69.6%) | 163 (71.5%) | 189 (69.7%) | |
| Protocol registration | No | 90 (94.7%) | 155 (93.9%) | 172 (93.5%) | 0.967 |
| | Yes | 5 (5.3%) | 10 (6.1%) | 12 (6.5%) | |
| Novelty | No | 31 (32.6%) | 74 (44.8%) | 100 (54.3%) | 0.002 |
| | Yes | 64 (67.4%) | 91 (55.2%) | 84 (45.7%) | |
| Replication | No | 80 (84.2%) | 151 (91.5%) | 165 (89.7%) | 0.194 |
| | Yes | 15 (15.8%) | 14 (8.5%) | 19 (10.3%) | |

*Reviewer concordance was very good between the two new reviewers (S.S. and D.G.C.I), as well as the two reviewers and the previous reviewer (J.D.W.). Appreciable deviations were only seen in the assessment of Novelty, presence of a Replication component and Funding disclosures, the latter of which reached statistical significance (95% CI, 0-18%). All three of these were manually re-extracted and adjudicated by both reviewers to ascertain that these discrepancies did not reflect systematic differences in extraction.*

### Supporting Figures

**S1 Fig. Types of data sharing statements and funding disclosures in 349 research articles (2015-2018).**

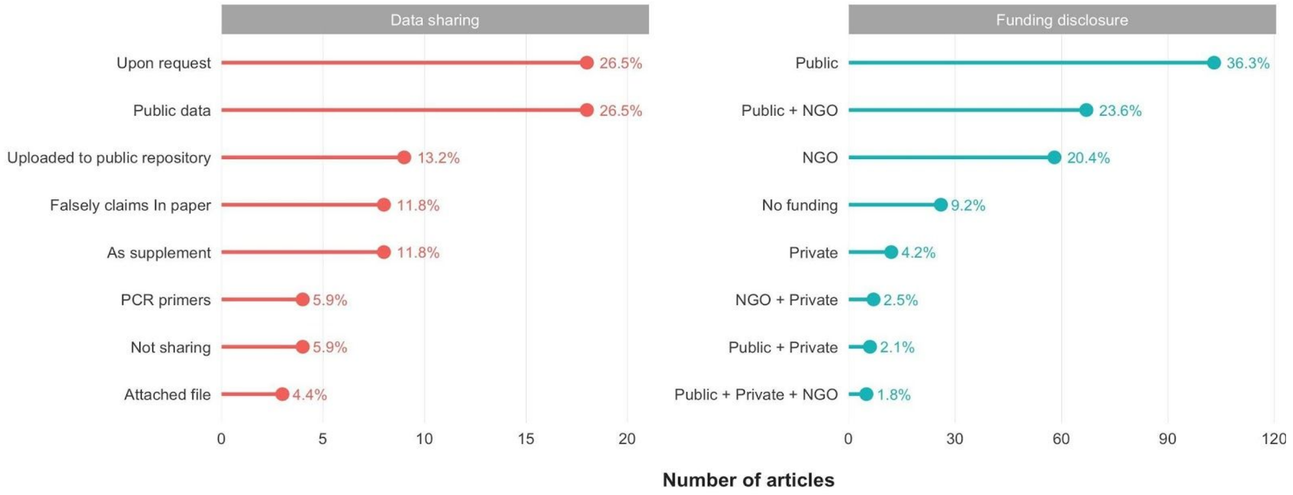

*Of 68 research articles with Data sharing statements, most claimed availability upon request or made use of public data. Of those actively sharing new data, most made their data available on an online repository (e.g. GenBank); 8 articles stated that all of their data were available in the text or supplements, but we could not locate any such raw data - all 8 were published in PLOS One; 4 articles only shared PCR primers; 4 articles actively indicated that they are not currently sharing their data. Of 284 research articles with Funding disclosures, most reported public funds (e.g. National Institutes of Health) or funds from non-governmental organisations (e.g. Gates Foundation). Very few indicated no or private funding. NGO = Non-Governmental Organisation.*

### S2 Fig. Example predictions.

| | | Positive | Negative |
|---|---|---|---|
| **COI disclosure** | **Both correct** | Competing interests: The authors declare that no competing interests exist. | - |
| | **Updated correct** | D.J.C., J.C., I.N., and C.C. are former employees of Hoffmann-LaRoche. | - |
| | **Both wrong** | F.Z. is a founder of Editas Medicineand a scientific advisor for Editas Medicine and Horizon Discovery. | - |
| **Funding disclosure** | **Both correct** | Funding: This work was supported by the National Institutes of Health (NIH) via DP2 OD001886 and R01AI079497. | - |
| | **Updated correct** | This work was accomplished with a generous support from the Bay Area Lyme Foundation. | - |
| | **Both wrong** | Antonio Trincone acknowledges BENTEN project within the Biotechnology Network of Campania Region (Italy). | - |
| **Protocol registration** | **Both correct** | Trial registration: This trial is registered with Chinese Clinical Trials Register, ChiCTR-IOR-17010860 | - |
| | **Updated correct** | (Funded by the National Institutes of Health; CHAMP ClinicalTrials.gov number, NCT01581281). | Ethics approval and consent to participate Ethical approval for the study was obtained by the Ghent University Hospital Ethics Committee (registration no B67020109977). |
| | **Both wrong** | Methods: EDITION 2 (NCT01499095) was a randomized, 6-month, multicentre, open- label, two-arm, phase IIIa study investi- gating once-daily Gla-300 versus Gla-100, plus OADs (excluding sulpho- nylureas). | Data were derived from mCRC patients included in two large phase III studies: CAIRO (Clinicaltrials.gov NCT0031200) and CAIRO2 (Clinicaltrials.gov NCT 00208546), of which the results have been published previously [21, 42]. |
| **Data sharing** | **Correct** | All data and r-syntax are available at the open science framework data repository https://osf.io/27bm4/. | - |
| | **Wrong** | The.xlsx data used to support the findings of this study are included within the supplementary information file(s). | Since there is no human BRCA2 structure containing our mutations available in the Protein Data Bank (PDB) we used a mouse BRCA2 structure PDB. |
| **Code sharing** | **Correct** | The r code for each method is provided in Note S2 in File S1. | - |
| | **Wrong** | The scripts and instructions of our algorithm are included online on Github (www.github.com/kandelj/MitoSPT). | Recently the TurkerGaze project [31, 96] was made available on GitHub. |

*This figure illustrates examples of extracted text that was correctly labelled by both algorithms (the algorithm developed in the initial sample of 499 articles and the updated algorithm using data from the 6017 articles), correctly labelled only by the updated algorithm (i.e. the one updated using data from the 6017 articles) and text that was incorrectly labelled by both algorithms. The Positive column refers to statements with the indicator of interest and the Negative column to statements without the indicator of interest. Notice that the green statements for COI disclosures, Funding disclosures and Protocol registration are very explicit about their content, the orange statements slightly less so and the red statements even less explicit - this illustrates how the algorithms were updated to capture more of the less explicit statements (see Supporting Information: Algorithm development for more details). Note that these statements were purposefully selected because they are small and clearly exemplify the points made - to access the complete evaluation of these algorithms and all sentences classified correctly or incorrectly, please see our data (see Data Availability in Methods).*

### Supporting Text

#### S1 Text. Novelty and replication.

Novelty and replication are rather ambiguous. We hereby attempt to clarify which phrases were deemed to imply novelty and which were deemed to imply replication.

##### *Novelty*

Phrases or words deemed to clearly indicate that the authors imply novelty:

1. Mentions of words such as: new/novel/revolutionary/ground-breaking/ innovative/etc.
    a. e.g. PMID 28035953 (abstract): "Three novel strains of photosynthetic bacteria from the family Ectothiorhodospiraceae were isolated"
2. Mentions of phrases such as: for the first time/never done before/not previously evaluated/etc.
    a. e.g. PMID 25758201 (introduction): "We, for the first time, show the involvement of GFAP-expressing cells in ductular reaction in chemically induced liver cirrhosis and show that there is no direct EMT in terms of myofibroblast development."
3. Mentions of phrases such as: currently unknown/speculative/overlooked/etc.
    a. e.g. PMID 27873074 (introduction): "Results regarding the association between adipokine levels and bone mineral density (BMD) have been inconsistent; the effects of sex, menopause, and central obesity remain unknown. We evaluated the association between serum leptin, adiponectin, and high-molecular-weight (HMW) adiponectin levels and BMD according to menopause and central obesity status in Korean women."

4. Mentions of phrases such as: we discovered that x is associated to y/our data show a novel function
5. Mentions of phrases such as: we developed an apparatus/tool/method/etc. to study x
    a. e.g. PMID 27087363 (introduction): "[...] a disposable bag clamped to a reusable base (Figure 1-B) was developed to enhance washout while allowing for the complete elimination of air during in vitro testing."

Phrases or words not deemed to *clearly* indicate that the authors imply novelty:

1. Mentions of phrases such as: it is still unclear/not fully understood/poorly understood/incompletely understood/remains elusive/only a few studies/little is known/limited knowledge/inconsistent findings/remains controversial/etc.
    a. e.g. PMID 29154423 (abstract): "Because of the effect of pre-load vs after-load on these mechanisms is not completely understood, we studied the effect in isolated muscle strips."
    b. e.g. PMID 27633504 (abstract): "It is not fully understood where and how people are exposed to sensitizing metals. Much can be learnt from studying occupational settings where metals are handled."
2. Mentions of phrases such as: provides further evidence/deepen our understanding/incremental knowledge/etc.
    a. e.g. PMID 26373460 (abstract): This study provides further evidence for the post-regression stability that characterises Rett syndrome. Emergent low mood in Rett syndrome requires further research."

b. e.g. PMID 29321952 (abstract): “Based on previous evidence suggesting a possible nuclear role for FEZ1, we wanted to deepen our understanding of this function by addressing the FEZ1-RAR interaction. We performed in vitro binding experiments and assessed the interface of interaction between both proteins.”

Phrases that depend on the context:

1. Mentions of phrases such as: extended this concept/extended this query/etc.

   a. e.g. PMID 25870189 (introduction) - implied novelty: “This method is proposed for assessing RhoA-Rho kinase activity in patients with hypertension and other cardiovascular diseases. In this study, we extended this concept to the analysis of GEF activity in human.”

***Replication***

Phrases or words deemed to *clearly* indicate that the authors imply some kind of replication:

1. Mentions of words such as: replication/replicate/repeat/validate/confirm/etc.

   a. PMID 28587893 (introduction): “It is therefore an important model in which to validate the use of IF and provide baseline data on neuronal numbers and neuronal density.”

   b. PMID 25625488 (introduction): “The aim of this study was to confirm this association and to elucidate the effects of the variant on protein function and Alzheimer-type pathology.”

2. Mentions of phrases such as: examine the estimate/accuracy/reliability.

    a. PMID 29274866 (introduction): "However, limited data on NS5A-inhibitor treatment escape have been obtained in such systems. We aimed at providing an independent head-to-head comparison of the efficacy of all currently licensed HCV NS5A inhibitors against the major HCV genotypes and important subtypes, as well as resistant escape variants, in infectious culture systems."

3. Mentions of phrases: long-term follow-up study/etc.

    a. PMID 20469608 (introduction): "The 18-year follow-up of the PIAF cohort was recently completed and airway responsiveness to histamine was assessed in all available subjects."

Phrases or words not deemed to clearly indicate that the authors imply some kind of replication:

1. Mentions of phrases: little is known/poorly understood/not clear/debated/limited information/etc.

    a. PMID 30051367 (introduction): "While some reports have indicated that sarcopenia is not associated with poor outcomes in resectable GC, other reports have indicated that visceral and subcutaneous fat are equivocal prognostic factors. 16 Thus, there are no clear data regarding the prognostic value of preoperative body composition parameters in patients of resectable GC."

2. Mentions of phrases: provide further evidence/further characterize/more data/data are scarce/information is limited/only one study/etc.

    a. PMID 26373460 (introduction): “The purpose here was to add to the evidence about age-related clinical and behavioural change into adulthood.”

3. Mentions of phrases: consistent with previous findings/etc.

    a. PMID 30030228 (abstract): “Consistent with previous publications, decreased membrane fluidity was associated with increased fatty acid production ability”

##### *Novelty and Replication*

Several articles mention words/phrases that classify them into both categories:

1. Mentions of phrases such as: repeated/tested/validated in a new sample/population/condition.

    a. e.g. PMID 27570184 (introduction): “The majority of previous studies were conducted in Western settings hence studies that explore attentional bias and eating behaviors among dieters and nondieters from Eastern cultures could shed light on relevant cultural differences.”

    b. e.g. PMID 27477952 (introduction): “Although JHFRAT has been validated in western countries, it may not be suit-able to use on Chinese population. Nunnally and Bernstein (1994) have also stated that validation of measurements is an unending process and that the validity of each use within a specific context must be documented empirically. Therefore, this study tried to test the validity and reliability of JHFRAT for its implication in Chinese population.”

#### S2 Text. Algorithm validation.

##### Data sharing

Our aim was to evaluate the performance of the data sharing identification algorithm by N.R. in a set of 6017 randomly identified articles of 2015-2019 from PubMed Central (PMC). Specifically, we were interested in the expected sensitivity, specificity, accuracy and AUROC of this algorithm for various definitions of data sharing in English research articles found on PMC. Note that by research articles we refer to any articles with empirical data and we do not include case studies, systematic reviews or meta-analyses.

In considering possible definitions of data sharing, we first define the following modes of data sharing: (a) upon request – a statement that data are being made available upon request, (b) effective – immediately accessible data (e.g. URL works, etc.), (c) active – new data made available online (a subset of effective), (d) per-paper - the paper refers to immediately accessible data (regardless of whether the URL works or not) (a superset of effective), (e) public – use of previously generated public data and (f) false claims – data sharing statements that make false claims (e.g. authors claim all data were made available but only some was made available, authors claim that data are made available in the text but only statistics are made available, etc.). It should be noted that the data sharing algorithm was designed to fulfil the definition of "active" data sharing and this is the definition being used in our automated assessment of the open access articles on PMC Open Access, as presented in the main manuscript.

On the basis of these, we constructed the following definitions of data sharing: (a) *any statement* sharing refers to any mention of openly available data or intent to share data (= upon request + effective sharing + false claims); (b) *effective* sharing refers to sharing any immediately available data (= any statement – upon request – false claims (no raw data, page not found)); (c) *active* sharing refers to effective data sharing of new data (= effective data sharing – public); (d) *per-paper* sharing refers to articles that, as far as we can tell from the paper alone, make at least some of their data immediately available (= effective data sharing + page not found (i.e. the subset of ineffective data sharing that we could tell is ineffective from the paper itself)); (e) *public* sharing refers to the availability of any sharing excluding that of public data (= any statement - public); (f) *upon request* refers to any data sharing excluding statements of upon request (= any statement – upon request). Note that if an article contained more than one type of data sharing, the most broadly applicable mode was selected (for example, if an article both made data available on GEO and also had a statement of data sharing upon request, we counted the former).

Performance was first assessed in 100/764 randomly identified articles that the algorithm predicted to share data. Within these data, two reviewers confirmed 100 research articles in English, of which 5 made a false claim (3 claimed to provide all data but only provided some data, 2 claimed to share data on an inaccessible URL and 1 claimed to share raw data where none could be found) and 4 used public data. As such, across different definitions, there were 100 true positives (TPs) for any data sharing, 97 TPs for effective data sharing, 93 TPs for active data sharing, 99 TPs for per-paper sharing, 96 for public data sharing and 100 for upon request sharing.

| is_open_data | isData | isData_comments | n |
|---|---|---|---|
| TRUE | FALSE | Public data | 3 |
| TRUE | TRUE | - | 90 |
| TRUE | TRUE | Data sharing (by SS) | 2 |
| TRUE | TRUE | False claim (no raw data) | 3 |
| TRUE | TRUE | False claim (page not found) | 1 |
| TRUE | TRUE | False claim (partial raw data); False claim (page not found) | 1 |

Performance was then assessed in 116/5253 randomly identified articles that the algorithm predicted to not share data. Within these articles, two reviewers confirmed 116 English articles, of which 89 were research articles; the three most common types of non-research articles were non-systematic reviews (6/27), case reports (6/27) and systematic reviews (4/27). Of the 89 research articles, 15 claimed at least some data sharing, of which 5 provided at least some effective data, 5 made false claims about data sharing (4 claimed raw data where none existed, 1 provided an inaccessible URL) and 1 at least partly used public data. This implies that there were 74 true negatives (TNs) for any data sharing, 83 TNs for effective data sharing, 84 TNs for active data sharing, 84 TNs for per-paper sharing, 75 TNs for public data and 78 TNs for upon request.

| is_open_data | isData | isData_comments | n |
|---|---|---|---|
| FALSE | FALSE | - | 74 |
| FALSE | TRUE | False claim (no raw data) | 4 |
| FALSE | TRUE | Upon request | 4 |
| FALSE | TRUE | As supplement | 3 |
| FALSE | TRUE | False claim (page not found) | 1 |
| FALSE | TRUE | Partial (GEO number) | 1 |
| FALSE | TRUE | Partial (primers) | 1 |
| FALSE | TRUE | Public data - exact dataset not provided | 1 |

Of the 5 missed by per-paper data sharing, the GEO was missed because the tokenizer split the GEO from the GSE number (i.e. it split the database name from the dataset ID number), the primer was missed because the algorithm does not capture primers and the 3 supplements were missed because the algorithm does not capture “supplementary information”, “supplemental material” and only considers supplements that mention some kind of “supplemental table”. The algorithm could be improved by (a) fixing the tokenizer (do not split on all full stops, only those that symbolize next sentence) (fixes article 01665), (b) identifying .xlsx as referring to sharing a supplementary table (fixes article 00853) and (c) making the algorithm recognize S1 Table (as well as the currently recognizable Table S1) (fixes article 02339). The primer article (article 02236) is difficult to fix and the remaining article's supplement could not have been recognized as true data sharing without access to the supplement (00856).

Overall, by extrapolating our sample to the full 6017 articles, we would expect the following performance.

| Definition | TP | FP | TN | FN | Sens | Spec | Acc | AUROC |
|---|---|---|---|---|---|---|---|---|
| **Predicted** | **n = 764 research** | | **n = 4030 research** | | - | - | - | - |
| **Any statement** | 764 | 0 | 3351 | 679 | 52.9 % | 100 % | 85.8 % | 76.5 % |
| **Effective** | 741 | 23 | 3759 | 271 | 73.2 % | 99.4 % | 93.9 % | 86.3 % |
| **Active** | 711 | 53 | 3804 | 226 | 75.9 % | 98.6 % | 94.2 % | 87.3 % |
| **Per paper** | 756 | 8 | 3804 | 226 | 77.0 % | 99.8 % | 95.1 % | 88.4 % |
| **Public** | 733 | 31 | 3396 | 634 | 53.6 % | 99.1 % | 86.1 % | 76.4 % |
| **Upon request** | 764 | 0 | 3532 | 498 | 60.5 % | 100 % | 89.6 % | 80.3 % |

By correcting the GEO and fixing the two supplements, by the per-paper definition, we would get 756 TP, 8 FP, 3940 TN and 90 FN, thus, 89.4% sensitivity, 99.8% specificity, 98.0% accuracy and 94.6% AUROC.

##### *Summary*

Upon manual inspection of a random sample of 100 research articles of 2015-2019 from PMC that were labelled as actively sharing data, 93 were indeed found to actively share data, but 7 were not - 4 used publicly available data, 2 referred to an inaccessible URL and 1 claimed that all raw data were in the text where none could be found. Similarly, out of 89 research articles labelled as not actively sharing data, 84 were found to not share data, but 5 did - 3 made their data available as supplements, 1 referred to a GSE number and 1 contained a primer sequence. Assuming similar proportions across all 6017 articles in our sample, in terms of active data sharing in research articles, this algorithm has an accuracy of 94.2% (95% CI, 89.7-97.99%), a sensitivity of 75.8% (95% CI, 61.4-93.9%) and a specificity of 98.6% (95% CI, 97.6-99.5%). Applying this algorithm across research articles of 2015-2019 on PMC is likely to underestimate the true proportion of data sharing by an absolute value of 3.6% (i.e. for every 4794 random PMC research articles of 2015-2019, we expect this algorithm to label 764 positive, whereas 935 are actually positive).

#### Code sharing

Our aim was to evaluate the performance of the code sharing identification algorithm developed by N.R. in a set of 6017 randomly identified articles of 2015-2019 from PubMed Central (PMC). Specifically, we were interested in the expected sensitivity, specificity, accuracy and AUROC of this algorithm for various definitions of code sharing in English research articles found on PubMed Central. Note that by research articles we refer to any articles with empirical data and we do not include case studies, systematic reviews or meta-analyses.

In considering possible definitions of code sharing, we first defined the following modes of code sharing: (a) any – any reference to openly available code or intent to share code, (b) effective – immediately accessible code (e.g. URL works, etc.), (c) active – statement that new code is being made available online (a subset of effective), (d) per-paper - the paper states that the code is immediately accessible (regardless of whether the URL works or not) (a superset of effective), (e) public – use of previously written code (e.g. a public tool) and (f) false claims – code availability statements that make false claims (e.g. authors claim all code was made available but only some was made available, etc.).

On the basis of these, we constructed the following definitions of code sharing: (a) *any statement* sharing refers to any mention of openly available code or intent to share code (= any); (b) *effective* sharing refers to sharing any immediately available code (= any statement – false claims); (c) *active* sharing refers to effective sharing of new code (= effective – public); (d) *per-paper* refers to articles that, as far as we can tell from the paper alone (without the supplement or accessing the web), make at least some of their code immediately available (= effective + page not found (i.e. the subset of ineffective code sharing that we could tell is ineffective from the paper itself)); (e) *public* sharing refers to the availability of any code sharing excluding public code (= any - public). Note that if an article contained more than one type of code sharing, the most broadly applicable mode was selected (for example, if an article both used public code and made a false claim that all code was available, we counted only the public code).

Performance was first assessed in all 117 articles that the algorithm predicted to share code. Two reviewers confirmed 110 research articles in English, of which 7 did not mention any code sharing, 4 referred to at least some use of publicly available tools and 3 represented ineffective code sharing (inaccessible URL; one of these three also used a publicly available tool). This implies that there were 103 true positives (TPs) for any statement sharing, 101 TPs for effective sharing, 97 TPs for active sharing, 103 TPs for per-paper sharing and 99 TPs for public sharing.

Of the articles falsely predicted to share code across different definitions of code sharing, 3 included references that mentioned GitHub, 2 mentioned GitHub without providing code on GitHub (e.g. "this dataset is available on GitHub") and 2 mentioned words of the main text misconstrued to refer to code sharing (e.g. “coding sequence”). The 3 mentions of GitHub from the references and the mention of a coding sequence are aspects of this code that can be fixed more easily than the rest.

| isCode_comments | n |
|---|---|
| - | 95 |
| Mentions GitHub (references) | 3 |
| Public tool | 3 |
| Shares code (by SS) | 2 |
| Checklist item about best practice for other | 1 |
| False claim (page not found) | 1 |
| Mentions "coding sequence" and "available bioinformatics tools" | 1 |
| Mentions GitHub (main text) | 1 |
| No code (by SS); Mentions GitHub | 1 |
| Public tool; False claim (page not found) | 1 |
| Shares code (by SS); False claim (page not found) | 1 |

Performance of code sharing was also assessed in the sample of articles assessed for data sharing. Of 100/764 randomly identified articles that the algorithm predicted to share data and 116/5253 articles that the algorithm predicted to not share data, there were 189 research articles in English.

Of 100/100 research articles predicted to share data, 7 were also predicted to share code (i.e. 93 were predicted to not share code). Of 93 predicted to not share code, 89 were indeed true negatives (TNs), but 4 were false negatives (FNs). Of 4 FNs, one mentioned BitBucket, one mentioned R-syntax (instead of R script), one mentioned GitHub within a URL rather than its own word (the algorithm is looking for "\\bgithub\\b", which only recognizes the word GitHub) and one was missed because it made its code available on ResearchGate (albeit this was a false claim as the code was not found on ResearchGate upon looking). The algorithm could be updated to identify all 4 of these cases in terms of per-paper code sharing.

Of 89/116 research articles predicted to not share data, 88 were predicted to not share code (i.e. negative) and 1 was predicted to share code (i.e. positive). Of 88 predicted negative, 1 was a FN and shared its code as a supplement: "Our methods were easy to implement in R, and the code is presented in Supplementary Table 1". There were no FPs. This could be fixed by identifying the combination of code with supplement, in a similar fashion to the identification of data in combination with supplement.

Performance metrics were calculated by extrapolating the observed performance across the expected number of research articles within all 6017 random articles. To calculate overall performance we used the articles predicted to share code to estimate TP/FP and the articles stratified by data sharing to estimate TN/FN. Given the association of data sharing with code sharing, this provided a more efficient approach to estimating performance and is formally known as importance or stratified sampling.

As an example, we hereby illustrate how this approach was used to calculate the performance metrics for Any statement. TP = 103; FP = 7; TN = 90/93 * 93/93 * 703 + 87/88 * 88/114 * 5197; FN = 3/93 * 93/93 * 703 + 1/88 * 88/114 * 5197. Notice that we are using 703 instead of 764 and 5197 instead of 5253 because we are only considering those predicted positive or negative for data sharing within the articles not sharing code (i.e. 6017 - 117 = 5900). Also notice that we are using 93 instead of 100 and 114 instead of 116 because we are only using observations that were predicted negative for code sharing - this can be done without complications because of random sampling (i.e. sampled positive are independent of sampled negative). Finally, notice that the total number of expected research articles is slightly different from that in data sharing (4825 vs 4794) because this was estimated using a different sample of the data; this uncertainty was taken into account when calculating the 95% confidence intervals presented in the main text.

| **Definition** | **TP** | **FP** | **TN** | **FN** | **Sens** | **Spec** | **Acc** | **AUROC** |
|---|---|---|---|---|---|---|---|---|
| **Predicted** | **n = 110 research** | | **n = 4715 research** | | - | - | - | - |
| **Any statement** | 103 | 7 | 4639 | 76 | 57.5 % | 99.8 % | 98.3 % | 78.7 % |
| **Effective** | 101 | 9 | 4647 | 68 | 59.8 % | 99.8 % | 98.4 % | 79.8 % |
| **Active** | 97 | 13 | 4647 | 68 | 58.8 % | 99.7 % | 98.3 % | 79.3 % |
| **Per paper** | 103 | 7 | 4639 | 76 | 57.5 % | 99.8 % | 98.3 % | 78.7 % |
| **Public** | 99 | 11 | 4639 | 76 | 56.6 % | 99.8 % | 98.2 % | 78.2 % |

With the suggested updates to the algorithm (reference avoidance, "coding sequence" avoidance and inclusion of BitBucket, R-syntax, ResearchGate, www.github.com, and supplementary tables) the per-paper performance would been perfect (i.e. 100% for all performance metrics), even though this is an overestimate due to overfitting.

***Summary***

Upon manual inspection of all 110 research articles of 2015-2019 from PMC that were labelled as actively sharing code, 97 indeed shared at least some code, whereas 4 did not – all 4 of these mentioned using code by publicly available code, e.g. "local realignment and variation call were analyzed using Samtools10 Picard (http://broadinstitute.github.io/picard/) and GATK". Similarly, out of 181 articles labelled as not sharing code, 177 indeed did not share code, but 4 did. Three of these also shared data and uploaded their code on BitBucket, OSF or referred to it as R-syntax, all three of which were not identified by the algorithm. The fourth did not share data and was missed because it had made its code available as a supplement: "Our methods were easy to implement in R, and the code is presented in Supplementary Table 1, http://links.lww.com/MD/B200." This final mistake had a major impact on the estimated sensitivity because the majority of articles did not share data (5253/6017) and as such this one article was dramatically overweighted. Assuming similar proportions across all estimated 4825 research articles, this algorithm has an accuracy of 98.3% (95% CI, 96.0-99.6%), a sensitivity of 58.7% (95% CI, 34.0-93.7%) and a specificity of 99.7% (95% CI, 99.6-99.9%). Applying this algorithm across the whole PMC is likely to underestimate the true proportion of code sharing by an absolute value of 1.1% (i.e. for every 4825 random PMC research articles of 2015-2019, we expect that this algorithm will label 110 as positive, whereas 164 are actually positive).

#### Conflicts of interest

Our aim was to develop and evaluate a conflict of interest (COI) disclosure identification algorithm. The algorithm was developed in a set of 500 randomly identified articles from PubMed (2015-2018) and evaluated in a set of 6017 randomly identified articles from PubMed Central (2015-2019). Specifically, we were interested in the expected sensitivity, specificity, accuracy and AUROC of this algorithm for various definitions of COI disclosures found on PubMed Central.

We understand COI as arising "whenever activities or relationships compromise the loyalty or independent judgment of an individual who is obligated to serve a party or perform certain roles." (Rodwin, 2017). In building this algorithm, we operationalize this definition by identifying all occasions where authors clearly declare what they call "conflicts of interest" (and synonyms of that, e.g. "Competing interests") and whenever authors clearly report evidence of financial gain (e.g. "J.K. receives fees/stock/benefits from GSK" or "J.K. is employed/on the advisory board of/a presenter for GSK"), but not when the financial gain is implied but not clearly stated (e.g. if an author states as their affiliation GSK, but does not clearly state "J.K. is employed by GSK." What authors call "disclosures" or "financial disclosures" were only considered to represent COIs whenever they used language reminiscent of our working definition (e.g. "Financial disclosures: no competing interests" or "Financial disclosures: J.K. is a consultant for GSK" or "Financial disclosures: Nothing to disclose.") and not otherwise (e.g. "Financial disclosures: J.K. received a grant by the NIH."). We recognize that there are many types of interests (for example, semantically "conflicts of interest" is not identical to "competing financial interests"), that commercial financial gain may not always be a conflict of interest, that reporting a conflict of interest does not imply you do not have other unreported conflicts of interest. These are left for future exploration, which we aid by creating an algorithm that, in its output, denotes why each disclosure was considered a COI (e.g. "J.K. is a consultant for GSK" would be labelled as "consultant").

We explored the performance of our algorithm across three definitions of performance: (a) performance across any article on PubMed Central, (b) performance across non-English articles, (c) performance across articles in which the PDF to text conversion was successful, (d) performance across articles with explicit vs non-explicit disclosures of COI (e.g. "The authors declare no conflicts of interest" vs "J.K. receives benefits from GSK."), (e) performance across research articles vs non-research articles.

***Algorithm development***

The 500 randomly identified articles from PubMed (2015-2018) were split into a train, validation and test set (7:1.5:1.5). We first developed an algorithm on the basis of the train and tested in the validation set. We then improved the algorithm on the basis of the validation set and tested in the test set. We then improved the algorithm on the basis of the test set and tested in the 6017 PubMed Central articles. This structure allowed us to appreciate the performance of our algorithm during development to understand whether the approach we were following was appropriate and whether the algorithm was likely to perform well in the final test set; by using pre-randomized dataset cuts, we avoided biases inherent to stopping rules.

Upon testing in the 6017 PubMed Central articles, we understood that our algorithm was significantly underperforming in recognizing COI disclosures using non-standard language (e.g. "J.K. received benefits from GSK."). Such disclosures were not encountered during training, which is why the algorithm was not built to capture them – upon rescreening the articles, we identified that such disclosures had been made on some occasions, but not identified by the screening team.

In testing within the 6017 articles, the initial algorithm identified 1225 articles that it predicted did not share any COIs and 4792 that did. We took the 1225 articles and split them again into 3 groups: a train group (500, including 180 in which we tested the initial algorithm), a validation group (500) and a test group (225). The test group was only used when algorithm development was complete and this is the performance that we report here – the algorithm has now been modified in view of the mistakes seen in this test set, but not re-evaluated, so we understand this performance as the minimum possible in the target population.

***Algorithm assessment***

As indicated above, the initial algorithm was used to test 6017 articles. We then went through a sub-sample of 100 articles predicted to share a COI disclosure and 180 articles predicted to not share a COI disclosure. These numbers were calculated on the basis of the initial assessment, which predicted no false positives and 3 false negatives.

As indicated above, a second algorithm was developed to cater for non-standard COI disclosures. This was trained and assessed in the 1225 articles that the first algorithm predicted negative (trained in 1000, tested in 225). We then went through all 225 test articles to evaluate the performance of the algorithm.

***Initial development***

The train had 356 articles, 339 of which were in English (this is quite different from the PubMed Central, which appears to have about 3 non-English articles per 200). The algorithm had 100% sensitivity and specificity by design in this set. Of 78 articles in the validation set, 70 were in English. There was only 1 false negative (FN) out of 41 known positives because of an unsuccessful conversion from PDF to text. The test set had 78 articles, 76 of which in English. In this set, the algorithm had 100% specificity. Of the 55 known positives, there were 3 FNs: one because of an unsuccessful conversion to text, one because of a spelling mistake in the text ("confict" instead of "conflict") and one because the algorithm did not identify a true disclosure of COI, even though it should have. In the process, the algorithm also identified 8 occasions where the reviewers failed to identify the COI disclosure (the reviewers had 7 FNs and 1 FP).

***Initial evaluation***

Given that there were 0 false positives (FPs) and 1/55 FNs in the test set during initial algorithm development and that we want to have an estimate within a margin of error of 2% (e.g. 97%, 95-99%), using the formula sqrt((1/55) * (54/55)/180) * 2 = 0.02, we used 100 articles that the algorithm predicted positive and 180 articles predicted negative.

In the test set of 100 FPs, all 100 were true positives (TPs). Of the 180 predicted negative, 136 were deemed research articles and 4 were non-English articles (2 French, 1 Italian, 1 Chinese); none of the non-English articles was non-research. Out of 180 predicted negative, there were 172 TN and 8 FN: 2 were in French, 1 was an unsuccessful conversion from PDF to text (if this were successful, the algorithm would have labelled it correctly) and 5 did not use the standard language observed in the sample of 500 articles on the basis of which this algorithm had been developed; 4/5 of these articles declared conflicts with the industry and 1/5 declared that their study was not commercially sponsored. The following table presents performance across definitions:

| Definition | Sensitivity | Specificity | Accuracy |
|---|---|---|---|
| **Any disclosures** | 97.5% | 100% | 98% |
| **English disclosures** | 97.8% | 100% | 98.2% |
| **Well-converted English disclosures** | 97.9% | 100% | 98.3% |

| | | | |
|---|---|---|---|
| **Explicit disclosures** | 99.6% | - | - |
| **Non-explicit disclosures** | 0.0% | - | - |
| **Any disclosures in research** | 97.6% | 100% | 98.0% |
| **Any disclosures in non-research** | 97.2% | 100% | 98.0% |

***Subsequent development***

The train had 500 articles previously predicted to not have a COI disclosure. Using an updated version of the algorithm according to performance in the previous test (the data of which were included in this train), 29 positives were predicted (5.8%). Out of 18 that had not been seen before, 6 were true positives (3 no involvement of funder disclosures, 2 in French, 1 employee of organization disclosure). The remaining 12 were FPs (67%) (6 because of "relationship", 1 because of "connection", 1 because of "employee", 1 because of "consulting", 3 unknown). One of the articles that was previously part of the test was also a FP. There were 8/146 assessed predicted negative that were FN (5%).

The algorithm was improved by creating a module designed to look for all of these non-explicit disclosures of conflict, as well as disclosures like “This study was not commercially sponsored.”, within the text typically found between Funding/Acknowledgement sections and the Reference section. It was then re-assessed in the validation set of another 500 articles that were previously deemed negative. Out of 500, 30 (6%) were deemed positive. Out of 30 predicted positive, 26 were TP, but 4 were FP (i.e. 13% of positive are false positive). This is a substantial improvement from the previous algorithm (from 67% to 13%), but worse than the initial algorithm, which had no false positives. 2/4 mistakes occurred because the algorithm was not constrained to the

Acknowledgements part due to non-standard language to denote Acknowledgements. Out of 250 predicted negative that were assessed, 16 were FN (6.4%). This remains within the same range as the FNs in the training set (5%), which suggests that many of the improvements simply overfitted to the train data, rather than meaningfully improving the algorithm (in terms of improving sensitivity).

On the basis of these results, the algorithm was improved to be better restricted to Acknowledgements, identify more words related to non-explicit COI disclosures (e.g. “honoraria”, “advisory board”, “commercial relationship”) and certain commonly seen phrases, such as “The funding sources of this study had no role in study design.” It was then reassessed in the final an unseen test set of 225 previously deemed negative studies.

***Subsequent evaluation***

Out of 225 previously deemed negative, 176 (78.2%) were research articles in English. Of the remaining 49, 3 (1.3%) were not in English (2 Chinese, 1 Spanish) and 46 were not research articles (11.6%). One of the 3 articles not in English (the one in Spanish) had a disclosure for conflicts of interest. For all articles (including non-English articles), there were 7 FN (out of 216 deemed negative, 3.2%) and 1 FP (out of 8 deemed positive, 12.5%). Both improvement in FNs (5%, then 6.5%, now 3.2%) and FPs (67%, then 13%, now 12.5%) was substantial, but the improvement in FPs from the validation to test was negligible. This suggests that, despite the improvements in the algorithm on the basis of its performance in the test set, we were unlikely to see substantial improvement in subsequent validation attempts. This is the assessment of the updated algorithm across definitions of COI:

| Definition | Sensitivity | Specificity | Accuracy |
|---|---|---|---|
| **Any disclosures** | 99.2% | 100% | 99.3% |

| | | | |
|---|---|---|---|
| **English disclosures** | 99.3% | 100% | 99.4% |
| **Well-converted English disclosures** | 99.4% | 100% | 99.5% |
| **Explicit disclosures** | 99.9% | - | - |
| **Non-explicit disclosures** | 55.6% | - | - |
| **Any disclosures in research** | 99.3% | 99.4% | 99.4% |
| **Any disclosures in non-research** | 97.2% | 100% | 98.0% |

***Current algorithm***

The absolute final algorithm was used to once again assess the very first test (the one with 180 articles), in which it successfully recognized 7 COI disclosures that the reviewer had not previously seen, raising the number of FNs from 11 to 18. In other words, within articles that were predicted negative by the initial algorithm, the final algorithm increased the number of COI disclosures recognized by 64% over the human reviewer.

***Summary***

Upon manual inspection of 100 articles labelled positive for COI, all 100 indeed reported a COI disclosure. Similarly, out of 225 labelled negative, 218 were indeed negative, but 7 were positive. Of these 7, 1 was in Spanish, 1 was an unsuccessful conversion of PDF to text (had it been successful, it would have been identified), and 4 of the remaining 5 used non-standard language to describe COIs (e.g. "No benefits in any form have been received or will be received from a commercial party."). Running the COI algorithm within the sample of 499 articles from PubMed, we identified 7 articles that had previously been missed by the two reviewers. Assuming similar proportions across all 6017 articles, our algorithm has an accuracy of 99.3% (95% CI, 98.8-99.7%), a sensitivity of 99.2% (95% CI, 98.6-99.7%) and a specificity of 99.5% (95% CI, 98.5-100.0%). Applying this algorithm across the whole PMCOA is likely to underestimate the true proportion of COIs by an absolute value of 0.5% (i.e. for every 6017 random PMCOA articles, we expect this algorithm to label 4840 vs 4873 positive).

#### Funding

Our aim was to develop and evaluate a funding disclosure identification algorithm. The algorithm was developed in a set of 500 randomly identified articles from PubMed (2015-2018) and evaluated in a set of 6017 randomly identified articles from PubMed Central (2015-2019). Specifically, we were interested in the expected sensitivity, specificity, accuracy and AUROC of this algorithm for various definitions of funding disclosures found on PubMed Central.

##### *Funding definitions*

We understand a funding disclosure as an explicit declaration of all sources from which funds were used in completing a study. In building this algorithm, we operationalized this definition by identifying any explicit mentions of funding or financial support. Some of these mentions were found within their own paragraphs, some within Acknowledgements, some within footnotes, some at the very start of a publication. We purposefully avoided mentions of financial relationships or financial disclosures more akin to a COI disclosure, e.g. "Financial disclosures: Nothing to disclose." This is a very unclear statement to us. We purposefully also avoided statements such as: "Information on authorship, contributions, and financial & other disclosures was provided by the authors and is available with the online version of this article at www.haematologica.org." We also understood mentions such as "J.F. has received grant money from GSK" as COI disclosures and classified them like so. Our algorithm does not guarantee that the funding disclosure is true or that it is complete. Ambiguous statements like "We received support from XXX" were interpreted as a funding disclosure from some kind of foundation (e.g. NIH) and not as a funding disclosure if XXX was not a foundation (e.g. Mark Johnson).

We explored the performance of our algorithm across a few definitions of performance: (a) performance across any article on PubMed Central, (b) performance across non-English articles, (c) performance across articles in which the PDF to text conversion was successful, (d) performance across articles with explicit vs non-explicit disclosures of funding (e.g. “Funding: The authors received funding from NIH” vs “Acknowledgements: The authors thank the Wellcome Trust for its support.”), (e) performance across research articles vs non-research articles.

***Algorithm development***

The 500 randomly identified articles from PubMed (2015-2018) were split into a train, validation and test set (7:1.5:1.5). We first developed an algorithm on the basis of the train and tested in the validation set. We then improved the algorithm on the basis of the validation set and tested it in the test set. We then improved the algorithm on the basis of the test set and tested in the 6017 PubMed Central articles. This structure allowed us to appreciate the performance of our algorithm during development to understand whether the approach we were following was appropriate and whether the algorithm was likely to perform well in the final test set; by using pre-randomized dataset cuts, we avoided biases inherent to stopping rules.

Upon testing in the 6017 PubMed Central articles, we understood that our algorithm was significantly underperforming in recognizing funding disclosures using non-standard language (e.g. “The authors thank the Wellcome Trust for its support.”). Such disclosures were not encountered during training, which is why the algorithm was not built to capture them.

In testing within the 6017 articles, the initial algorithm identified 995 articles that it predicted do not share any funding disclosures and 5022 that did. We took the 995 articles, saved 225 as a test and then tested the algorithm in batches of 50. Every time, we tested in the next 50, then developed the algorithm to correctly identify those 50 and then tested in the text 50, and so on. The rule was to stop when performance appears to remain stable across batches of 50. The test group was only touched when algorithm development was complete and this is the performance that we report – note that the algorithm has now been modified in view of the mistakes seen in this test set, but not re-evaluated, so we understand this performance as the minimum possible in the target population.

***Algorithm assessment***

As indicated above, the initial algorithm was used to test 6017 articles. We then went through a sub-sample of 100 articles predicted to share a funding disclosure and 100 articles predicted to not share a funding disclosure. These numbers were arbitrary as we knew that we would be redeveloping the algorithm – we merely wanted an informally representative enough sample based on our experience while developing.

As indicated above, a second algorithm was developed to cater for non-standard funding disclosures. This was trained and assessed in the 995 articles that the first algorithm predicted negative (trained in 769, tested in 226). We then went through all 226 test articles to evaluate the performance of the algorithm.

***Initial development***

The train had 356 articles, 339 of which were in English (this is quite different from the PubMed Central, which appears to have about 3 non-English articles per 200). The algorithm had 100% sensitivity and specificity by design in this set. Of 77 articles in the validation set, there were 5 false negatives (FN) out of 51 known positives and 1 false positive (FP) out of 24 known negatives (Accuracy, 92.2%). Of 78 articles in the test set, there were 2 FNs out of 53 known positives and 4 FPs out of 25 known negatives (Accuracy, 92.3%). In the process, the algorithm also identified 9 occasions where the reviewers made an error (7 FN, 2 FP).

***Initial evaluation***

In the test set of 100 FPs, all 100 were true positives (TPs). Of the 100 predicted negative, 50 were deemed research articles (i.e. provided empirical data or a new method) and 2 were non-English articles (1 French, 1 Chinese); none of the non-English articles was non-research. Out of 100 predicted negative, there were 88 TN and 12 FN, 1 of which errors was due to an unsuccessful conversion from PDF to text (if this were successful, the algorithm would have labelled it correctly) and 8 did not use the standard language observed in the sample of 500 articles on the basis of which this algorithm had been developed; 9 of the missed articles declared some kind of funding and 3 declared no funding. The following table presents performance across definitions:

| Definition | Sensitivity | Specificity | Accuracy |
|---|---|---|---|
| **Any disclosures** | 97.7% | 100% | 98.0% |
| **English disclosures** | 97.7% | 100% | 98.0% |
| **Well-converted English disclosures** | 97.9% | 100% | 98.2% |
| **Explicit disclosures** | 99.2% | - | - |
| **Non-explicit disclosures** | 00.0% | - | - |
| **Any disclosures in research** | 97.9% | 100% | 98.1% |
| **Any disclosures in non-research** | 94.6% | 100% | 97.3% |

***Subsequent development***

Development proceeded to consider 450 articles in detail and another 319 for just false positives (i.e. only the articles that the algorithm said were positive were examined). The algorithm was improved with more types of funding titles and by trying to capture more non-explicit mentions of funding. Accuracy for successfully converted PDFs changed like so: 86% (1:50), 92% (51:100), 98% (101:150), 90% (151:200), 96% (201:250), 88% (251:300), 96% (201:350), 90% (351:400), 98% (401:450) and no false positives in 451:769 – the mean accuracy was 93.7%, excluding the first 100.

***Subsequent evaluation***

Out of 226 previously deemed negative, 116 (51.3%) were research articles in English. Of the remaining 110, 1 (0.9%) was not in English (1 French) and did not have a funding disclosure. For all articles (including non-English articles), there were 4 FN (out of 211 predicted negative, 1.9%) and 4 FP (out of 15 predicted positive, 26.7%). As such, there was a substantial decrease in FNs (from 12% to 1.9%), but also, within this set a substantial increase in FPs (from 0% to 26.7%) – had these articles been assessed using the old algorithm, the accuracy would have dropped from 96.5% (218/226) to 93.4% (211/226). This suggested that the algorithm was better able to now identify non-standard mentions of funding, but at the cost of a loss in specificity.

| Definition | Sensitivity | Specificity | Accuracy |
|---|---|---|---|
| **Any disclosures** | 99.7% | 98.1% | 99.4% |
| **English disclosures** | 99.7% | 98.1% | 99.4% |
| **Well-converted English disclosures** | 99.7% | 98.1% | 99.5% |
| **Explicit disclosures** | 99.7% | - | - |

| | | | |
|---|---|---|---|
| **Non-explicit disclosures** | 100% | - | - |
| **Any disclosures in research** | 99.7% | 99.0% | 99.7% |
| **Any disclosures in non-research** | 100% | 97.2% | 98.4% |

Had we been using the old algorithm in this test set, the performance would have been: 98.7% sensitivity, 100% specificity, 100% PPV, 93.4% NPV and 98.9% accuracy for any disclosure.

***Current algorithm***

The final algorithm was used to once again assess the very first test (the one with 100 articles), in which no more funding disclosures were recognized. This suggests that human performance is superior to machine performance. It should also be noted that adjudication of the 226 occurred before learning the response of the algorithm, which clearly indicates that human performance is superior.

***Summary***

Upon manual inspection of 100 articles labelled positive for Funding, all 100 indeed had an explicit Funding disclosure. The algorithm was then calibrated to PMC articles that were initially labelled negative and then tested in a random unseen sample. Of 226 test articles initially labelled negative, the algorithm correctly predicted 218. Of the remaining 8, 4 were falsely predicted negative (FN) and 4 false predicted positive (FP). Of 4 FNs, 1 was an unsuccessful conversion of PDF to text (had it been successful, it would have been predicted positive) and 3 used uncommon language (e.g. “We would like to thank the U.S. Embassy in Addis Ababa, Ethiopia, Jigjiga University, and Texas Tech University for funding this research”). Of 4 FPs, 2 referred to funds received by another study and 2 contained words commonly seen in funding disclosures ("support" and "scholar"). Running the Funding algorithm within the sample of 520 articles from PubMed, we identified 7 articles previously mislabelled negative and 2 previously mislabelled positive. Assuming similar proportions across all 6017 articles, our algorithm has an accuracy of 99.4% (95% CI, 99.0-99.8%), a sensitivity of 99.7% (95% CI, 99.3-99.9%) and a specificity of 98.1% (95% CI, 96.1-99.5%). Applying this algorithm across the whole PMCOA is expected to identify the true proportion of funding disclosures (i.e. for every 6017 random PMCOA articles, we expect this algorithm to label 5084 vs 5084 positive). This algorithm can be improved in the future by (a) using a probabilistic model (e.g. a random forest) to predict outcome on the basis of the exported features and (b) by improving the quality of the extracted text from the PDFs.

#### Protocol registration

Our aim was to develop and evaluate a registration statement identification algorithm. The algorithm was developed in a set of 500 randomly identified articles from PubMed (2015-2018) and evaluated in a set of 6017 randomly identified articles from PubMed Central (2015-2019). Specifically, we were interested in the expected sensitivity, specificity, accuracy and AUROC of this algorithm for various definitions of registration statements found on PubMed Central.

##### *Registration definitions*

We understand a registration statement as any explicit declaration of study registration. This can include registrations on known websites, such as registration on ClinicalTrials.gov, as well as a previous publication of the protocol, as long as these registrations/protocols refer to the current study. Registered protocols that are not publicly available, such as statements like “The study was approved by the Ruakura Animal Ethics Committee (protocol no. 13899)” or “Alliance for Clinical Trials in Oncology (formerly Cancer and Leukemia Group B) Protocol #369901”, were not considered as protocol registration statements because they do not enhance transparency. Even though the algorithm was set up to also identify mentions such as “This study was not registered on any registry”, this ability is turned off by default because this statement does not enhance transparency in any way (i.e. it is not informative).

We explored the performance of our algorithm across a few definitions of performance: (a) performance across any article on PubMed Central, (b) performance across articles with explicit vs non-explicit statements of registration (e.g. “This study was registered on ClinicalTrials.gov with registration number NCT01234567”), (c) performance across research articles vs non-research articles and (d) performance within articles that mention NCT.

Note that by explicit we refer to any mentions of registration as a title, e.g. “Trial Registration: NCT…” or “Trial Number: NCT…” but not “Clinical Trial information: NCT…”, as clear registration statement, e.g. “Clinical Trial Registration: NCT…”, or as a clear phrase, e.g. “This study was registered on PROPSERO with ID number (…)” – for a phrase to be clear, it had to mention “registration” somewhere, in addition with the ID.

***Algorithm development***

The 500 randomly identified articles from PubMed (2015-2018) were split into a train, validation and test set (7:1.5:1.5). We first developed an algorithm on the basis of the train and tested in the validation set. We then improved the algorithm on the basis of the validation set and tested it in the test set. We then improved the algorithm on the basis of the test set and tested in the 6017 PubMed Central articles. This structure allowed us to appreciate the performance of our algorithm during development to understand whether the approach we were following was appropriate and whether the algorithm was likely to perform well in the final test set; by using pre-randomized dataset cuts, we avoided biases inherent to stopping rules.

Upon testing in the 6017 PubMed Central articles, we understood that our algorithm was significantly underperforming in terms of false positives because of identifying mentions within references, mentions of NCT registrations within the introduction, mentions of NCT references as “this trial is underway” or in referring to the data being used, e.g. “We are using data from this trial (NCT01234567)”. This is why we redeveloped the algorithm in the first 100 articles and then tested the algorithm in the next 161 articles, out of 261 deemed positive by the initial algorithm.

In terms of false negatives, the initial algorithm had excellent performance. However, the re-developed algorithm falsely identified mentions of “registry” as evidence of a registration statement, many of which were not. This was fixed, for which reason we expect on average the reported test performance to be an underestimation.

##### *Algorithm assessment*

As indicated above, the initial algorithm was used to test 6017 articles. We then went through 100 articles of predicted positives to appreciate algorithm performance and then assessed both the initial and redeveloped algorithm in the remaining 161 positive articles. Out of 5756 predicted negative, we used a stratified/importance sampling procedure by sampling 10/3248 articles deemed irrelevant (i.e. do not mention the words regist*/trial/NCT), 20/452 that were deemed relevant, 55/1962 out of those that were deemed relevant and had a Methods sections, 61/94 that contained an NCT identification number.

##### *Initial development*

The train had 356 articles, 339 of which were in English. Developing this algorithm was not trivial because we had to compromise in what phrases were identified to balance sensitivity with specificity; given the very small number of positive examples in our train set, this was challenging. After initial development, the algorithm ran in 7 seconds in the train set with 4 FN (one of which because the line was inappropriately split) and 0 FPs (98.9% accuracy). The algorithm was corrected on the basis of these and was then re-tested in the 77 cases of the validation set, where there were eventually 76 TNs and 1 FN (the algorithm did not identify PROSPERO registration) (98.7% accuracy). This was corrected and the algorithm was then re-tested in the 78 records of the test set, where there were 73 TNs and 5 FNs (93.6% accuracy).

Eventually, using the latest iteration of the algorithm, we discovered another 3 TPs that had been missed by the reviewers and 1 TN, that had been labelled as positive by the reviewers.

***Initial evaluation***

In the test set of 261 articles predicted positive, we randomly identified 161 as a test set. Of the 161 predicted positive, there were 147 TPs and 14 FPs (i.e. there were 14 errors). Most mistakes occurred because of articles referring to clinical trials underway, using the data of previously completed clinical trials or registration of their protocol with an IRB committee. Of the articles predicted negative, the following performance was observed (non-relevant articles, 21/21 TNs; relevant articles with no methods or NCT, 10/10 TN; relevant articles with methods and no NCT, 53/55 TN and 2/55 FN; articles referring to NCT, 55/61 TNs, 6/61 FNs). The following table presents performance across definitions when adjusted for the weighted sampling:

| Definition | Sensitivity | Specificity | Accuracy |
|---|---|---|---|
| **Any statements** | 97.3% | 99.6% | 99.5% |
| **Explicit statements** | 99.6% | - | - |
| **Non-explicit statements** | 69.3% | - | - |
| **Any statements in research** | 97.9% | 99.5% | 99.5% |
| **Any statements in non-research** | 91.9% | 100% | 99.8% |

***Subsequent development***

Subsequent development occurred in 100/261 articles predicted positive. Given the small sample size, no assessment across collections of 50 articles was done. Additionally, no development in the articles predicted negative was done. Overall, given the performance in the test set, the development did not particularly improve performance, even though it substantially improved ability to identify non-explicit statements.

***Subsequent evaluation***

Out of 161 articles previously predicted positive, the new algorithm identified 21 N and 140 P. Of these there were 147 TP, 4 FP, 10 TN, 5 FN (i.e. there were 9/161 errors). Of the 5756 articles predicted negative, the new algorithm predicted that 15 were in fact positive – of those there were 4 TP and 11 FPs. This suggests that the new algorithm is better in terms of specificity at the expense of sensitivity.

| Definition | Sensitivity | Specificity | Accuracy |
|---|---|---|---|
| **Any statements** | 95.6% | 99.7% | 99.5% |
| **Explicit statements** | 96.4% | - | - |
| **Non-explicit statements** | 85.4% | - | - |
| **Any statements in research** | 95.1% | 99.7% | 99.5% |
| **Any statements in non-research** | 100% | 99.8% | 99.8% |

***Current algorithm***

The final algorithm was used to once again assess the very first test, identifying 3 previously non-identified statements as indicated above.

***Summary***

Upon manual inspection of 161/261 articles labelled positive for a protocol registration statement, we found 5 FNs and 4 FPs. Of 5 FNs, 4 were grammatical failures of the algorithm to understand that the registration statement referred to the current and not some other study (e.g. "This registered study on www.clinicaltrials.gov (NCT01375270) was approved ...") and 1 was a statement contained within financial disclosures. Of 4 FPs, 2 were mentions of registration in the references, 1 referred to the registration of a study of which the data it was using and 1 was using a registered study as an example. Similarly, of 147/5657 articles initially labelled negative, there were 11 FPs and 2 FNs - note that the large number of errors occurred because of sampling from articles in which the algorithm was more likely to underperform (see Methods). Of 11 FPs, most errors occurred because of registrations that did not refer to open protocol registrations (e.g. approval by a medical ethics committee) and because of referral to other registries (e.g. a patient registry). Of the 2 FNs, 1 was a grammatical failure of the algorithm to understand that the registration statement referred to this study and 1 did not mention anything about registration, other than the NCT number (“EDITION 2 (NCT01499095) was a randomized, 6-month, multicentre, open-label, two-arm, phase IIIa study investigating ….”). Running the Registration algorithm within the sample of 499 articles from PubMed, we identified  2 positive and 1 negative studies that were previously erroneously labelled. Assuming similar proportions across all 6017 articles, our algorithm has an accuracy of 99.5% (95% CI, 99.3-99.7%), a sensitivity of 95.6% (95% CI, 92.0-98.6%) and a specificity of 99.7% (95% CI, 99.5-99.8%). Applying this algorithm across the whole PMCOA is likely to underestimate the true proportion of protocol registration statements by an absolute value of -0.14% (i.e. for every 6017 random PMCOA articles, we expect this algorithm to label 249 vs 241 positive).

#### S3 Text. Associations of transparency with article characteristics.

Considering only publications from 2000 onwards, publications with a Conflict of Interest (COI) disclosure had substantially more references (Median, 38 vs 22), more often represented research articles (Proportion, 90.0% vs 85.3%) and tended to be published in journals of lower impact factor (Median, 2.8 vs 3.4) (Supporting Information Table 5). Publications with funding disclosures tended to be primarily found in research articles (Proportion, 91.92% vs 78.7%) and have a median of more references (40 vs 18), more affiliations (3 vs 2), more authors (6 vs 4) and more citations (6 vs 4). Publications with registration statements tended to have a median of more affiliations (4 vs 3), more authors (7 vs 5) and more tables (3 vs 1).

#### S4 Text. Performance metrics.

We hereby define the performance metrics shown in Figure 2 and illustrate how they were calculated. The code for this procedure has been made available (see Code Availability). The performance metrics and their calculation are illustrated by using the test data for COI disclosures. The initial COI disclosures algorithm predicted that out of 6,017 random articles from PMC, 1,225 do not share COI disclosures and 4,792 do.

First, we took a random subsample of 100/4,792 articles predicted positive (i.e. a COI disclosure was found). We found:

| | | **Manual assessment** | | |
|---|---|---|---|---|
| | | **True** | **False** | |
| **Automated assessment** | **True** | True positive (TP) = 100 | False positive (FP) = 0 | PPV = 100 / (100+0) = 100% |
| | **False** | False negative (FN) = 0 | True negative (TN) = 0 | NPV = Not applicable |
| | | Sensitivity = 100 / (100+0) = 100% | Specificity = Not applicable | Accuracy = 100 / 100 = 100% |

*PPV = Positive Predictive Value (Precision). NPV = Negative Predictive Value.*

Prevalence (true) = (TP+FN) / (TP+FP+FN+TN) = 100/100 = 100%

Prevalence (estimated) = (TP+FP) / (TP+FP+FN+TN) = 100/100 = 100%

Error = Prevalence (true) - Prevalence (estimated) = 100% - 100% = 0%

Second, we took a random subsample of 225/1,225 initially predicted negative (i.e. a COI disclosure was not found). We found:

| | | Manual assessment | | |
|---|---|---|---|---|
| | | **True** | **False** | |
| **Automated assessment** | **True** | True positive (TP) = 8 | False positive (FP) = 1 | PPV = 8 / (8+1) = 89% |
| | **False** | False negative (FN) = 7 | True negative (TN) = 209 | NPV = 209 / (7+209) = 97% |
| | | Sensitivity = 8 / (8+7) = 53% | Specificity = 209 / (1+209) = 100% | Accuracy = (8+209) / 225 = 96% |

PPV = Positive Predictive Value (Precision). NPV = Negative Predictive Value.

Prevalence (true) = (TP+FN) / (TP+FP+FN+TN) = 15/225 = 7%

Prevalence (estimated) = (TP+FP) / (TP+FP+FN+TN) = 9/225 = 4%

Error = Prevalence (true) - Prevalence (estimated) = 7% - 4% = 3%

Finally, we weighted the two tables into the statistics presented in Figure 2. This may be done by using the law of total probability with Bayes' theorem. However, the same end-result may be achieved by using the aforementioned tables to estimate the expected table, had we manually assessed all 6017 articles:

| | Manual assessment | |
|---|---|---|
| | **True** | **False** |

| | | | | |
|---|---|---|---|---|
| **Automated assessment** | **True** | TP =<br>100/100 * 4792 +<br>8/225 * 1225 = 4836 | FP =<br>0/100 * 4792 +<br>1/22 * 1225 = 56 | PPV =<br>4836 / (4836+56) = 99% |
| | **False** | FN =<br>0/100 * 4792 +<br>7/225 * 1225 = 38 | TN =<br>0/100 * 4792 +<br>209/225 * 1225 = 1138 | NPV =<br>1138 / (1138+38) = 97% |
| | | Sensitivity =<br>4836 / (4836+38) = 99% | Specificity =<br>1138 / (1138+56) = 95% | Accuracy =<br>(4836+1138) / 6017 = 99% |

*PPV = Positive Predictive Value (Precision). NPV = Negative Predictive Value.*

Prevalence (true) = (TP+FN) / (TP+FP+FN+TN) = (4836+38) / 6017 = 81%

Prevalence (estimated) = (TP+FP) / (TP+FP+FN+TN) = (4836+56) / 6017 = 81%

Error = Prevalence (true) - Prevalence (estimated) = 81% - 81% = 0%
