## Supplemental Table 3 for "Assessment of transparency indicators across the biomedical literature: how open is open?": s3-table_meta-data-desc.html

Data Frame Summary


### Data Frame Summary

#### Meta-data

**Dimensions**: 2751420 x 40
  
**Duplicates**: 0
  

| **No** | **Variable** | **Stats / Values** | **Freqs (% of Valid)** | **Graph** | **Valid** | **Missing** |
| --- | --- | --- | --- | --- | --- | --- |
| 1 | PMCID [character] | 1. 100320 2. 100321 3. 100322 4. 100323 5. 100324 6. 100325 7. 100326 8. 100327 9. 100357 10. 100780 [ 2751410 others ] | 1 \| ( \| 0.0% \| ); 1 \| ( \| 0.0% \| ); 1 \| ( \| 0.0% \| ); 1 \| ( \| 0.0% \| ); 1 \| ( \| 0.0% \| ); 1 \| ( \| 0.0% \| ); 1 \| ( \| 0.0% \| ); 1 \| ( \| 0.0% \| ); 1 \| ( \| 0.0% \| ); 1 \| ( \| 0.0% \| ); 2751410 \| ( \| 100.0% \| ) |  | 2751420 (100%) | 0 (0%) |
| 2 | PMID [character] | 1. 1001287 2. 1001288 3. 1001289 4. 1001290 5. 1001291 6. 1001292 7. 1001293 8. 1001294 9. 1001295 10. 1001296 [ 2591839 others ] | 1 \| ( \| 0.0% \| ); 1 \| ( \| 0.0% \| ); 1 \| ( \| 0.0% \| ); 1 \| ( \| 0.0% \| ); 1 \| ( \| 0.0% \| ); 1 \| ( \| 0.0% \| ); 1 \| ( \| 0.0% \| ); 1 \| ( \| 0.0% \| ); 1 \| ( \| 0.0% \| ); 1 \| ( \| 0.0% \| ); 2591839 \| ( \| 100.0% \| ) |  | 2591849 (94.2%) | 159571 (5.8%) |
| 3 | DOI [character] | 1. 10.4103/1742-6413.84993 2. 10.1102/1470-7330.2001.00 3. 10.1102/1470-7330.2005.00 4. 10.1186/s40348-014-0002-2 5. 10.15537/smj.2019.1.23890 6. 10.1590/1678-775720150285 7. 10.1590/abd1806-4841.2014 8. 10.2106/JBJS.ST.18.00083 9. 10.30699/IJP.2019.77233.1 10. 10.31372/20190404.1078 [ 2439609 others ] | 16 \| ( \| 0.0% \| ); 2 \| ( \| 0.0% \| ); 2 \| ( \| 0.0% \| ); 2 \| ( \| 0.0% \| ); 2 \| ( \| 0.0% \| ); 2 \| ( \| 0.0% \| ); 2 \| ( \| 0.0% \| ); 2 \| ( \| 0.0% \| ); 2 \| ( \| 0.0% \| ); 2 \| ( \| 0.0% \| ); 2439611 \| ( \| 100.0% \| ) |  | 2439645 (88.67%) | 311775 (11.33%) |
| 4 | PII [character] | 1. sc271\_5\_1835 2. Doc01 3. Doc02 4. Doc03 5. Doc05 6. Doc06 7. Doc04 8. Doc07 9. Doc09 10. Doc08 [ 130675 others ] | 1523 \| ( \| 1.1% \| ); 26 \| ( \| 0.0% \| ); 25 \| ( \| 0.0% \| ); 25 \| ( \| 0.0% \| ); 25 \| ( \| 0.0% \| ); 25 \| ( \| 0.0% \| ); 23 \| ( \| 0.0% \| ); 23 \| ( \| 0.0% \| ); 20 \| ( \| 0.0% \| ); 19 \| ( \| 0.0% \| ); 131041 \| ( \| 98.7% \| ) |  | 132775 (4.83%) | 2618645 (95.17%) |
| 5 | Title [character] | 1. Book Reviews 2. Annotations 3. Issue Information 4. Current Topics 5. Notes and News 6. Medical News 7. Erratum 8. Book Review 9. Surgery 10. Medicine [ 2679666 others ] | 3039 \| ( \| 0.1% \| ); 1291 \| ( \| 0.0% \| ); 1206 \| ( \| 0.0% \| ); 1149 \| ( \| 0.0% \| ); 1023 \| ( \| 0.0% \| ); 897 \| ( \| 0.0% \| ); 856 \| ( \| 0.0% \| ); 854 \| ( \| 0.0% \| ); 846 \| ( \| 0.0% \| ); 845 \| ( \| 0.0% \| ); 2738824 \| ( \| 99.6% \| ) |  | 2750830 (99.98%) | 590 (0.02%) |
| 6 | Date (electronic) [character] | 1. 08-11-2019 2. 30-04-2019 3. 23-10-2019 4. 26-11-2018 5. 04-10-2017 6. 01-10-2015 7. 28-02-2019 8. 30-06-2017 9. 27-11-2019 10. 27-05-2016 [ 7838 others ] | 4845 \| ( \| 0.2% \| ); 3667 \| ( \| 0.2% \| ); 3656 \| ( \| 0.2% \| ); 3378 \| ( \| 0.2% \| ); 2242 \| ( \| 0.1% \| ); 1973 \| ( \| 0.1% \| ); 1709 \| ( \| 0.1% \| ); 1701 \| ( \| 0.1% \| ); 1675 \| ( \| 0.1% \| ); 1665 \| ( \| 0.1% \| ); 2214840 \| ( \| 98.8% \| ) |  | 2241351 (81.46%) | 510069 (18.54%) |
| 7 | Year (electronic) [numeric] | Mean (sd) : 2015.2 (3.5) min < med < max: 1959 < 2016 < 2020 IQR (CV) : 5 (0) | 35 distinct values |  | 2241351 (81.46%) | 510069 (18.54%) |
| 8 | Date (publication) [character] | 1. 2014 2. 2015 3. 2017 4. 2016 5. 2013 6. 2012 7. 2018 8. 2019 9. 2011 10. 2010 [ 13285 others ] | 22903 \| ( \| 2.1% \| ); 22499 \| ( \| 2.0% \| ); 22485 \| ( \| 2.0% \| ); 21443 \| ( \| 1.9% \| ); 19088 \| ( \| 1.7% \| ); 16242 \| ( \| 1.5% \| ); 14467 \| ( \| 1.3% \| ); 11559 \| ( \| 1.0% \| ); 10410 \| ( \| 0.9% \| ); 6179 \| ( \| 0.6% \| ); 934513 \| ( \| 84.8% \| ) |  | 1101788 (40.04%) | 1649632 (59.96%) |
| 9 | Year (publication) [numeric] | Mean (sd) : 1995.6 (43.3) min < med < max: 1781 < 2014 < 2020 IQR (CV) : 11 (0) | 240 distinct values |  | 1101788 (40.04%) | 1649632 (59.96%) |
| 10 | Journal [character] | 1. PLoS ONE 2. Scientific Reports 3. The Hospital 4. The Indian Medical Gazett 5. Oncotarget 6. The Journal of Cell Biolo 7. Nature Communications 8. British Journal of Cancer 9. The Journal of Experiment 10. Acta Crystallographica Se [ 10560 others ] | 229888 \| ( \| 8.4% \| ); 106480 \| ( \| 3.9% \| ); 34263 \| ( \| 1.2% \| ); 29143 \| ( \| 1.1% \| ); 25501 \| ( \| 0.9% \| ); 25041 \| ( \| 0.9% \| ); 24768 \| ( \| 0.9% \| ); 24763 \| ( \| 0.9% \| ); 24099 \| ( \| 0.9% \| ); 23627 \| ( \| 0.9% \| ); 2203847 \| ( \| 80.1% \| ) |  | 2751420 (100%) | 0 (0%) |
| 11 | Journal abbreviation (NLM) [character] | 1. PLoS One 2. Sci Rep 3. Hospital (Lond 1886) 4. Ind Med Gaz 5. Nat Commun 6. Oncotarget 7. J Cell Biol 8. Br J Cancer 9. J Exp Med 10. Acta Crystallogr Sect E S [ 9414 others ] | 229888 \| ( \| 8.4% \| ); 106480 \| ( \| 3.9% \| ); 34263 \| ( \| 1.2% \| ); 29143 \| ( \| 1.1% \| ); 25610 \| ( \| 0.9% \| ); 25501 \| ( \| 0.9% \| ); 25041 \| ( \| 0.9% \| ); 24763 \| ( \| 0.9% \| ); 24099 \| ( \| 0.9% \| ); 23627 \| ( \| 0.9% \| ); 2202921 \| ( \| 80.1% \| ) |  | 2751336 (100%) | 84 (0%) |
| 12 | Journal abbreviation (ISO) [character] | 1. PLoS ONE 2. Sci Rep 3. Nat Commun 4. Oncotarget 5. J. Cell Biol 6. J. Exp. Med 7. Int J Mol Sci 8. Sensors (Basel) 9. Molecules 10. Medicine (Baltimore) [ 9416 others ] | 229888 \| ( \| 10.0% \| ); 106167 \| ( \| 4.6% \| ); 25325 \| ( \| 1.1% \| ); 25307 \| ( \| 1.1% \| ); 24184 \| ( \| 1.1% \| ); 23730 \| ( \| 1.0% \| ); 21335 \| ( \| 0.9% \| ); 21279 \| ( \| 0.9% \| ); 19487 \| ( \| 0.8% \| ); 18794 \| ( \| 0.8% \| ); 1777544 \| ( \| 77.5% \| ) |  | 2293040 (83.34%) | 458380 (16.66%) |
| 13 | ISSN (print) [character] | 1. 2041-1723 2. 1540-8140 3. 1532-1827 4. 1540-9538 5. 1422-0067 6. 1362-4962 7. 1420-3049 8. 1536-5964 9. 2314-6133 10. 1466-609X [ 8926 others ] | 25610 \| ( \| 1.4% \| ); 25041 \| ( \| 1.4% \| ); 24125 \| ( \| 1.4% \| ); 24099 \| ( \| 1.4% \| ); 22912 \| ( \| 1.3% \| ); 19717 \| ( \| 1.1% \| ); 19487 \| ( \| 1.1% \| ); 18853 \| ( \| 1.1% \| ); 17210 \| ( \| 1.0% \| ); 16270 \| ( \| 0.9% \| ); 1569831 \| ( \| 88.0% \| ) |  | 1783155 (64.81%) | 968265 (35.19%) |
| 14 | ISSN (electronic) [character] | 1. 1932-6203 2. 2045-2322 3. 0267-6478 4. 0019-5863 5. 1949-2553 6. 0021-9525 7. 0007-0920 8. 0022-1007 9. 1424-8220 10. 1600-5368 [ 8172 others ] | 229888 \| ( \| 11.0% \| ); 106480 \| ( \| 5.1% \| ); 34263 \| ( \| 1.6% \| ); 29143 \| ( \| 1.4% \| ); 25501 \| ( \| 1.2% \| ); 25041 \| ( \| 1.2% \| ); 24125 \| ( \| 1.2% \| ); 24099 \| ( \| 1.2% \| ); 23158 \| ( \| 1.1% \| ); 22962 \| ( \| 1.1% \| ); 1541060 \| ( \| 73.9% \| ) |  | 2085720 (75.81%) | 665700 (24.19%) |
| 15 | Publisher [character] | 1. BioMed Central 2. Public Library of Science 3. Nature Publishing Group 4. MDPI 5. Frontiers Media S.A. 6. Hindawi Publishing Corpor 7. Medknow Publications 8. Springer 9. Wiley-Blackwell Publishin 10. Oxford University Press [ 1159 others ] | 393489 \| ( \| 14.7% \| ); 271128 \| ( \| 10.1% \| ); 173100 \| ( \| 6.5% \| ); 148981 \| ( \| 5.6% \| ); 130078 \| ( \| 4.8% \| ); 113473 \| ( \| 4.2% \| ); 101474 \| ( \| 3.8% \| ); 100687 \| ( \| 3.8% \| ); 97316 \| ( \| 3.6% \| ); 75085 \| ( \| 2.8% \| ); 1078053 \| ( \| 40.2% \| ) |  | 2682864 (97.51%) | 68556 (2.49%) |
| 16 | Publisher ID [character] | 1. plos 2. Acta Cryst. E 3. ijms 4. Oncotarget; ImpactJ 5. molecules 6. MEDI 7. BMRI 8. sensors 9. nar 10. bmjopen [ 6310 others ] | 263962 \| ( \| 16.3% \| ); 26560 \| ( \| 1.6% \| ); 24687 \| ( \| 1.5% \| ); 20019 \| ( \| 1.2% \| ); 19487 \| ( \| 1.2% \| ); 18804 \| ( \| 1.2% \| ); 17359 \| ( \| 1.1% \| ); 16779 \| ( \| 1.0% \| ); 16740 \| ( \| 1.0% \| ); 16419 \| ( \| 1.0% \| ); 1178965 \| ( \| 72.8% \| ) |  | 1619781 (58.87%) | 1131639 (41.13%) |
| 17 | Authors [character] | 1. LeBrasseur Nicole 2. Leslie Mitch 3. Short Ben 4. Wells William A. 5. Sedwick Caitlin 6. Wiwanitkit Viroj 7. Macgregor A. S. M. 8. Jamieson W. Allan 9. Craig William 10. Robinson Richard [ 2465934 others ] | 633 \| ( \| 0.0% \| ); 497 \| ( \| 0.0% \| ); 464 \| ( \| 0.0% \| ); 434 \| ( \| 0.0% \| ); 261 \| ( \| 0.0% \| ); 255 \| ( \| 0.0% \| ); 245 \| ( \| 0.0% \| ); 243 \| ( \| 0.0% \| ); 228 \| ( \| 0.0% \| ); 212 \| ( \| 0.0% \| ); 2612180 \| ( \| 99.9% \| ) |  | 2615652 (95.07%) | 135768 (4.93%) |
| 18 | Affiliation [character] | 1. From the Laboratories of 2. From the Laboratories of 3. Wellcome Institute 4. Centers for Disease Contr 5. Department of Chemistry, 6. From the Hospital of The 7. Department of Chemistry, 8. Medical Officer of Health 9. From the Hospital of The 10. From The Rockefeller Inst [ 2411684 others ] | 766 \| ( \| 0.0% \| ); 633 \| ( \| 0.0% \| ); 437 \| ( \| 0.0% \| ); 395 \| ( \| 0.0% \| ); 377 \| ( \| 0.0% \| ); 295 \| ( \| 0.0% \| ); 274 \| ( \| 0.0% \| ); 243 \| ( \| 0.0% \| ); 208 \| ( \| 0.0% \| ); 189 \| ( \| 0.0% \| ); 2549698 \| ( \| 99.9% \| ) |  | 2553515 (92.81%) | 197905 (7.19%) |
| 19 | Affiliation country [character] | 1. USA 2. China 3. China; China 4. USA; USA 5. USA; USA; USA 6. UK 7. China; China; China 8. UK; UK 9. South Korea 10. Germany [ 74117 others ] | 22769 \| ( \| 4.7% \| ); 18605 \| ( \| 3.9% \| ); 18301 \| ( \| 3.8% \| ); 16784 \| ( \| 3.5% \| ); 13183 \| ( \| 2.7% \| ); 10719 \| ( \| 2.2% \| ); 10244 \| ( \| 2.1% \| ); 7501 \| ( \| 1.6% \| ); 6950 \| ( \| 1.4% \| ); 6265 \| ( \| 1.3% \| ); 348522 \| ( \| 72.6% \| ) |  | 479843 (17.44%) | 2271577 (82.56%) |
| 20 | Affiliation institute [character] | 1. University of California 2. University of Oxford 3. University of Cambridge 4. University College London 5. University of Pennsylvani 6. University of Manchester 7. Imperial College London 8. University of Michigan 9. Massachusetts Institute o 10. The Wellcome Trust Centre [ 652675 others ] | 299 \| ( \| 0.0% \| ); 295 \| ( \| 0.0% \| ); 249 \| ( \| 0.0% \| ); 211 \| ( \| 0.0% \| ); 144 \| ( \| 0.0% \| ); 141 \| ( \| 0.0% \| ); 138 \| ( \| 0.0% \| ); 132 \| ( \| 0.0% \| ); 126 \| ( \| 0.0% \| ); 122 \| ( \| 0.0% \| ); 708009 \| ( \| 99.7% \| ) |  | 709866 (25.8%) | 2041554 (74.2%) |
| 21 | Author-Affiliation ID [character] | 1. ; 2. ; cor1 3. cor1; 4. aff1, cor1 5. I1; I1 6. I1 7. Aff1; Aff1 8. I1; I1; I1 9. Aff1; Aff1; Aff1 10. Aff1 [ 1208099 others ] | 105707 \| ( \| 4.2% \| ); 16724 \| ( \| 0.7% \| ); 13521 \| ( \| 0.5% \| ); 12284 \| ( \| 0.5% \| ); 11774 \| ( \| 0.5% \| ); 11649 \| ( \| 0.5% \| ); 11012 \| ( \| 0.4% \| ); 10634 \| ( \| 0.4% \| ); 10301 \| ( \| 0.4% \| ); 8784 \| ( \| 0.3% \| ); 2302632 \| ( \| 91.6% \| ) |  | 2515022 (91.41%) | 236398 (8.59%) |
| 22 | Affiliation ID [character] | 1. aff1; aff2 2. NA 3. aff1 4. aff1; aff2; aff3 5. I1 6. I1; I2 7. Aff1; Aff2 8. Aff1; Aff2; Aff3 9. I1; I2; I3 10. aff1; aff2; aff3; aff4 [ 438673 others ] | 134976 \| ( \| 5.3% \| ); 116548 \| ( \| 4.6% \| ); 113332 \| ( \| 4.5% \| ); 102647 \| ( \| 4.0% \| ); 84149 \| ( \| 3.3% \| ); 83929 \| ( \| 3.3% \| ); 76315 \| ( \| 3.0% \| ); 70477 \| ( \| 2.8% \| ); 66963 \| ( \| 2.6% \| ); 66171 \| ( \| 2.6% \| ); 1627207 \| ( \| 64.0% \| ) |  | 2542714 (92.41%) | 208706 (7.59%) |
| 23 | Correspondence [character] Emails | Valid Invalid Duplicates | 1727526 \| ( \| 98.7% \| ); 23137 \| ( \| 1.3% \| ); 211564 \| ( \| 12.1% \| ) |  | 1750663 (63.63%) | 1000757 (36.37%) |
| 24 | License [character] | 1. This is an open-access ar 2. Open AccessThis article i 3. This is an open access ar 4. Licensee MDPI, Basel, Swi 5. This is an open access ar 6. This is an Open Access ar 7. Open Access This article 8. http://creativecommons.or 9. This is an open access ar 10. This is an open-access ar [ 48648 others ] | 167957 \| ( \| 6.5% \| ); 143222 \| ( \| 5.5% \| ); 121133 \| ( \| 4.7% \| ); 106487 \| ( \| 4.1% \| ); 97020 \| ( \| 3.8% \| ); 86295 \| ( \| 3.3% \| ); 83017 \| ( \| 3.2% \| ); 72054 \| ( \| 2.8% \| ); 60044 \| ( \| 2.3% \| ); 58149 \| ( \| 2.3% \| ); 1585827 \| ( \| 61.4% \| ) |  | 2581205 (93.81%) | 170215 (6.19%) |
| 25 | Type of publication [character] | 1. research-article 2. review-article 3. case-report 4. other 5. abstract 6. brief-report 7. letter 8. editorial 9. correction 10. book-review [ 25 others ] | 2001933 \| ( \| 72.8% \| ); 194578 \| ( \| 7.1% \| ); 126424 \| ( \| 4.6% \| ); 90582 \| ( \| 3.3% \| ); 73672 \| ( \| 2.7% \| ); 45631 \| ( \| 1.7% \| ); 42804 \| ( \| 1.6% \| ); 40855 \| ( \| 1.5% \| ); 36333 \| ( \| 1.3% \| ); 35682 \| ( \| 1.3% \| ); 62926 \| ( \| 2.3% \| ) |  | 2751420 (100%) | 0 (0%) |
| 26 | Subject [character] | 1. Article 2. Research Article 3. Original Article 4. Research 5. Articles 6. Case Report 7. Review 8. Original Research 9. Review Article 10. Research Paper [ 288316 others ] | 398622 \| ( \| 14.5% \| ); 321338 \| ( \| 11.7% \| ); 152190 \| ( \| 5.5% \| ); 119539 \| ( \| 4.3% \| ); 105833 \| ( \| 3.8% \| ); 100880 \| ( \| 3.7% \| ); 84650 \| ( \| 3.1% \| ); 52268 \| ( \| 1.9% \| ); 43353 \| ( \| 1.6% \| ); 37929 \| ( \| 1.4% \| ); 1334793 \| ( \| 48.5% \| ) |  | 2751395 (100%) | 25 (0%) |
| 27 | Number of authors [numeric] | Mean (sd) : 6 (27.8) min < med < max: 0 < 5 < 3573 IQR (CV) : 4 (4.6) | 653 distinct values |  | 2751420 (100%) | 0 (0%) |
| 28 | Number of affiliations [numeric] | Mean (sd) : 3.1 (5) min < med < max: 0 < 2 < 2858 IQR (CV) : 3 (1.6) | 289 distinct values |  | 2751420 (100%) | 0 (0%) |
| 29 | Number of references [numeric] | Mean (sd) : 36.9 (34.6) min < med < max: 0 < 32 < 3112 IQR (CV) : 35 (0.9) | 756 distinct values |  | 2751420 (100%) | 0 (0%) |
| 30 | Number of figures (in body) [numeric] | Mean (sd) : 2.4 (3.4) min < med < max: 0 < 1 < 404 IQR (CV) : 4 (1.4) | 164 distinct values |  | 2751420 (100%) | 0 (0%) |
| 31 | Number of figures (as floats) [numeric] | Mean (sd) : 0.9 (2.4) min < med < max: 0 < 0 < 186 IQR (CV) : 0 (2.7) | 85 distinct values |  | 2751420 (100%) | 0 (0%) |
| 32 | Number of tables (in body) [numeric] | Mean (sd) : 1.2 (1.9) min < med < max: 0 < 0 < 513 IQR (CV) : 2 (1.6) | 103 distinct values |  | 2751420 (100%) | 0 (0%) |
| 33 | Number of tables (as floats) [numeric] | Mean (sd) : 0.4 (1.2) min < med < max: 0 < 0 < 106 IQR (CV) : 0 (3.1) | 45 distinct values |  | 2751420 (100%) | 0 (0%) |
| 34 | Has supplement [logical] | 1. FALSE 2. TRUE | 1856638 \| ( \| 67.5% \| ); 894782 \| ( \| 32.5% \| ) |  | 2751420 (100%) | 0 (0%) |
| 35 | Research article (OCC) [logical] | 1. FALSE 2. TRUE | 284850 \| ( \| 11.1% \| ); 2285193 \| ( \| 88.9% \| ) |  | 2570043 (93.41%) | 181377 (6.59%) |
| 36 | Citation count (OCC) [numeric] | Mean (sd) : 16.6 (61.8) min < med < max: 1 < 6 < 15552 IQR (CV) : 13 (3.7) | 1453 distinct values |  | 1881234 (68.37%) | 870186 (31.63%) |
| 37 | Journal Impact Factor (WOS) [numeric] | Mean (sd) : 4.1 (3.5) min < med < max: 0 < 3 < 70.7 IQR (CV) : 2 (0.8) | 3219 distinct values |  | 1842322 (66.96%) | 909098 (33.04%) |
| 38 | Eigenfactor (WOS) [numeric] | Mean (sd) : 0.3 (0.6) min < med < max: 0 < 0 < 1.7 IQR (CV) : 0.1 (1.8) | 2110 distinct values |  | 1842524 (66.97%) | 908896 (33.03%) |
| 39 | Field of science (SciTech) [character] | 1. MEDICINE 2. HEALTH 3. BIOLOGY 4. INF DIS 5. BRAIN 6. CHEMISTRY 7. ENGNG 8. SOC SCI 9. COMP SCI 10. PHYS/MATH [ 2 others ] | 1175753 \| ( \| 49.9% \| ); 299299 \| ( \| 12.7% \| ); 261326 \| ( \| 11.1% \| ); 233527 \| ( \| 9.9% \| ); 194256 \| ( \| 8.2% \| ); 137313 \| ( \| 5.8% \| ); 22869 \| ( \| 1.0% \| ); 16499 \| ( \| 0.7% \| ); 11172 \| ( \| 0.5% \| ); 2544 \| ( \| 0.1% \| ); 531 \| ( \| 0.0% \| ) |  | 2355089 (85.6%) | 396331 (14.4%) |
| 40 | Uncorrupted XML file [logical] | 1. TRUE | 2751420 \| ( \| 100.0% \| ) |  | 2751420 (100%) | 0 (0%) |
