## Supplemental Table 4 for "Assessment of transparency indicators across the biomedical literature: how open is open?": s4-table_indicator-data-desc.html

Data Frame Summary


### Data Frame Summary

#### PMCOA

**Dimensions**: 2751420 x 120
  
**Duplicates**: 0
  

| **No** | **Variable** | **Stats / Values** | **Freqs (% of Valid)** | **Graph** | **Valid** | **Missing** |
| --- | --- | --- | --- | --- | --- | --- |
| 1 | PMCID [character] | 1. 100320 2. 100321 3. 100322 4. 100323 5. 100324 6. 100325 7. 100326 8. 100327 9. 100357 10. 100780 [ 2751410 others ] | 1 \| ( \| 0.0% \| ); 1 \| ( \| 0.0% \| ); 1 \| ( \| 0.0% \| ); 1 \| ( \| 0.0% \| ); 1 \| ( \| 0.0% \| ); 1 \| ( \| 0.0% \| ); 1 \| ( \| 0.0% \| ); 1 \| ( \| 0.0% \| ); 1 \| ( \| 0.0% \| ); 1 \| ( \| 0.0% \| ); 2751410 \| ( \| 100.0% \| ) |  | 2751420 (100%) | 0 (0%) |
| 2 | is\_data\_pred [logical] | 1. FALSE 2. TRUE | 2507637 \| ( \| 91.1% \| ); 243783 \| ( \| 8.9% \| ) |  | 2751420 (100%) | 0 (0%) |
| 3 | data\_text [character] | 1. (Empty string) 2. s1 table (xlsx 3. s1 file (xlsx 4. xls) click here for addit 5. xlsx) click here for addi 6. crystallographic data in 7. file s1 table (xlsx 8. xlsx) click here for addi 9. s1 file (xls 10. s1 table (xls [ 216810 others ] | 2511514 \| ( \| 91.3% \| ); 2168 \| ( \| 0.1% \| ); 1603 \| ( \| 0.1% \| ); 1373 \| ( \| 0.0% \| ); 1217 \| ( \| 0.0% \| ); 1183 \| ( \| 0.0% \| ); 1090 \| ( \| 0.0% \| ); 823 \| ( \| 0.0% \| ); 446 \| ( \| 0.0% \| ); 392 \| ( \| 0.0% \| ); 229611 \| ( \| 8.3% \| ) |  | 2751420 (100%) | 0 (0%) |
| 4 | is\_relevant\_data [logical] | 1. FALSE 2. TRUE | 304309 \| ( \| 11.1% \| ); 2447111 \| ( \| 88.9% \| ) |  | 2751420 (100%) | 0 (0%) |
| 5 | com\_specific\_db [character] | 1. the experimental structur 2. the suggested wording for 3. gene expression data pres 4. normalized and non-normal 5. sequence reads are availa 6. the icelandic population 7. a detailed description of 8. acc=gse34465 ) and is acc 9. accession numbers the ncb 10. according to miame requir [ 130791 others ] | 9 \| ( \| 0.0% \| ); 6 \| ( \| 0.0% \| ); 3 \| ( \| 0.0% \| ); 3 \| ( \| 0.0% \| ); 3 \| ( \| 0.0% \| ); 3 \| ( \| 0.0% \| ); 2 \| ( \| 0.0% \| ); 2 \| ( \| 0.0% \| ); 2 \| ( \| 0.0% \| ); 2 \| ( \| 0.0% \| ); 130917 \| ( \| 100.0% \| ) |  | 130952 (4.76%) | 2620468 (95.24%) |
| 6 | com\_general\_db [character] | 1. anonymized trial data wil 2. more information is avail 3. a data set of de-identifi 4. data file is available fr 5. data availability data ar 6. supporting data are also 7. all data are also availab 8. all supporting data and m 9. diffraction image data se 10. anonymized trial data wil [ 29083 others ] | 25 \| ( \| 0.1% \| ); 14 \| ( \| 0.0% \| ); 11 \| ( \| 0.0% \| ); 11 \| ( \| 0.0% \| ); 6 \| ( \| 0.0% \| ); 5 \| ( \| 0.0% \| ); 4 \| ( \| 0.0% \| ); 4 \| ( \| 0.0% \| ); 4 \| ( \| 0.0% \| ); 3 \| ( \| 0.0% \| ); 29152 \| ( \| 99.7% \| ) |  | 29239 (1.06%) | 2722181 (98.94%) |
| 7 | com\_github\_data [character] | 1. we used a custom script ( 2. a curated version of this 3. availability of data and 4. data availability opensim 5. reuse also applies to dat 6. the clean fastq files obt 7. the individual alignments 8. the specific scripts used 9. (14 answer options e g 'a 10. 0989141 anti-gapdh cell s [ 8615 others ] | 3 \| ( \| 0.0% \| ); 2 \| ( \| 0.0% \| ); 2 \| ( \| 0.0% \| ); 2 \| ( \| 0.0% \| ); 2 \| ( \| 0.0% \| ); 2 \| ( \| 0.0% \| ); 2 \| ( \| 0.0% \| ); 2 \| ( \| 0.0% \| ); 1 \| ( \| 0.0% \| ); 1 \| ( \| 0.0% \| ); 8615 \| ( \| 99.8% \| ) |  | 8634 (0.31%) | 2742786 (99.69%) |
| 8 | dataset [character] | 1. data s1. 2. supplementary dataset 1 p 3. supporting data s1. 4. supporting information ad 5. supplementary dataset 1 p 6. supplementary dataset 1. 7. data s1 . 8. supplementary information 9. data s1.; data s2. 10. data set s1 click here fo [ 35098 others ] | 260 \| ( \| 0.7% \| ); 199 \| ( \| 0.5% \| ); 49 \| ( \| 0.1% \| ); 48 \| ( \| 0.1% \| ); 37 \| ( \| 0.1% \| ); 31 \| ( \| 0.1% \| ); 25 \| ( \| 0.1% \| ); 23 \| ( \| 0.1% \| ); 21 \| ( \| 0.1% \| ); 20 \| ( \| 0.1% \| ); 35511 \| ( \| 98.0% \| ) |  | 36224 (1.32%) | 2715196 (98.68%) |
| 9 | com\_file\_formats [character] | 1. crystallographic data in 2. crystallographic data cen 3. crystallographic data cen 4. raw data (xlsx 5. crystallographic data ( c 6. crystallographic data (ci 7. crystallographic data cen 8. raw data (xls 9. all data csv 10. raw data in csv [ 2109 others ] | 1334 \| ( \| 28.7% \| ); 447 \| ( \| 9.6% \| ); 136 \| ( \| 2.9% \| ); 134 \| ( \| 2.9% \| ); 81 \| ( \| 1.7% \| ); 44 \| ( \| 0.9% \| ); 32 \| ( \| 0.7% \| ); 22 \| ( \| 0.5% \| ); 16 \| ( \| 0.3% \| ); 14 \| ( \| 0.3% \| ); 2385 \| ( \| 51.3% \| ) |  | 4645 (0.17%) | 2746775 (99.83%) |
| 10 | com\_supplemental\_data [character] | 1. s1 table (xlsx 2. xlsx) click here for addi 3. xls) click here for addit 4. file s1 table (xlsx 5. s1 file (xlsx 6. xlsx) click here for addi 7. xlsx) click here for addi 8. s1 table (xls 9. xls) click here for addit 10. s1 file (xls [ 31263 others ] | 2963 \| ( \| 4.4% \| ); 2908 \| ( \| 4.3% \| ); 2281 \| ( \| 3.4% \| ); 2160 \| ( \| 3.2% \| ); 1719 \| ( \| 2.5% \| ); 1465 \| ( \| 2.2% \| ); 1315 \| ( \| 2.0% \| ); 539 \| ( \| 0.8% \| ); 498 \| ( \| 0.7% \| ); 476 \| ( \| 0.7% \| ); 51105 \| ( \| 75.8% \| ) |  | 67429 (2.45%) | 2683991 (97.55%) |
| 11 | com\_data\_availibility [character] | 1. data availability data ar 2. data accessibility data a 3. data sharing statement: e 4. data availability all dat 5. data availability all dat 6. data availability data ar 7. data availability data ar 8. data availability all rel 9. data availability all rel 10. data accessibility data a [ 38463 others ] | 222 \| ( \| 0.5% \| ); 199 \| ( \| 0.5% \| ); 191 \| ( \| 0.4% \| ); 128 \| ( \| 0.3% \| ); 118 \| ( \| 0.3% \| ); 114 \| ( \| 0.3% \| ); 109 \| ( \| 0.2% \| ); 99 \| ( \| 0.2% \| ); 86 \| ( \| 0.2% \| ); 80 \| ( \| 0.2% \| ); 42575 \| ( \| 96.9% \| ) |  | 43921 (1.6%) | 2707499 (98.4%) |
| 12 | is\_code\_pred [logical] | 1. FALSE 2. TRUE | 2718015 \| ( \| 98.8% \| ); 33405 \| ( \| 1.2% \| ) |  | 2751420 (100%) | 0 (0%) |
| 13 | code\_text [character] | 1. (Empty string) 2. these routines are availa 3. the source code for the a 4. all macros and plugins ar 5. all r scripts used for th 6. analysis scripts are avai 7. the source code is availa 8. we used a custom script ( 9. a curated version of this 10. a current summary is avai [ 33312 others ] | 2718015 \| ( \| 98.8% \| ); 8 \| ( \| 0.0% \| ); 4 \| ( \| 0.0% \| ); 3 \| ( \| 0.0% \| ); 3 \| ( \| 0.0% \| ); 3 \| ( \| 0.0% \| ); 3 \| ( \| 0.0% \| ); 3 \| ( \| 0.0% \| ); 2 \| ( \| 0.0% \| ); 2 \| ( \| 0.0% \| ); 33374 \| ( \| 1.2% \| ) |  | 2751420 (100%) | 0 (0%) |
| 14 | is\_relevant\_code [logical] | 1. FALSE 2. TRUE | 1208523 \| ( \| 43.9% \| ); 1542897 \| ( \| 56.1% \| ) |  | 2751420 (100%) | 0 (0%) |
| 15 | com\_code [character] | 1. these routines are availa 2. the source code for the a 3. all macros and plugins ar 4. all r scripts used for th 5. analysis scripts are avai 6. the source code is availa 7. we used a custom script ( 8. a curated version of this 9. a current summary is avai 10. additional information ab [ 31077 others ] | 8 \| ( \| 0.0% \| ); 4 \| ( \| 0.0% \| ); 3 \| ( \| 0.0% \| ); 3 \| ( \| 0.0% \| ); 3 \| ( \| 0.0% \| ); 3 \| ( \| 0.0% \| ); 3 \| ( \| 0.0% \| ); 2 \| ( \| 0.0% \| ); 2 \| ( \| 0.0% \| ); 2 \| ( \| 0.0% \| ); 31139 \| ( \| 99.9% \| ) |  | 31172 (1.13%) | 2720248 (98.87%) |
| 16 | com\_suppl\_code [character] | 1. appendix a supplementary 2. availability of data and 3. data accessibility all da 4. data availability the fol 5. data availability the fol 6. in addition the configura 7. the r code is available i 8. the r code used for this 9. } more information on the 10. #circrnas / # bona fide ( [ 6278 others ] | 2 \| ( \| 0.0% \| ); 2 \| ( \| 0.0% \| ); 2 \| ( \| 0.0% \| ); 2 \| ( \| 0.0% \| ); 2 \| ( \| 0.0% \| ); 2 \| ( \| 0.0% \| ); 2 \| ( \| 0.0% \| ); 2 \| ( \| 0.0% \| ); 1 \| ( \| 0.0% \| ); 1 \| ( \| 0.0% \| ); 6278 \| ( \| 99.7% \| ) |  | 6296 (0.23%) | 2745124 (99.77%) |
| 17 | is\_coi\_pred [logical] | 1. FALSE 2. TRUE | 864513 \| ( \| 31.4% \| ); 1886907 \| ( \| 68.6% \| ) |  | 2751420 (100%) | 0 (0%) |
| 18 | coi\_text [character] | 1. (Empty string) 2. Competing interests The a 3. Competing Interests: The 4. Conflicts of Interest The 5. Conflict of Interest Stat 6. Competing interests: The 7. The authors have declared 8. Conflict of interest stat 9. Conflicts of interest The 10. Conflict of Interest: Non [ 354945 others ] | 864516 \| ( \| 31.4% \| ); 197246 \| ( \| 7.2% \| ); 185363 \| ( \| 6.7% \| ); 98470 \| ( \| 3.6% \| ); 63793 \| ( \| 2.3% \| ); 37739 \| ( \| 1.4% \| ); 34756 \| ( \| 1.3% \| ); 33324 \| ( \| 1.2% \| ); 30540 \| ( \| 1.1% \| ); 29042 \| ( \| 1.1% \| ); 1176631 \| ( \| 42.8% \| ) |  | 2751420 (100%) | 0 (0%) |
| 19 | is\_coi\_pmc\_fn [logical] | 1. FALSE 2. TRUE | 2134903 \| ( \| 77.6% \| ); 616517 \| ( \| 22.4% \| ) |  | 2751420 (100%) | 0 (0%) |
| 20 | is\_coi\_pmc\_title [logical] | 1. FALSE 2. TRUE | 962868 \| ( \| 45.1% \| ); 1172035 \| ( \| 54.9% \| ) |  | 2134903 (77.59%) | 616517 (22.41%) |
| 21 | is\_relevant\_coi [logical] | 1. FALSE 2. TRUE | 647854 \| ( \| 25.2% \| ); 1918178 \| ( \| 74.8% \| ) |  | 2566032 (93.26%) | 185388 (6.74%) |
| 22 | is\_relevant\_coi\_hi [logical] | 1. FALSE 2. TRUE | 824647 \| ( \| 32.1% \| ); 1741385 \| ( \| 67.9% \| ) |  | 2566032 (93.26%) | 185388 (6.74%) |
| 23 | is\_relevant\_coi\_lo [logical] | 1. FALSE 2. TRUE | 670684 \| ( \| 76.8% \| ); 202117 \| ( \| 23.2% \| ) |  | 872801 (31.72%) | 1878619 (68.28%) |
| 24 | is\_explicit\_coi [logical] | 1. FALSE 2. TRUE | 8288 \| ( \| 8.4% \| ); 90067 \| ( \| 91.6% \| ) |  | 98355 (3.57%) | 2653065 (96.43%) |
| 25 | coi\_1 [logical] | 1. FALSE 2. TRUE | 274682 \| ( \| 87.2% \| ); 40332 \| ( \| 12.8% \| ) |  | 315014 (11.45%) | 2436406 (88.55%) |
| 26 | coi\_2 [logical] | 1. FALSE 2. TRUE | 242605 \| ( \| 77.0% \| ); 72409 \| ( \| 23.0% \| ) |  | 315014 (11.45%) | 2436406 (88.55%) |
| 27 | coi\_disclosure\_1 [logical] | 1. FALSE 2. TRUE | 305436 \| ( \| 97.0% \| ); 9578 \| ( \| 3.0% \| ) |  | 315014 (11.45%) | 2436406 (88.55%) |
| 28 | commercial\_1 [logical] | 1. FALSE 2. TRUE | 280540 \| ( \| 89.1% \| ); 34474 \| ( \| 10.9% \| ) |  | 315014 (11.45%) | 2436406 (88.55%) |
| 29 | benefit\_1 [logical] | 1. FALSE 2. TRUE | 314724 \| ( \| 99.9% \| ); 290 \| ( \| 0.1% \| ) |  | 315014 (11.45%) | 2436406 (88.55%) |
| 30 | consultant\_1 [logical] | 1. FALSE 2. TRUE | 313900 \| ( \| 99.7% \| ); 1114 \| ( \| 0.4% \| ) |  | 315014 (11.45%) | 2436406 (88.55%) |
| 31 | grants\_1 [logical] | 1. FALSE 2. TRUE | 314484 \| ( \| 99.8% \| ); 530 \| ( \| 0.2% \| ) |  | 315014 (11.45%) | 2436406 (88.55%) |
| 32 | brief\_1 [logical] | 1. FALSE | 315014 \| ( \| 100.0% \| ) |  | 315014 (11.45%) | 2436406 (88.55%) |
| 33 | fees\_1 [logical] | 1. FALSE 2. TRUE | 70185 \| ( \| 98.4% \| ); 1157 \| ( \| 1.6% \| ) |  | 71342 (2.59%) | 2680078 (97.41%) |
| 34 | consults\_1 [logical] | 1. FALSE 2. TRUE | 68020 \| ( \| 95.3% \| ); 3322 \| ( \| 4.7% \| ) |  | 71342 (2.59%) | 2680078 (97.41%) |
| 35 | connect\_1 [logical] | 1. FALSE 2. TRUE | 70830 \| ( \| 99.3% \| ); 512 \| ( \| 0.7% \| ) |  | 71342 (2.59%) | 2680078 (97.41%) |
| 36 | connect\_2 [logical] | 1. FALSE 2. TRUE | 71185 \| ( \| 99.8% \| ); 157 \| ( \| 0.2% \| ) |  | 71342 (2.59%) | 2680078 (97.41%) |
| 37 | commercial\_ack\_1 [logical] | 1. FALSE 2. TRUE | 71210 \| ( \| 99.8% \| ); 132 \| ( \| 0.2% \| ) |  | 71342 (2.59%) | 2680078 (97.41%) |
| 38 | rights\_1 [logical] | 1. FALSE 2. TRUE | 71259 \| ( \| 99.9% \| ); 83 \| ( \| 0.1% \| ) |  | 71342 (2.59%) | 2680078 (97.41%) |
| 39 | founder\_1 [logical] | 1. FALSE 2. TRUE | 71028 \| ( \| 99.6% \| ); 314 \| ( \| 0.4% \| ) |  | 71342 (2.59%) | 2680078 (97.41%) |
| 40 | advisor\_1 [logical] | 1. FALSE 2. TRUE | 71138 \| ( \| 99.7% \| ); 204 \| ( \| 0.3% \| ) |  | 71342 (2.59%) | 2680078 (97.41%) |
| 41 | paid\_1 [logical] | 1. FALSE 2. TRUE | 70745 \| ( \| 99.2% \| ); 597 \| ( \| 0.8% \| ) |  | 71342 (2.59%) | 2680078 (97.41%) |
| 42 | board\_1 [logical] | 1. FALSE 2. TRUE | 70928 \| ( \| 99.4% \| ); 414 \| ( \| 0.6% \| ) |  | 71342 (2.59%) | 2680078 (97.41%) |
| 43 | no\_coi\_1 [logical] | 1. FALSE 2. TRUE | 70926 \| ( \| 99.4% \| ); 416 \| ( \| 0.6% \| ) |  | 71342 (2.59%) | 2680078 (97.41%) |
| 44 | no\_funder\_role\_1 [logical] | 1. FALSE 2. TRUE | 68710 \| ( \| 96.3% \| ); 2632 \| ( \| 3.7% \| ) |  | 71342 (2.59%) | 2680078 (97.41%) |
| 45 | is\_fund\_pred [logical] | 1. FALSE 2. TRUE | 893398 \| ( \| 32.5% \| ); 1858022 \| ( \| 67.5% \| ) |  | 2751420 (100%) | 0 (0%) |
| 46 | fund\_text [character] | 1. (Empty string) 2. Financial support and spo 3. Source of Support: Nil 4. Funding This research rec 5. The authors received no s 6. The authors have no suppo 7. FUNDING 8. The funders had no role i 9. Source of Support: Nil. 10. Funding None. [ 1593777 others ] | 893399 \| ( \| 32.5% \| ); 32330 \| ( \| 1.2% \| ); 27721 \| ( \| 1.0% \| ); 12264 \| ( \| 0.4% \| ); 10570 \| ( \| 0.4% \| ); 7879 \| ( \| 0.3% \| ); 7292 \| ( \| 0.3% \| ); 6681 \| ( \| 0.2% \| ); 6267 \| ( \| 0.2% \| ); 5894 \| ( \| 0.2% \| ); 1741123 \| ( \| 63.3% \| ) |  | 2751420 (100%) | 0 (0%) |
| 47 | fund\_pmc\_institute [character] | 1. (Empty string) 2. National Natural Science 3. National Institutes of He 4. National Research Foundat 5. Deutsche Forschungsgemein 6. National Natural Science 7. Japan Society for the Pro 8. Wellcome Trust 9. National Institute for He 10. National Health and Medic [ 155352 others ] | 2495885 \| ( \| 90.7% \| ); 4266 \| ( \| 0.2% \| ); 1890 \| ( \| 0.1% \| ); 1820 \| ( \| 0.1% \| ); 1325 \| ( \| 0.0% \| ); 1253 \| ( \| 0.0% \| ); 1075 \| ( \| 0.0% \| ); 971 \| ( \| 0.0% \| ); 911 \| ( \| 0.0% \| ); 874 \| ( \| 0.0% \| ); 241150 \| ( \| 8.8% \| ) |  | 2751420 (100%) | 0 (0%) |
| 48 | fund\_pmc\_source [character] | 1. (Empty string) 2. National Natural Science 3. National Institutes of He 4. http://dx.doi.org/10.1303 5. National Institutes of He 6. National Natural Science 7. National Institutes of He 8. Deutsche Forschungsgemein 9. http://dx.doi.org/10.1303 10. https://doi.org/10.13039/ [ 265892 others ] | 2319255 \| ( \| 84.3% \| ); 4655 \| ( \| 0.2% \| ); 4549 \| ( \| 0.2% \| ); 3305 \| ( \| 0.1% \| ); 2582 \| ( \| 0.1% \| ); 2442 \| ( \| 0.1% \| ); 1616 \| ( \| 0.1% \| ); 1438 \| ( \| 0.1% \| ); 1401 \| ( \| 0.1% \| ); 1252 \| ( \| 0.0% \| ); 408925 \| ( \| 14.9% \| ) |  | 2751420 (100%) | 0 (0%) |
| 49 | fund\_pmc\_anysource [character] | 1. (Empty string) 2. National Institutes of He 3. Wellcome Trust 4. NIH 5. National Natural Science 6. Austrian Science Fund 7. Medical Research Council 8. Austrian Science Fund (FW 9. National Science Foundati 10. Biotechnology and Biologi [ 33530 others ] | 2707125 \| ( \| 98.4% \| ); 637 \| ( \| 0.0% \| ); 588 \| ( \| 0.0% \| ); 518 \| ( \| 0.0% \| ); 464 \| ( \| 0.0% \| ); 209 \| ( \| 0.0% \| ); 198 \| ( \| 0.0% \| ); 147 \| ( \| 0.0% \| ); 139 \| ( \| 0.0% \| ); 128 \| ( \| 0.0% \| ); 41267 \| ( \| 1.5% \| ) |  | 2751420 (100%) | 0 (0%) |
| 50 | is\_fund\_pmc\_group [logical] | 1. FALSE 2. TRUE | 2163660 \| ( \| 78.6% \| ); 587760 \| ( \| 21.4% \| ) |  | 2751420 (100%) | 0 (0%) |
| 51 | is\_fund\_pmc\_title [logical] | 1. FALSE 2. TRUE | 1890612 \| ( \| 75.3% \| ); 619674 \| ( \| 24.7% \| ) |  | 2510286 (91.24%) | 241134 (8.76%) |
| 52 | is\_fund\_pmc\_anysource [logical] | 1. FALSE 2. TRUE | 1846317 \| ( \| 97.7% \| ); 44295 \| ( \| 2.3% \| ) |  | 1890612 (68.71%) | 860808 (31.29%) |
| 53 | is\_relevant\_fund [logical] | 1. FALSE 2. TRUE | 540700 \| ( \| 29.4% \| ); 1297601 \| ( \| 70.6% \| ) |  | 1838301 (66.81%) | 913119 (33.19%) |
| 54 | is\_explicit\_fund [logical] | 1. FALSE 2. TRUE | 81728 \| ( \| 5.1% \| ); 1535160 \| ( \| 95.0% \| ) |  | 1616888 (58.77%) | 1134532 (41.23%) |
| 55 | support\_1 [logical] | 1. FALSE 2. TRUE | 690030 \| ( \| 51.1% \| ); 659882 \| ( \| 48.9% \| ) |  | 1349912 (49.06%) | 1401508 (50.94%) |
| 56 | support\_3 [logical] | 1. FALSE 2. TRUE | 689504 \| ( \| 51.1% \| ); 660408 \| ( \| 48.9% \| ) |  | 1349912 (49.06%) | 1401508 (50.94%) |
| 57 | support\_4 [logical] | 1. FALSE 2. TRUE | 870632 \| ( \| 64.5% \| ); 479280 \| ( \| 35.5% \| ) |  | 1349912 (49.06%) | 1401508 (50.94%) |
| 58 | support\_5 [logical] | 1. FALSE 2. TRUE | 781086 \| ( \| 57.9% \| ); 568826 \| ( \| 42.1% \| ) |  | 1349912 (49.06%) | 1401508 (50.94%) |
| 59 | support\_6 [logical] | 1. FALSE 2. TRUE | 1310742 \| ( \| 97.1% \| ); 39170 \| ( \| 2.9% \| ) |  | 1349912 (49.06%) | 1401508 (50.94%) |
| 60 | support\_7 [logical] | 1. FALSE 2. TRUE | 1344182 \| ( \| 99.6% \| ); 5730 \| ( \| 0.4% \| ) |  | 1349912 (49.06%) | 1401508 (50.94%) |
| 61 | support\_8 [logical] | 1. FALSE 2. TRUE | 1346895 \| ( \| 99.8% \| ); 3017 \| ( \| 0.2% \| ) |  | 1349912 (49.06%) | 1401508 (50.94%) |
| 62 | support\_9 [logical] | 1. FALSE 2. TRUE | 1343577 \| ( \| 99.5% \| ); 6335 \| ( \| 0.5% \| ) |  | 1349912 (49.06%) | 1401508 (50.94%) |
| 63 | support\_10 [logical] | 1. FALSE 2. TRUE | 1329402 \| ( \| 98.5% \| ); 20510 \| ( \| 1.5% \| ) |  | 1349912 (49.06%) | 1401508 (50.94%) |
| 64 | developed\_1 [logical] | 1. FALSE 2. TRUE | 1349372 \| ( \| 100.0% \| ); 540 \| ( \| 0.0% \| ) |  | 1349912 (49.06%) | 1401508 (50.94%) |
| 65 | received\_1 [logical] | 1. FALSE 2. TRUE | 1333990 \| ( \| 98.8% \| ); 15922 \| ( \| 1.2% \| ) |  | 1349912 (49.06%) | 1401508 (50.94%) |
| 66 | received\_2 [logical] | 1. FALSE 2. TRUE | 1329882 \| ( \| 98.5% \| ); 20030 \| ( \| 1.5% \| ) |  | 1349912 (49.06%) | 1401508 (50.94%) |
| 67 | recipient\_1 [logical] | 1. FALSE 2. TRUE | 1337017 \| ( \| 99.0% \| ); 12895 \| ( \| 1.0% \| ) |  | 1349912 (49.06%) | 1401508 (50.94%) |
| 68 | authors\_1 [logical] | 1. FALSE 2. TRUE | 1281051 \| ( \| 94.9% \| ); 68861 \| ( \| 5.1% \| ) |  | 1349912 (49.06%) | 1401508 (50.94%) |
| 69 | authors\_2 [logical] | 1. FALSE 2. TRUE | 1342588 \| ( \| 99.5% \| ); 7324 \| ( \| 0.5% \| ) |  | 1349912 (49.06%) | 1401508 (50.94%) |
| 70 | thank\_1 [logical] | 1. FALSE 2. TRUE | 1329217 \| ( \| 98.5% \| ); 20695 \| ( \| 1.5% \| ) |  | 1349912 (49.06%) | 1401508 (50.94%) |
| 71 | thank\_2 [logical] | 1. FALSE 2. TRUE | 1296615 \| ( \| 96.0% \| ); 53297 \| ( \| 4.0% \| ) |  | 1349912 (49.06%) | 1401508 (50.94%) |
| 72 | fund\_1 [logical] | 1. FALSE 2. TRUE | 1336035 \| ( \| 99.0% \| ); 13877 \| ( \| 1.0% \| ) |  | 1349912 (49.06%) | 1401508 (50.94%) |
| 73 | fund\_2 [logical] | 1. FALSE 2. TRUE | 1296140 \| ( \| 96.0% \| ); 53772 \| ( \| 4.0% \| ) |  | 1349912 (49.06%) | 1401508 (50.94%) |
| 74 | fund\_3 [logical] | 1. FALSE 2. TRUE | 1342013 \| ( \| 99.4% \| ); 7899 \| ( \| 0.6% \| ) |  | 1349912 (49.06%) | 1401508 (50.94%) |
| 75 | supported\_1 [logical] | 1. FALSE 2. TRUE | 1337413 \| ( \| 99.1% \| ); 12499 \| ( \| 0.9% \| ) |  | 1349912 (49.06%) | 1401508 (50.94%) |
| 76 | financial\_1 [logical] | 1. FALSE 2. TRUE | 1331445 \| ( \| 98.6% \| ); 18467 \| ( \| 1.4% \| ) |  | 1349912 (49.06%) | 1401508 (50.94%) |
| 77 | financial\_2 [logical] | 1. FALSE 2. TRUE | 1349910 \| ( \| 100.0% \| ); 2 \| ( \| 0.0% \| ) |  | 1349912 (49.06%) | 1401508 (50.94%) |
| 78 | financial\_3 [logical] | 1. FALSE 2. TRUE | 1341248 \| ( \| 99.4% \| ); 8664 \| ( \| 0.6% \| ) |  | 1349912 (49.06%) | 1401508 (50.94%) |
| 79 | grant\_1 [logical] | 1. FALSE 2. TRUE | 1340542 \| ( \| 99.3% \| ); 9370 \| ( \| 0.7% \| ) |  | 1349912 (49.06%) | 1401508 (50.94%) |
| 80 | french\_1 [logical] | 1. FALSE 2. TRUE | 1349891 \| ( \| 100.0% \| ); 21 \| ( \| 0.0% \| ) |  | 1349912 (49.06%) | 1401508 (50.94%) |
| 81 | common\_1 [logical] | 1. FALSE 2. TRUE | 1346418 \| ( \| 99.7% \| ); 3494 \| ( \| 0.3% \| ) |  | 1349912 (49.06%) | 1401508 (50.94%) |
| 82 | common\_2 [logical] | 1. FALSE 2. TRUE | 1344506 \| ( \| 99.6% \| ); 5406 \| ( \| 0.4% \| ) |  | 1349912 (49.06%) | 1401508 (50.94%) |
| 83 | common\_3 [logical] | 1. FALSE 2. TRUE | 1349911 \| ( \| 100.0% \| ); 1 \| ( \| 0.0% \| ) |  | 1349912 (49.06%) | 1401508 (50.94%) |
| 84 | common\_4 [logical] | 1. FALSE 2. TRUE | 1339680 \| ( \| 99.2% \| ); 10232 \| ( \| 0.8% \| ) |  | 1349912 (49.06%) | 1401508 (50.94%) |
| 85 | common\_5 [logical] | 1. FALSE 2. TRUE | 1348880 \| ( \| 99.9% \| ); 1032 \| ( \| 0.1% \| ) |  | 1349912 (49.06%) | 1401508 (50.94%) |
| 86 | acknow\_1 [logical] | 1. FALSE 2. TRUE | 1348688 \| ( \| 99.9% \| ); 1224 \| ( \| 0.1% \| ) |  | 1349912 (49.06%) | 1401508 (50.94%) |
| 87 | disclosure\_1 [logical] | 1. FALSE 2. TRUE | 1349701 \| ( \| 100.0% \| ); 211 \| ( \| 0.0% \| ) |  | 1349912 (49.06%) | 1401508 (50.94%) |
| 88 | disclosure\_2 [logical] | 1. FALSE 2. TRUE | 1340801 \| ( \| 99.3% \| ); 9111 \| ( \| 0.7% \| ) |  | 1349912 (49.06%) | 1401508 (50.94%) |
| 89 | fund\_ack [logical] | 1. FALSE 2. TRUE | 127540 \| ( \| 63.9% \| ); 71930 \| ( \| 36.1% \| ) |  | 199470 (7.25%) | 2551950 (92.75%) |
| 90 | project\_ack [logical] | 1. FALSE 2. TRUE | 197493 \| ( \| 99.0% \| ); 1977 \| ( \| 1.0% \| ) |  | 199470 (7.25%) | 2551950 (92.75%) |
| 91 | is\_register\_pred [logical] | 1. FALSE 2. TRUE | 2680391 \| ( \| 97.4% \| ); 71029 \| ( \| 2.6% \| ) |  | 2751420 (100%) | 0 (0%) |
| 92 | register\_text [character] | 1. (Empty string) 2. Trial registration Not ap 3. Trial registration: 4. Clinical Procedure: None 5. Trial registration Retros 6. TRIAL REGISTRATION NUMBER 7. Trial registration 8. TRIAL REGISTRATION NUMBER 9. Trial Registration: 10. Clinical trial registrati [ 69987 others ] | 2680391 \| ( \| 97.4% \| ); 93 \| ( \| 0.0% \| ); 36 \| ( \| 0.0% \| ); 32 \| ( \| 0.0% \| ); 28 \| ( \| 0.0% \| ); 23 \| ( \| 0.0% \| ); 22 \| ( \| 0.0% \| ); 19 \| ( \| 0.0% \| ); 19 \| ( \| 0.0% \| ); 11 \| ( \| 0.0% \| ); 70746 \| ( \| 2.6% \| ) |  | 2751420 (100%) | 0 (0%) |
| 93 | is\_research [logical] | 1. FALSE 2. TRUE | 653941 \| ( \| 23.8% \| ); 2097479 \| ( \| 76.2% \| ) |  | 2751420 (100%) | 0 (0%) |
| 94 | is\_review [logical] | 1. FALSE 2. TRUE | 2554902 \| ( \| 92.9% \| ); 196518 \| ( \| 7.1% \| ) |  | 2751420 (100%) | 0 (0%) |
| 95 | is\_reg\_pmc\_title [logical] | 1. FALSE 2. TRUE | 2713910 \| ( \| 98.6% \| ); 37510 \| ( \| 1.4% \| ) |  | 2751420 (100%) | 0 (0%) |
| 96 | is\_relevant\_reg [logical] | 1. FALSE 2. TRUE | 1878880 \| ( \| 81.9% \| ); 415117 \| ( \| 18.1% \| ) |  | 2293997 (83.38%) | 457423 (16.62%) |
| 97 | is\_method [logical] | 1. FALSE 2. TRUE | 29373 \| ( \| 7.8% \| ); 348234 \| ( \| 92.2% \| ) |  | 377607 (13.72%) | 2373813 (86.28%) |
| 98 | is\_NCT [logical] | 1. FALSE 2. TRUE | 324621 \| ( \| 93.2% \| ); 23613 \| ( \| 6.8% \| ) |  | 348234 (12.66%) | 2403186 (87.34%) |
| 99 | is\_explicit\_reg [logical] | 1. FALSE 2. TRUE | 2648 \| ( \| 3.7% \| ); 68381 \| ( \| 96.3% \| ) |  | 71029 (2.58%) | 2680391 (97.42%) |
| 100 | prospero\_1 [logical] | 1. FALSE 2. TRUE | 343073 \| ( \| 98.5% \| ); 5161 \| ( \| 1.5% \| ) |  | 348234 (12.66%) | 2403186 (87.34%) |
| 101 | registered\_1 [logical] | 1. FALSE 2. TRUE | 328288 \| ( \| 94.3% \| ); 19946 \| ( \| 5.7% \| ) |  | 348234 (12.66%) | 2403186 (87.34%) |
| 102 | registered\_2 [logical] | 1. FALSE 2. TRUE | 347428 \| ( \| 99.8% \| ); 806 \| ( \| 0.2% \| ) |  | 348234 (12.66%) | 2403186 (87.34%) |
| 103 | registered\_3 [logical] | 1. FALSE 2. TRUE | 347999 \| ( \| 99.9% \| ); 235 \| ( \| 0.1% \| ) |  | 348234 (12.66%) | 2403186 (87.34%) |
| 104 | registered\_4 [logical] | 1. FALSE 2. TRUE | 346949 \| ( \| 99.6% \| ); 1285 \| ( \| 0.4% \| ) |  | 348234 (12.66%) | 2403186 (87.34%) |
| 105 | registered\_5 [logical] | 1. FALSE 2. TRUE | 348198 \| ( \| 100.0% \| ); 36 \| ( \| 0.0% \| ) |  | 348234 (12.66%) | 2403186 (87.34%) |
| 106 | not\_registered\_1 [logical] | 1. FALSE 2. TRUE | 347674 \| ( \| 99.8% \| ); 560 \| ( \| 0.2% \| ) |  | 348234 (12.66%) | 2403186 (87.34%) |
| 107 | registration\_1 [logical] | 1. FALSE 2. TRUE | 347124 \| ( \| 99.7% \| ); 1110 \| ( \| 0.3% \| ) |  | 348234 (12.66%) | 2403186 (87.34%) |
| 108 | registration\_2 [logical] | 1. FALSE 2. TRUE | 346945 \| ( \| 99.6% \| ); 1289 \| ( \| 0.4% \| ) |  | 348234 (12.66%) | 2403186 (87.34%) |
| 109 | registration\_3 [logical] | 1. FALSE 2. TRUE | 347438 \| ( \| 99.8% \| ); 796 \| ( \| 0.2% \| ) |  | 348234 (12.66%) | 2403186 (87.34%) |
| 110 | registration\_4 [logical] | 1. FALSE 2. TRUE | 344116 \| ( \| 98.8% \| ); 4118 \| ( \| 1.2% \| ) |  | 348234 (12.66%) | 2403186 (87.34%) |
| 111 | registry\_1 [logical] | 1. FALSE 2. TRUE | 345296 \| ( \| 99.2% \| ); 2938 \| ( \| 0.8% \| ) |  | 348234 (12.66%) | 2403186 (87.34%) |
| 112 | reg\_title\_1 [logical] | 1. FALSE 2. TRUE | 345132 \| ( \| 99.1% \| ); 3102 \| ( \| 0.9% \| ) |  | 348234 (12.66%) | 2403186 (87.34%) |
| 113 | reg\_title\_2 [logical] | 1. FALSE 2. TRUE | 344743 \| ( \| 99.0% \| ); 3491 \| ( \| 1.0% \| ) |  | 348234 (12.66%) | 2403186 (87.34%) |
| 114 | reg\_title\_3 [logical] | 1. FALSE 2. TRUE | 346454 \| ( \| 99.5% \| ); 1780 \| ( \| 0.5% \| ) |  | 348234 (12.66%) | 2403186 (87.34%) |
| 115 | reg\_title\_4 [logical] | 1. FALSE 2. TRUE | 347518 \| ( \| 99.8% \| ); 716 \| ( \| 0.2% \| ) |  | 348234 (12.66%) | 2403186 (87.34%) |
| 116 | funded\_ct\_1 [logical] | 1. FALSE 2. TRUE | 348197 \| ( \| 100.0% \| ); 37 \| ( \| 0.0% \| ) |  | 348234 (12.66%) | 2403186 (87.34%) |
| 117 | ct\_2 [logical] | 1. FALSE 2. TRUE | 315592 \| ( \| 99.4% \| ); 1771 \| ( \| 0.6% \| ) |  | 317363 (11.53%) | 2434057 (88.47%) |
| 118 | ct\_3 [logical] | 1. FALSE 2. TRUE | 316674 \| ( \| 99.8% \| ); 689 \| ( \| 0.2% \| ) |  | 317363 (11.53%) | 2434057 (88.47%) |
| 119 | protocol\_1 [logical] | 1. FALSE 2. TRUE | 317063 \| ( \| 99.9% \| ); 300 \| ( \| 0.1% \| ) |  | 317363 (11.53%) | 2434057 (88.47%) |
| 120 | is\_success\_pmc [logical] | 1. TRUE | 2751420 \| ( \| 100.0% \| ) |  | 2751420 (100%) | 0 (0%) |
